# Phylogenomic analyses reveal that Panguiarchaeum is a clade of genome-reduced Asgard archaea within the Njordarchaeia

**DOI:** 10.1101/2025.02.13.637844

**Authors:** Wen-Cong Huang, Maraike Probst, Zheng-Shuang Hua, Lénárd L. Szánthó, Gergely J. Szöllősi, Thijs J. G. Ettema, Christian Rinke, Tom A. Williams, Anja Spang

## Abstract

The Asgard archaea are a diverse archaeal phylum important for our understanding of cellular evolution because they include the lineage that gave rise to eukaryotes. Recent phylogenomic work has focused on characterising the diversity of Asgard archaea in an effort to identify the closest extant relatives of eukaryotes. However, resolving archaeal phylogeny is challenging, and the positions of two recently-described lineages - Njordarchaeales and Panguiarchaeales - are uncertain, in ways that directly bear on hypotheses of early evolution. In initial phylogenetic analyses, these lineages branched either with Asgards or with the distantly-related Korarchaeota, and it has been suggested that their genomes may be affected by metagenomic contamination. Resolving this debate is important because these clades include genome-reduced lineages that may help inform our understanding of the evolution of symbiosis within Asgard archaea. Here, we performed phylogenetic analyses revealing that the Njordarchaeales and Pangiuarchaeales constitute the new class Njordarchaeia within Asgard archaea. We found no evidence of metagenomic contamination affecting phylogenetic analyses. Njordarchaeia exhibit hallmarks of adaptations to (hyper-)thermophilic lifestyles, including biased sequence compositions that can induce phylogenetic artifacts unless adequately modelled. *Panguiarchaeum* is metabolically distinct from its relatives, with reduced metabolic potential and various auxotrophies. Phylogenetic reconciliation recovers a complex common ancestor of Asgard archaea that encoded the Wood-Ljungdahl pathway. The subsequent loss of this pathway during the reductive evolution of *Panguiarchaeum* may have been associated with the switch to a symbiotic lifestyle based on H_2_-syntrophy. Thus, *Panguiarchaeum* may contain the first obligate symbionts within Asgard archaea.

## Introduction

Asgard archaea are a diverse phylum from which the archaeal ancestor of eukaryotes emerged (Spang et al. 2015; Zaremba-Niedzwiedzka et al. 2017; Liu et al. 2021; Zhang et al. 2025). The genomes of Asgard archaea (Spang et al. 2015; Zaremba-Niedzwiedzka et al. 2017; Liu et al. 2021; Wu et al. 2022; Valentin-Alvarado et al. 2024) encode eukaryotic signature proteins (ESPs) (Vosseberg et al. 2024), which are homologs of eukaryotic proteins that underlie cellular complexity. Experimental evidence suggests that at least some ESPs of Asgard archaea have functions similar to those in eukaryotes (Akıl and Robinson 2018; Akıl et al. 2020; Lu et al. 2020; Hatano et al. 2022; Hurtig et al. 2023). Genomic predictions and the cultivation of the first representatives of the Asgard archaea suggest that at least some members form symbiotic, (that is, syntrophic), interactions with other cells (Spang et al. 2019; Imachi et al. 2020). Indeed, eukaryotes are thought to have evolved from one such interaction (Martin and Müller 1998; Moreira and Lopez-Garcia 1998; Lane and Martin 2010; Guy et al. 2014; Koonin and Yutin 2014; Poole and Gribaldo 2014; López-García and Moreira 2015; Ettema 2016; Roger et al. 2017; Vosseberg et al. 2024). The phylogeny and genomic features of Asgard archaea and related groups therefore provide an important framework for understanding the origin of eukaryotic cells and the evolution of prokaryotic symbioses (Spang et al. 2015; Zaremba-Niedzwiedzka et al. 2017; Williams et al. 2020; Liu et al. 2021; Zhang et al. 2025). A major current challenge lies in resolving the evolutionary relationships among Asgard archaea and their relatives, including Thermoproteota (formerly Thaumarchaeota, Aigarchaeota, and Crenarchaeota) and Korarchaeota (Elkins et al. 2008). Early analyses supported a clade of “TACK” Archaea (Guy and Ettema 2011) (initially comprising the phyla Thaumarchaeota, Aigarchaeota, Crenarchaeota, and Koarchaeota now classified as class-level lineages within the Thermoproteota) as the sister lineage to the Asgard archaea (Spang et al. 2015; Zaremba-Niedzwiedzka et al. 2017; Baker et al. 2020). However, more recent phylogenetic analyses taking advantage of improved genomic sampling have instead placed the Korarchaeota at the base of a monophyletic Thermoproteota and Asgard clade (Tahon et al. 2023).

Two additional lineages have recently been described that appear to be part of a broader Asgard/Thermoproteota/Korarchaeota clade, although their phylogenetic affinities remain uncertain: the Njordarchaeales (Liu and Li 2022; Xie et al. 2022) and the Panguiarchaeales (Qu et al. 2023). Njordarchaeales have been recovered either as sister to Korarchaeota (Liu and Li 2022) - with whom they share a thermophilic lifestyle - or, intriguingly, within Asgard archaea, either as sister to the eukaryotic nuclear lineage (Xie et al. 2022), or an order within the Heimdallarchaeia (Eme et al. 2023; Valentin-Alvarado et al. 2024). Placement outside Asgard is observed when the hyperthermophilic Korarchaeota are included (Liu and Li 2022; Eme et al. 2023; Qu et al. 2023), while their inclusion within Asgard is more common when these taxa are excluded (Eme et al. 2023; Valentin-Alvarado et al. 2024), depending on the dataset and phylogenetic method. Similarly, the recently-reported Panguiarchaeales, composed of 10 metagenome-assembled genomes (MAGs) derived from terrestrial geothermal metagenomes, all belonging to the same species, were suggested to branch with Korarchaeota based on four distinct marker sets using site-homogeneous model (Qu et al. 2023). But conflicting placement as Asgard archaea was observed using 53 marker genes from GTDB (release 207v2) (Dombrowski et al. 2020; Rinke et al. 2021). Panguiarchaeales were proposed to represent anaerobic amino acid fermenters with a symbiotic lifestyle (Qu et al. 2023) because comparative genomic analyses indicated that they lack certain genes for archaeal lipid, purine and amino acid biosynthesis pathways reminiscent of DPANN archaea (Qu et al. 2023). However, the authors could not identify direct evidence for recent reductive genome evolution since key genes for DNA mismatch repair mechanisms and homologous recombination were present in the investigated MAGs, which did not significantly differ in size from other Korarchaeota with which they were proposed affiliated (Qu et al. 2023). Reduced metabolic capabilities are hallmarks of the DPANN archaea, which represent diverse phylum-level lineages composed of putative symbionts (Rinke et al. 2013; Castelle et al. 2018), but are otherwise uncommon among archaea. Thus, resolving the evolutionary relationships within the Asgard/Thermoproteota/Korarchaeota clade, and the positions of Njordarchaeales and Panguiarchaeales within them, is key to understanding the evolutionary trajectories of genome reduction and symbiotic associations in Archaea as well as for placing the origin of eukaryotic cells in the broader context of archaeal evolution. An additional challenge in this regard is the spectre of metagenomic contamination, with a recent study (Zhang et al. 2025) suggesting that at least some Njordarchaeales MAGs might be chimeras, containing a mixture of sequences derived from both Asgard archaea and Korarchaeota. If correct, this would provide an alternative explanation for the difficulty of placing Njordarchaeles in phylogenies, because combining marker genes with different evolutionary histories would undermine a core assumption of concatenated phylogenetic analyses.

Here, we carefully assembled an archaeal taxa set, inspected MAGs quality of key representatives of the Pangui- and Njordarchaeales and applied a variety of comparative genomics, phylogenomics, and network approaches to determine the phylogenetic placement of Njordarchaeales and Panguiarchaeales within the archaeal tree of life and to assess the genomic potential and proposed symbiotic lifestyle of the latter. Our analyses reveal that the previously described Njordarchaeales and Panguiarchaeales orders are order- or family-level lineages which together form the new class Njordarchaeia within the Asgard archaea rather than the Korarchaeota. We found no evidence for metagenomic contamination affecting marker gene sequences; instead, our analyses show that the difficulty in placing Njordarchaeia in the archaeal tree is due to pervasive site-wise and branch-wise compositional heterogeneity in the sequence data that are challenging to model adequately, particularly with simpler substitution models. While all Njordarchaeia encode a similar set of ESPs, *Panguiarchaeaceae* MAGs show indications of reductive genome evolution and auxotrophies, suggesting that they may be dependent on partner organisms or other community members to supplement their growth. Indeed, our co-occurrence analyses reveal a higher-than-expected frequency of association between *Panguiarchaeum* and members of the Thermoproteota. Overall, this study shows that MAG contamination is not the cause for conflicting phylogenetic signal observed for members of the Njordarchaeia and instead highlights the need to use phylogenetic models that provide an adequate fit to the data when inferring the archaeal phylogeny. It further provides the first example of a putative symbiotic lineage within the Asgard archaea previously assigned to Korarchaeota that has experienced gene loss rather than genome expansion.

## Results

### *Panguiarchaeum genomes* form a sister group to *Njordarchaeaceae* and branch within Asgard archaea

#### Assembling a set of vertically-evolving archaeal marker genes

We used a range of phylogenetic analyses to investigate the evolutionary relationships among Panguiarchaeales and Njordarchaeales (13 MAGs, **see Supplementary Data 1**), with Korarchaeota (Tahon et al. 2023) and Asgard archaea (officially referred to as Asgardarchaeota based on SeqCode (Tamarit et al. 2024) and as Promethearchaeota (Imachi et al. 2024) based on International Code of Nomenclature of Prokaryotes). Our starting point was the set of vertically evolved marker genes that have been used to investigate archaeal phylogeny and Njordarchaeia placement in several previous analyses, including ribosomal proteins as well as genes with alternative functions (Dombrowski et al. 2020; Williams et al. 2020; Rinke et al. 2021; Moody et al. 2022), totalling 120 distinct homologous Archaeal Clusters of Orthologous Genes (arCOG) gene families. We then performed a series of analyses to select the most reliable marker genes for concatenation, filtering out families that have been affected by gene replacement transfers from Bacteria (9 families) that were found in less than 60% of taxa (4 markers) or which have a complex history of gene duplication, transfer and loss within Archaea, and so are unreliable markers for inferring vertical evolution (11 families, including some subunits of DNA topoisomerase - **see Supplementary Information 1.1**, **Supplementary Data 2-4**, **Supplementary Figures 1-8**). We ranked the remaining markers based on split score quantifying the degree to which they recovered accepted monophyletic archaeal clades (Dombrowski et al. 2020; Liu and Li 2022; Moody et al. 2022) (see Methods: Ranking analysis, **Supplementary Data 4**) and selected the top 50% of these markers corresponding to 43 proteins(**Supplementary Data 4**) for phylogenetic analyses. The overlap between this marker set and those used in previous studies (Dombrowski et al. 2020; Williams et al. 2020; Liu et al. 2021; Moody et al. 2022; Eme et al. 2023; Qu et al. 2023; Valentin-Alvarado et al. 2024; Zhang et al. 2025), for both the initial and final curated set of markers, is summarised in **Supplementary Data 2-4**.

#### Phylogenetic analyses that model site- and branch- compositional heterogeneity place Panguiarchaeaceae and Njordarchaeaceae within Asgard archaea

Previous analyses have suggested that Panguiarchaeales and/or Njordarchaeales may branch either with Korarchaeota (Qu et al. 2023; Zhang et al. 2025) or Asgard archaea (Eme et al. 2023; Valentin-Alvarado et al. 2024) (**Figure 1A**). To distinguish between these hypotheses (i.e. Korarchaeota versus Asgard archaea topologies (**Figure 1A**)) and assess the taxonomic ranks of these clades, we performed phylogenetic analyses on the 43-gene concatenation using distinct taxon sets as well as relative evolutionary distance (RED) scaling to normalise the taxonomic ranks (Rinke et al. 2021; Tamarit et al. 2024) (see Methods and **Supplementary Figure 1** for method overview). All analyses provided strong support for the monophyly of the previously proposed Njordarchaeales and Panguiarchaeales clades, together forming a putative class-level taxon, i.e. the Njordarchaeia. This contrasts previous findings which suggested that Panguiarchaea are part of the Korarchaeota (Qu et al. 2023) while Njordarchaeales were assigned to the Asgard archaea (Eme et al. 2023; Valentin-Alvarado et al. 2024). Specifically, our results reveal that the Njordarchaeia comprises the Njordarchaeales order with two family-level lineages, the *Panguiarchaeaceae* and *Njordarchaeaceae* (**Figure 1BC, Supplementary Figure 9**, See also **Supplementary Information 1.2**). Furthermore, analyses using the best-fitting substitution models robustly supported the placement of the Njordarchaeia within the Asgard archaea (**Figure 1BC**, **Supplementary Figure 1**). However, we did observe the Korarchaeota topology, i.e. the placement of all Njordarchaeia with Korarchaeota, in several analyses using simpler substitution and empirical distribution mixture (EDM) models with few components (LG+G+F, LG+EDM0004LCLR+G+F, LG+EDM0008LCLR+G+F, **Supplementary Figure 1** and **Supplementary Figure 10A-C**), consistent with the hypothesis that this inferred relationship might be a phylogenetic artefact (Eme et al. 2023).

**Figure 1.**
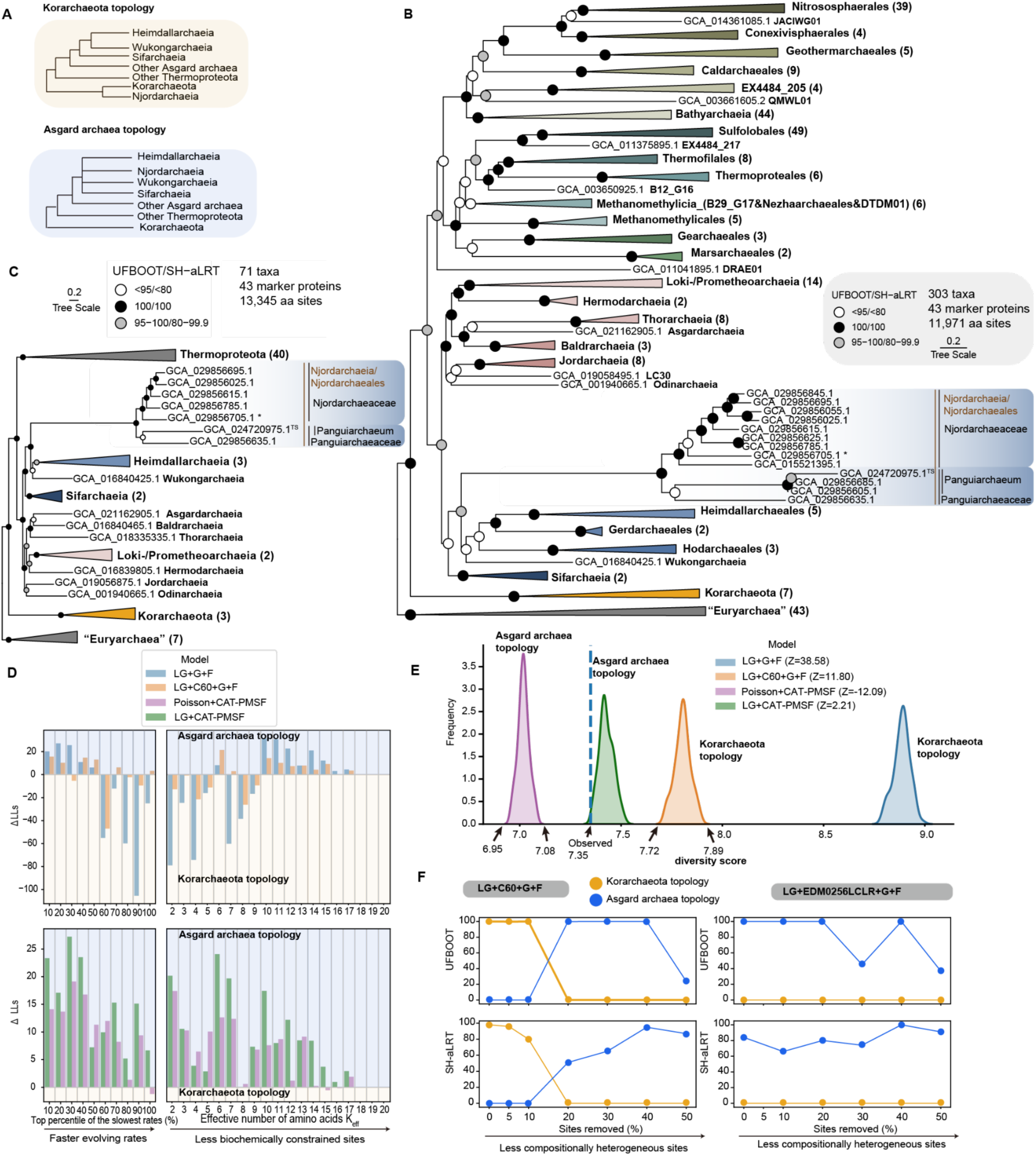
The phylogenetic placement of Njordarchaeia. **A**. Schematic of the phylogenetic hypotheses tested in this study. Korarchaeota topology: Njordarchaeia branch sister to Korarchaeota. Asgard archaea topology: Njordarchaeia branch within Asgard archaea. **B**. Maximum-likelihood phylogenetic analysis of the concatenated 50% top-ranked marker proteins (n=43) and streamlined set2 (303 taxa) using a custom site-heterogeneous substitution model (LG+EDM0256LCLR+G+F). Scale bar: average substitutions per site. The number of taxa in each collapsed clade is shown by the number in parenthesis next to the clade name. GCA_029856705.1* is the closest genome to the first proposed type material of *Njordarchaeum guaymasis* (See Supplementary Discussion). The GCA_024720975.1^TS^ is the proposed type material for *P.* symposium. The tree is rooted by “Euryarchaea” (including Halobacteriota, Thermoplasmatota, Hydrothermarchaeota, Methanobacteriota, Methanobacteriota_A and Methanobacteriota_B). **C**. Maximum-likelihood phylogenetic analysis of the concatenated 50% top-ranked marker proteins (n=43) and streamlined set3 (71 taxa) using LG+CAT-PMSF model. Scale bar: average substitutions per site. The number of taxa in each collapsed clade is shown by the number in parenthesis next to the clade name. **D**. Evaluation of phylogenetic signals for Asgard archaea and Korarchaeota topology under LG+G+F, LG+C60+G+F, Poisson+CAT-PMSF and LG+CAT-PMSF models, listed in an order of increasing model fit. Sites underlying the signals were binned by substitution rates (based on LG+C60+G+F) and effective amino acids per site (based on LG+CAT-PMSF). **E.** Distribution of amino acid diversity scores for the simulated alignments under LG+G+F, LG+C60+G+F, one representative Poisson+CAT-PMSF and LG+CAT-PMSF. The dashed line indicates the amino acid diversity score from the real dataset. *Z*-scores quantifying the difference between the compositional heterogeneity of the real dataset and that of the simulated datasets from each model are labelled. Note that *Z*-scores of LG+C60+G+F and Poisson+CAT-PMSF are similar but right tail of Poisson+CAT-PMSF is closer to real dataset, indicating a better fit. **F.** Impact of progressive removal of the most compositionally biased sites ranked by chi-square score on the statistical support of Asgard archaea and Korarchaeota topology. The support values, UFBOOT and SH-aLRT, were estimated on the 50% top-ranked markers of streamlined set2 (303 taxa) based on LG+C60+G+F and LG+EDM0256LCLR+G+F model.

Current phylogenetic methods are not consummate, and even the best models are mere approximations of the evolutionary processes that gave rise to the observed data. However, previous work has identified two aspects of the evolutionary process that are not well-captured by simple models, which have, in previous case studies, had a substantial impact on phylogenetic inference (Williams et al. 2020; Muñoz-Gómez et al. 2022). These are variations in amino acid preferences across the alignment sites (due to varying protein biochemical constraints) and across the tree branches (due to varying environmental pressures and mutational biases).

##### Modelling site-wise compositional heterogeneity

Site-wise compositional variation results from selective constraints on protein function: typically, only a small subset of the twenty possible amino acids are permissible at a given position in the protein due to the requirement for particular types of side chains to maintain protein structure and function (Lartillot and Philippe 2004). While being a pervasive feature of molecular sequence data, this site-wise variation is not accounted for in simple substitution models such as LG+G, which assume that all sites have the same equilibrium amino acid frequencies (Kapli et al. 2021; Williams et al. 2021). As they do not account for site-specific selective constraints, simple models under-estimate the probability of convergent evolution to the same amino acid in independent, distantly-related lineages. This means they can sometimes mis-interpret instances of molecular convergence as evidence that these taxa are closely related, an artefact that is expected to be strongest at the most selectively-constrained positions. To evaluate whether under-estimation of convergent evolution might drive the support for the cladehood of Njordarchaeia and Korarchaeota in analyses with simpler models, we grouped amino acid sites by compositional constraint (**see Methods**) and investigated support for the two trees from each class of sites (**Figure 1D**). This analysis revealed that the support for the Korarchaeota tree in simpler/underfitting model (**Figure 1E, assessed using model adequacy test** (Giacomelli et al. 2025)), LG+G+F/LG+C60+G+F, was greater at the more compositionally constrained sites, i.e., effective number of amino acids (K_eff_) ≤ 9, whereas with the best-fitting model and second best-fitting model, LG+CAT-PMSF and Poisson+CAT-PMSF (**Figure 1E, assessed usingmodel adequacy test** (Giacomelli et al. 2025)), both constrained and compositionally variable sites supported the Asgard archaea tree (**Figure 1D)**. Additionally, under the best-fitting model LG+CAT-PMSF and second best-fitting model Poisson+CAT-PMSF, Asgard archaea tree were consistenly recovered (**Supplementary Figures 10-13, Figure 1E**), and the Korarchaeota tree is confidently rejected by the approximately unbiased (AU) test (Shimodaira 2002) (P < 0.05) (**Supplementary Data 5**). These analyses indicate that failure to model site-wise compositional heterogeneity contributes to the incorrect placement of Njordarchaeia with Korarchaeota in analyses with poorly-fitting models such as LG+G+F and LG+C60+G+F (**Figure 1E**).

##### Modelling branch-wise compositional heterogeneity

In addition to site-wise variation, sequence composition can also vary among lineages with, for example, distantly-related thermophiles converging on greater use of thermostable amino acids (Ile, Val, Tyr, Trp, Arg, Glu, Leu (IVYWREL (Zeldovich et al. 2007)) or charged amino acids: Asp, Glu, Lys, Arg (DEKR) (Cambillau and Claverie 2000; Szilágyi and Závodszky 2000)). Njordarchaeia and Korarchaeota were both predicted to be (hyper-)thermophiles, and the placement of Njordarchaeia with Korarchaeota was suggested to be a result of compositional biases (Eme et al. 2023). In line with a previous study (Eme et al. 2023), a comparison of proteome composition across Archaea indicated that they indeed have similar amino acid compositions (**Supplementary Figure 14AB**). We subsequently investigated whether compositional attraction between the two lineages might also contribute to the recovery of the Korarchaeota tree under simple models. First, we performed a site-filtering analysis. We ranked sites by compositional bias, iteratively removing the sites that made the greatest individual contribution to across-branch compositional heterogeneity (Dombrowski et al. 2020; Baker et al. 2024) (**see Methods**), and re-evaluated support for the competing topologies (**Figure 1F**). Removal of the top 20% of most biased sites shifted support from the Korarchaeota topology to the Asgard archaea topology under LG+C60+G+F, while under the better-fitting and custom site-heterogeneous model, LG+EDM0256LCLR+G+F, support was unchanged, with all analyses favouring the Asgard archaea topology. We interpret these results as evidence that compositional attraction between thermophilic branches favours the Korarchaeota topology in the full dataset, but whether this non-phylogenetic signal drives the final result depends on other aspects of the analysis. While neither the LG+C60+G+F model nor the LG+EDM0256LCLR+G+F model account for across-branch compositional variation, C60 fares worse than EDM00256 in modelling across-site variation (**Supplementary Figure 11**; Bayesian information criterion (BIC) of LG+C60+G+F: 6668927.72, LG+EDM0256LCLR+G+F: 6532626.81). We therefore suggest that support for the Korarchaeota topology from the 303 taxa set in the C60 analysis reflects a combination of phylogenetic artifacts from both unmodelled branch-wise compositional attraction and inadequately-modelled site-wise compositional variation (that is, long branch attraction), with support shifting to the Asgard archaea topology when one of these sources of error is removed, either by filtering out highly biased sites or by better modelling of site-wise compositions with LG+EDM0256LCLR+G+F.

Second, we identify RYPEIWKVL and QSNTDGC, as overrepresented and underrepresented amino acids, respectively, in the concatenation (Zeldovich et al. 2007) (**Supplementary Figure 14CD**). Subsequently, we performed phylogenetic analysis using a substitution model that accounts for changing compositions across the tree, i.e. GFMix (Muñoz-Gómez et al. 2022). The LG+EDM00256LCLR+G+F+GFMix (RYPEIWKVL/QSNTDGC and RYPEI/QSNT) and LG+C60+G+F+GFmix (RYPEIWKVL/QSNTDGC and RYPEI/QSNT) model provided strong support for the Asgard archaea tree over the Korarchaeota tree (**Supplementary Data 6**), again consistent with the hypothesis that compositional attraction due to a shared environmental adaptation contributes to the Korarchaeota tree in analyses with models that do not account for branch heterogeneity (Eme et al. 2023).

#### Is the inferred phylogenetic position of Panguiarchaeceae and Njordarchaeceae affected by metagenomic contamination?

The phylogenetic position of *Njordarchaeia* has been the subject of debate based on different marker sets. A recent analysis (Dombrowski et al. 2020; Zhang et al. 2025) argued that these MAGs may be chimeric, comprising sequences derived from both Thermoproteota and Asgard archaea, which contributes to conflicting phylogenetic placements observed in different marker sets. It has been pointed out that MAGs often contains various degree of contamination (Parks et al. 2015; Shaiber and Eren 2019; Chen et al. 2020), and the MAGs analysed in this study are no exception as assessed by CheckM and differential coverage (**Figure 2; Supplementary Data 1**). Nevertheless, the key question is whether the marker genes used to place these lineages are affected by contamination. Using the same analytical approach, we inspected the distribution of marker genes, and within-bin coverage variation of Panguiarchaeceae and Njordarchaeceae MAGs across different metagenomes from hydrothermal vents in the Guaymas Basin (Dombrowski et al. 2018) and a hot spring in Yunnan, China (Qu et al. 2023). CheckM analysis indicates that all representative Njordarchaeia MAGs analysed here (**taxon set 303, Supplementary Data 1**) are at least medium quality (Bowers et al. 2017) with only a moderate level of contamination (<10%), with 53% (7 MAGs) having low level of contamination (<5%) (**Supplementary Data 1**). MAGs with/above medium quality are the majority in different databases such as the Genomes from Earth’s Microbiomes catalogs (Nayfach et al. 2021), and reported in various studies (Dombrowski et al. 2020; Martijn et al. 2020; Zhang et al. 2025). To investigate whether our marker gene phylogenies are nevertheless affected by this low level of contaminations, we chose to examine 4 MAGs from our focal taxon set that derive from hydrothermal vents in Guaymas basin and a hot spring from China (estimated contamination: 0.93% to 7.01%; **Figure 2**) that were also examined in a previous study (Zhang et al. 2025). While our analyses confirmed the presence of small potentially contaminating contigs (**Figure 2ABCD**), all marker genes mapped back to a single majority cluster corresponding to the target population, even in the MAG estimated with the highest level of contamination (**Figure 2C**). This suggests that contamination, though present at a low/moderate level in some of these MAGs, does not affect the marker genes used, and so does not underpin the phylogenetic conflicts. Instead, conflicting support appears to depend on the substitution model used. For example, analyses of two marker genes situated within a non-contaminated contig (**Figure 2ABC, highlighted in red box**) under the poorly-fitting LG+F+G and LG+C60+F+G models differed in their support for the Asgard and Korarchaeota topologies, whereas analysis of the same marker genes under the best-fitting LG+CAT-PMSF model provides moderate support for Asgard archaea topology (**Figure 2ABC**). We also observed in our dataset that LG+C60+G+F did not significantly distinguish these two topologies (AU tests (Shimodaira 2002), P < 0.05; **Supplementary Data 5)** nor describe site-heterogeneity adequately **(Figure 1E**). However, as model fit improved, consistent support for the Asgard topology emerged (**Supplementary Data 5, Figure 1DEF**).

**Figure 2.**
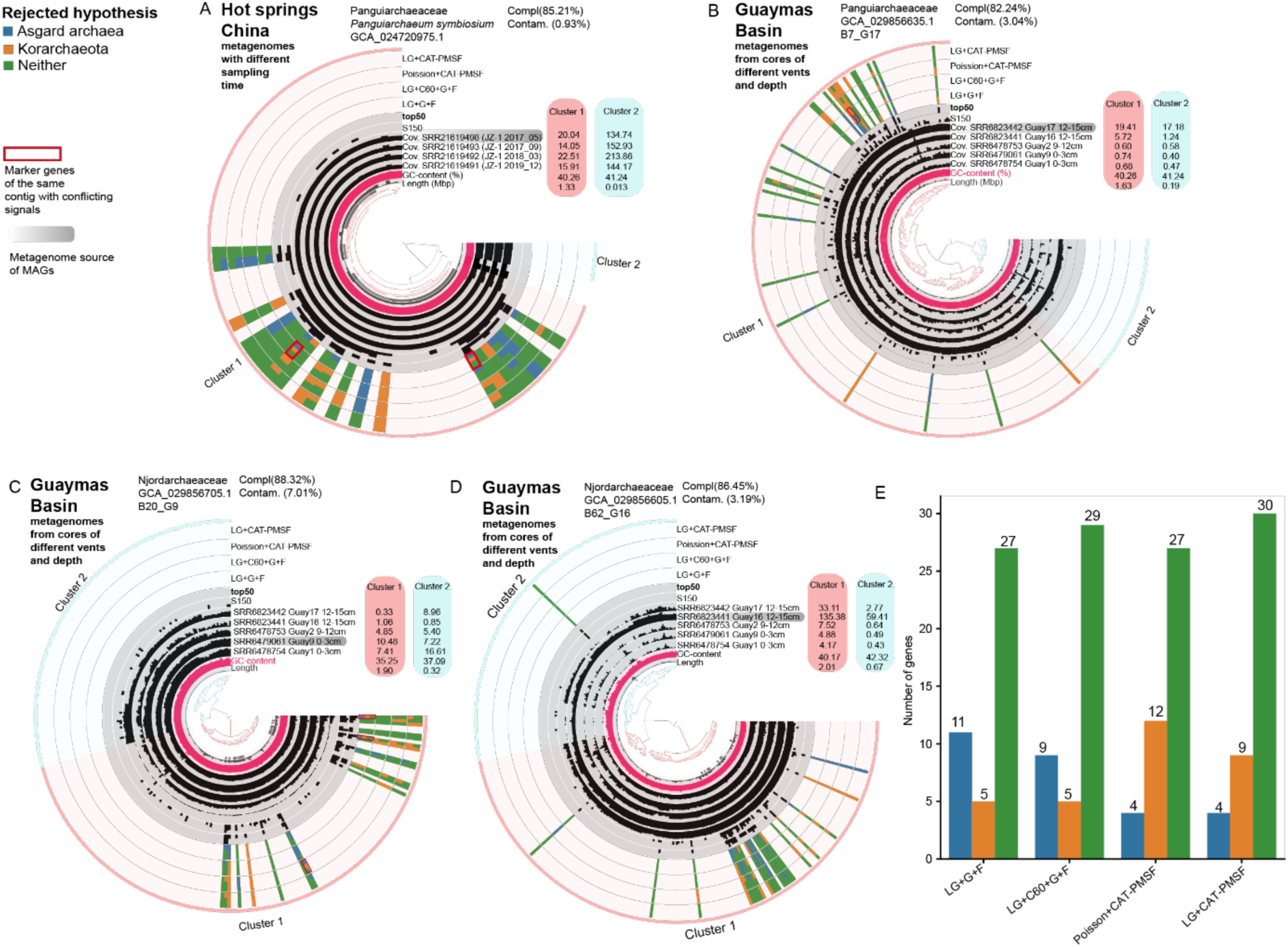
Hierarchical clustering of contigs in four MAGs from Panguiarchaeaceae (A, B) and Njordarchaeaceae (C, D) based on sequence composition and depth of coverage across different metagenomes. Note that the tip of hierarchical clustering tree represents splits of long contigs at a size of 20000 bp. This analysis demonstrates metagenomic contamination is not a source of conflicting phylogenetic signals. The ring indicating coverage across the source metagenomes is highlighted in a gray box. The locations of marker gene sequences used in this study (ring labeled **top50**) and marker gene homologs from a previous study (ring labeled **S150**) are shown as bar plots, with bar height corresponding to the number of genes. Notably, our marker gene sequences are located on clusters of contigs that share similar coverage profiles across metagenomes, as are the majority of the **S150** homologs. Support for Asgard archaea and Korarchaeota topologies is shown in a stacked bar plot, with genes on the same contig that show conflicting topological support highlighted in red boxes. **E**. Barplot showing the number of genes that confidently reject Asgard archaea, Korarchaeota or neither of the two topologies under different models.

While we saw no evidence of contamination in our analyses, we did see some variation in support for the Korarchaeota or Asgard topologies on a per-gene basis when the data were analysed using different models. To quantify disagreements among marker genes, we used AU tests (Shimodaira 2002) to determine, for each marker gene and substitution model, whether either the Korarchaeota tree or Asgard archaea tree could be rejected (at P < 0.05). Across all models tested, 27 of 43 marker genes did not reject either hypothesis, suggesting that individual marker genes often do not contain sufficient information to discriminate among phylogenetic hypotheses (**Figure 2E**). For the remaining marker genes, support on a per-gene basis shifted from the Korarchaeota to the Asgard archaea topology as model fit (assessed using model adequacy test (Giacomelli et al. 2025)) improved. Under the worst-fitting LG+G+F model, 5 genes rejected the Korarchaeota topology and 11 rejected the Asgard topology at the significance level (**Figure 2E**). Under the site-heterogenous LG+C60+G+F model, 5 genes rejected Korarchaeota and 9 rejected Asgard archaea. Under the best-fitting (LG+CAT-PMSF) and second best-fitting (Poisson+CAT-PMSF) models, 9 and 12 genes, respectively, rejected the Korarchaeota topology, while 4 rejected the Asgard topology (**Figure 2E**). The improvement in congruence across marker genes and the shift in support to the Asgard topology as model fit improves suggests that the apparent strong but inconsistent support observed in analyses under the poorly-fitting LG+G+F and LG+C60+G+F models is artefactual, resulting from the failure of these models to adequately account for non-historical signal in sequence data (**Figure 1E**; **Supplementary Information 1.3-1.4**).

#### Assignment of Njordarchaeia to Asgard archaea is supported by shared eukaryotic signature proteins (ESPs) and protein presence-absence profiles

The phylogenetic analyses described above provided strong support for the placement of *Panguiarchaeaceae* within Njorarchaeales as a sister to *Njordarchaeaceae* within the Asgard archaea (**Figure 1B-F**). Consistent with that placement, and in line with (Eme et al. 2023), we observed, we observed that *Panguiarchaeaceae* and *Njordarchaeaceae* encode a number of eukaryotic signature proteins (ESPs) shared with other Asgard archaea **(Figure 3, Supplementary Data 7**), based on analyses using the Asgard Cluster of Orthologues (AsCOGs) database (Liu et al. 2021). Of particular interest, we identified homologs of ribosomal L28e/Mak16 (PF01778) in *Panguiarchaeaceae* MAGs, whose distribution in published archaeal genomes is otherwise restricted to other Njordarchaeia, the Hodarchaeales, and Wukongarchaeia (Eme et al. 2023). Phylogenetic analysis of ribosomal L28e/Mak16 revealed a topology consistent with the species tree **(Supplementary Figure 15, Supplementary Information 2.2.1**), with the Njordarchaeia sequences branching sister to Wukongarchaeia and other Heimdallarchaeia. Furthermore, *Panguiarchaeaceae* MAGs encode an actin homolog that branches with the conserved lokiactin clade present in all asgard archaeal lineages (**Supplementary Figures 16-17, Supplementary Information 2.2.2**). Homologs of proteins involved in the N-glycosylation process were also identified in *Panguiarchaeaceae* as well as *Njordarchaeaceae* MAGs (Eme et al. 2023), and the phylogenetic analyses support their affiliation with Asgard archaeal homologs (**Supplementary Figures 18-19, Supplementary Information 2.2.3**). Phylogenetic analyses of Njordarchaeia SNF7 proteins also support them as Asgard archaeal homologs (**Supplementary Figure 20, Supplementary Information 2.2.4**).

**Figure 3.**
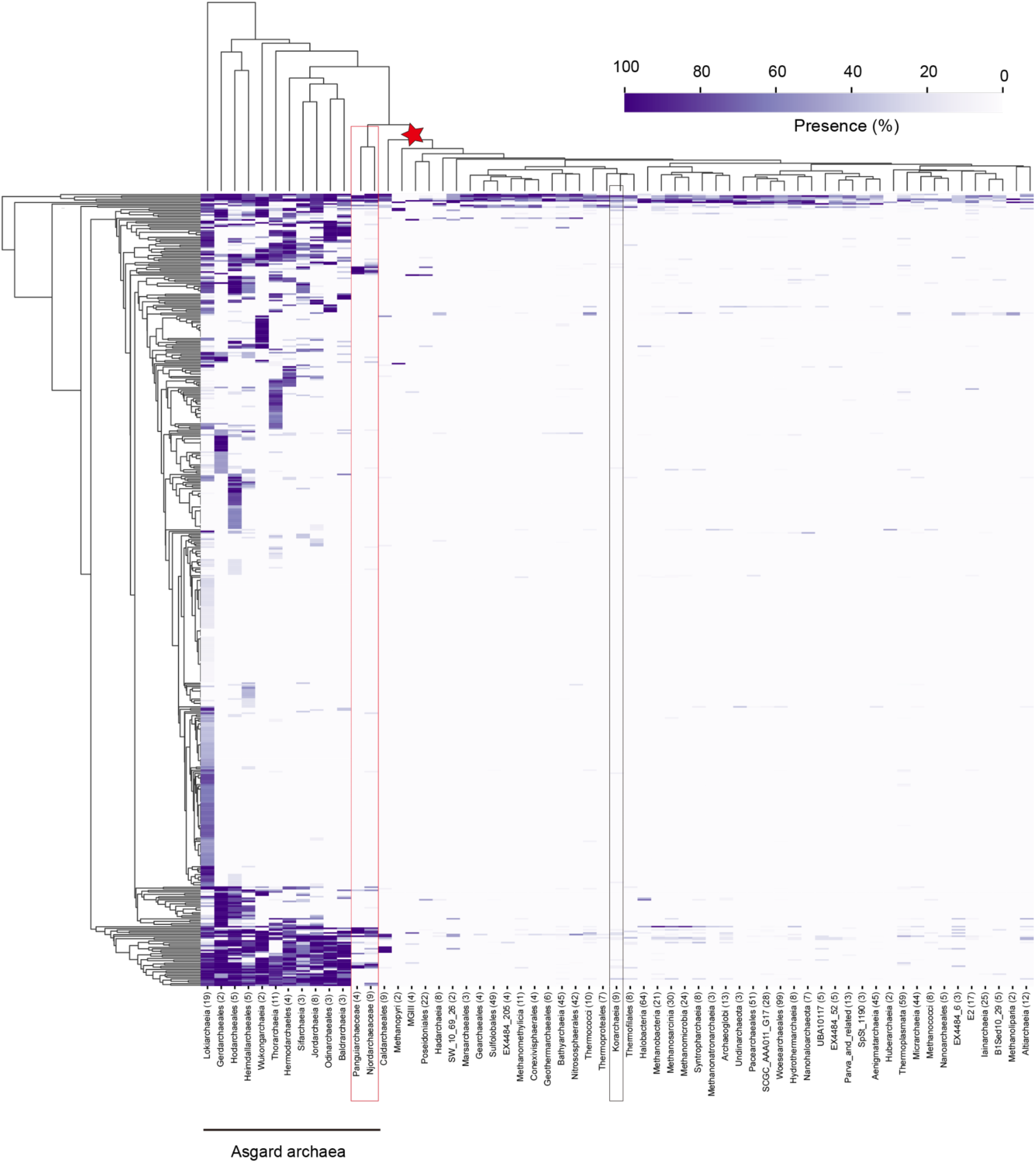
Presence of Asgard cluster of orthologues (AsCOGs) across major archaeal lineages. Panguiarchaeceae and Njordarchaeceae shared a number of AsCOGs homologs with other Asgard archaea albeit reduced, which is distinct from Korarchaeota. AsCOG presence was determined across 966 archaeal genomes, and the occurrence of AsCOGs within any taxonomic clade of interest (based on the genome taxonomy database) was recorded in percentage. Clusters with less than two genomes were not included. The number of genomes for each cluster is shown in parentheses.

Finally, protein presence-absence profiles revealed a pattern of transcription, translation and replication distinct from Korarchaeota (**Supplementary Figures 21-22**). Interestingly however, the profile for other COG categories was more similar to that of Korarchaeota than to other Asgard archaea (**Supplementary Figures 21-22**), likely reflecting adaptation to high temperature environment.

### Incomplete biosynthetic pathways indicate that *Panguiarchaeum* may depend on other organisms

Comparative genome analyses and the inference of metabolic potentials for Njordarchaeia MAGs (i.e. the *Panguiarchaeaceae* clade including *Panguiarchaeum* and GCA_029856635.1 and the *Njordarcheaceae* clade) suggest that these organisms are thermophilic fermentative heterotrophs in agreement with previous work (Qu et al. 2023) that have varying levels of auxotrophies; that is, they cannot synthesise all organic compounds needed for growth (**Supplementary Figure 23**). Of note, members of the *Panguiarchaeum* show a reduced metabolic potential, in particular in purine and lipid biosynthesis pathways, compared to Njordarchaeacea and Korarchaeota (**Supplementary Figure 23-27, Supplementary Information 3.1-3.2**).

#### Catabolism and energy conservation

The catabolic potential of the *Panguiarchaeum* clade (i.e. upon divergence from GCA_029856635.1) is more limited than that of *Njordarchaeaceae* (**Figure 4**, **Supplementary Data 8-10, Supplementary Figure 23**). In particular, while both clades encode a nearly complete Embden-Meyerhof-Parnas (EMP) pathway, members of *Njordarchaeaceae* encode more genes for enzymes of the tricarboxylic acid cycle (TCA) and beta-oxidation pathway. Furthermore, members of both clades might have the potential to grow on organic substrates, which could be fermented to acetate by a putative acetate-CoA ligase (ACD) with concomitant production of ATP (Glasemacher et al. 1997). Specifically, *Panguiarchaeum* representatives revealed a potential to harness electrons from organic substrates such as amino acids, peptides, 2-ketoacids (e.g., pyruvate, oxoglutarate, and indole-pyruvate) (Qu et al. 2023), and aldehydes (**Supplementary Data 8**, **Supplementary Information 3.2**). Additionally, we identified homologues of the small and large subunits of [NiFe]-hydrogenases belonging to group 3 and group 4 in *Panguiarchaeum* MAGs. Phylogenetic analyses of the large subunit of the [NiFe] hydrogenase indicated that panguiarchaeal homologs belong to subgroup 4g (Qu et al. 2023), forming a clade with Odinarchaeia sequences (**Supplementary Figure 28A**). Group 4g [NiFe]-hydrogenases of *Panguiarchaeum* encode a Nuo-L-like subunit, which might be involved in sodium/ion translocation (Yu et al. 2018) (**Supplementary Figure 28B**). This suggests that *Panguiarchaeum* may conserve energy by coupling the oxidation of organic substrates via reduced ferredoxin to the formation of H_2_ to generate a proton gradient for ATP synthase. Members of *Panguiarchaeum* also contain various group 3 [NiFe]-hydrogenases assigned to subgroup 3c and 3b (Qu et al. 2023), which may be involved in the reoxidation of cofactors during carbon oxidation and the production of H_2_, a reversible process (**Supplementary Figure 29, Supplementary Data 11**). Together, this indicates that representatives of the *Panguiarchaeum* clade can conserve energy through substrate-level phosphorylation as well as via the ATP synthase using a proton gradient generated by the oxidation of ferredoxin similar to Odinarchaeia and Thermococci (Spang et al. 2019; Qu et al. 2023).

**Figure 4.**
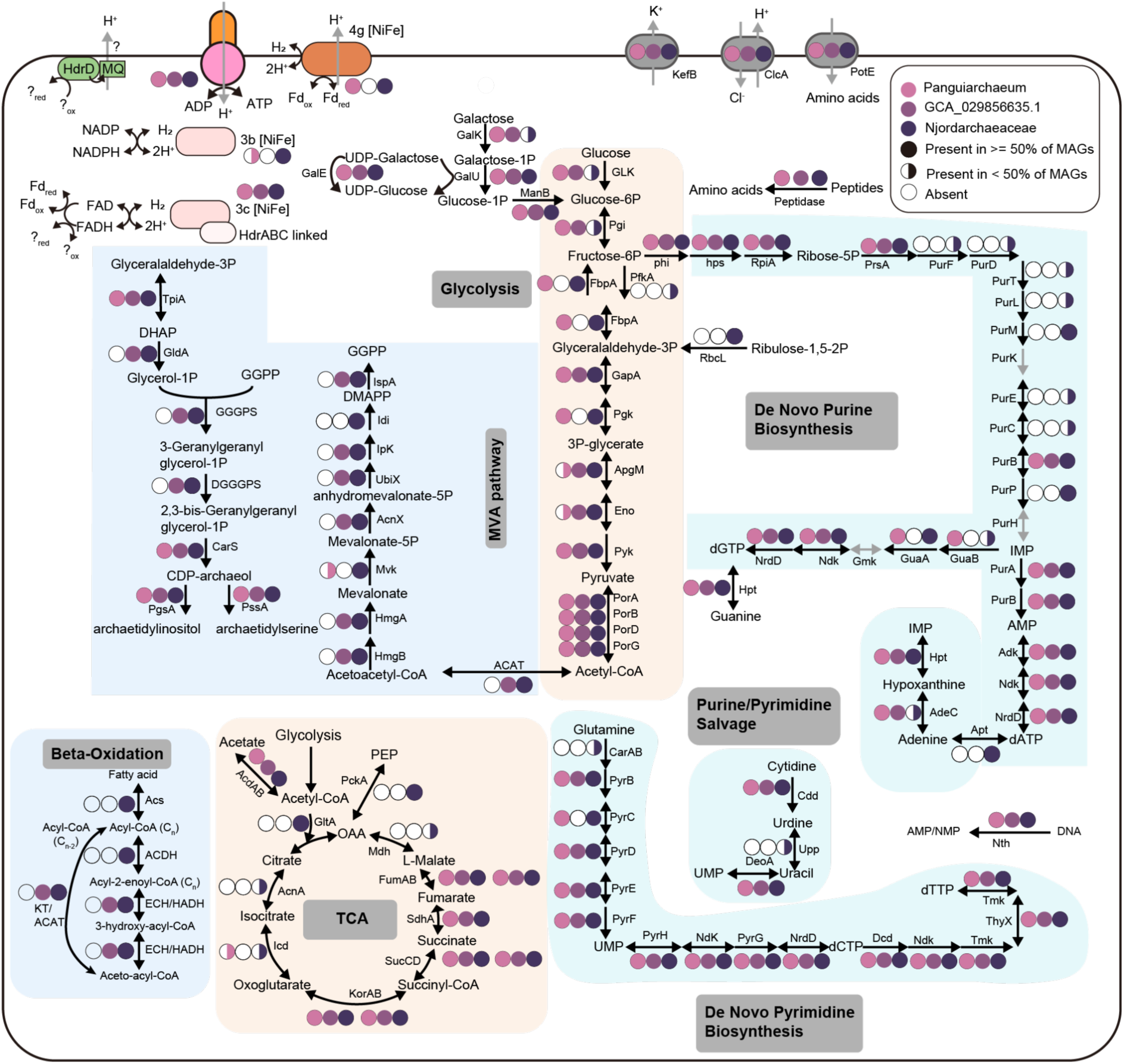
Metabolic characteristics of Njordarchaeia. Overall, Njordarchaeia are thermophilic fermentative heterotrophs, with core pathways of varying completeness suggesting some core metabolites must be obtained from other organisms in the environment. Full circles indicate that a gene is present in all or more than 50% of the MAGs, while half-circles indicate that a gene is found in at least one, but less than half, of MAGs. Open circles denote genes absent from all Njordarchaeia MAGs. A detailed list of genes encoded by Njordarchaeia can be found in Supplementary Data 7,8,9.

#### Anabolism

The biosynthetic potential of members of the *Panguiarchaeum* clade seemed more restricted than that of *Njordarchaeaceae* (**Supplementary Figure 23**). For instance, all three *Panguiarchaeum* MAGs lack various genes encoding enzymes for *de novo* purine biosynthesis (from phosphoribosyl pyrophosphate to inosine monophosphate) (Qu et al. 2023), such as glutamine phosphoribosylpyrophosphate amidotransferase (PurF) and Phosphoribosylaminoimidazole (AIR) synthetase (PurM) indicating a possible dependence on external purine sources (Brown et al. 2011). Notably, *Panguiarchaeum* clade MAGs do not encode enzymes of the mevalonate pathway (MVAP) for isoprenoid biosynthesis and fatty acid biosynthesis, as indicated previously (Qu et al. 2023). However, the presence of CDP-archaeol synthase (CarS), phosphatidylglycerophosphate synthase (PgsA), phosphatidylglycerophosphate synthase (PssA) and cardiolipin synthase (Cls), suggests that *Panguiarchaeum* have the potential to attach polar head groups to di-o-geranylgeranylglyceryl phosphate (DGGGP). The *Panguiarchaeum* MAGs also encode a gene for digeranylgeranylglycerophospholipid reductase (GGR), which could catalyse the hydrogenation/saturation of geranylgeranyl chains of DGGGP (Nishimura and Eguchi 2006). Phylogenetic analyses of CarS showed that the homologs of *Panguiarchaeum* branched sister to Heimdallarchaeia with moderate support (UFBOOT/SH-aLRT: 88.8/83; **Supplementary Figure 30**). Similarly, although the PgsA homologs of *Panguiarchaeum* branched sister to a clade comprising one Thermococcus and two Heimdallarchaeia sequences with weak support (UFBOOT/SH-aLRT: 50.5/83), they were nested within a cluster of predominantly Asgard archaeal sequences (**Supplementary Figure 31**), Thus, these genes were likely inherited vertically from their common ancestor with Heimdallarchaeia (**Supplementary Figure 30-32**).

### *Panguiarchaeum* has evolved by genome streamlining from a larger ancestor with Heimdallarchaeia

Reconciliation analyses have been used to infer the ancestral genome content of various archaeal lineages (Williams et al. 2017; Martijn et al. 2020; Huang et al. 2021; Baker et al. 2024; Williams et al. 2024), including Asgard archaea (Eme et al. 2023), shedding light on ancestral gene repertoires and gene content evolution. To reconstruct the evolution of a putative symbiotic lifestyle of *Panguiarchaeum*, we reconciled single gene trees of 6077 arCOG gene families with the inferred species tree (from streamlined set 2 with 303 taxa, **Figure 1B**) to determine the pattern and extent of reductive genome evolution in the *Panguiarchaeum* clade and its sister lineages.

Our analysis suggests an ongoing decrease in gene content through time in the evolution of Njordarchaeia, similar to a previous analysis (Eme et al. 2023), from around 1951 arCOG families in the common ancestor with Wukong-Heimdallarchaeia to 911-1545 (mean: 1186.5) arCOG families among extant Njordarchaeia (**Supplementary Data 12**). Based on the relationship between arCOG family members and the genome size of extant taxa (Pearson’s correlation coefficient: 0.89, P-value: 1.96×1e-110), we inferred the genome size of the last common ancestor of Wukong-Njord-Heimdallarchaeia to be 2.85 Mbp (95% confidence interval, 2.57 to 3.06, **Supplementary Data 12**). A notable reduction in genome size is inferred at the branch leading to the Njordarchaeia (from 0.53 to 0.84 Mbp; node id: 571-563) after divergence from their common ancestor with Wukong-Heimdallarchaeia (**Figure 5A**, **Supplementary Data 12**). The arCOG families lost along this branch include components of amino acid biosynthesis, de novo purine biosynthesis (from ribose-5-phosphate to inosine monophosphate (IMP)), Wood-Ljungdahl pathway genes and cofactors (methanofuran) biosynthesis (**Figure 5B**, **Supplementary Data 13**). Additional genes, including those encoding various enzymes involved in archaeal lipid biosynthesis and beta-oxidation, appear to have been lost along the branch leading to the *Panguiarchaeum* clade after its divergence from GCA_029856635.1 and the common ancestor with *Njordarchaeaceae*, which resulted in an inferred genome reduction of 0.13 Mbp to 0.17 Mbp. *Panguiarchaeum* and *Njordarchaeaceae* were inferred to encode fewer ESPs than the other Asgard archaea (**Figure 3**), and the loss on the *Njordarchaeaceae* and *Panguiarchaeaceae* stem is predicted to include the reduction in ESP repertoire as well. For instance, we inferred the loss of a Roadblock/LC7 domain-containing protein (arCOG02605_01) on the *Njordarchaeaceae* branch; the loss of Ras family GTPase (arCOG05343_01) and Yip1 domain family (arCOG02054_01) on the *Panguiarchaeaceae* branch; the loss of VPS4 (arCOG01307_01) on the *Panguiarchaeum* branch (i.e. upon divergence from GCA_029856635.1) (**Supplementary Data 13**); and the absence of Snf7 family proteins in the extant *Panguiarchaeum* MAGs indicated their loss on the *Panguiarchaeum* branch (**Supplementary Figure 20**).

**Figure 5.**
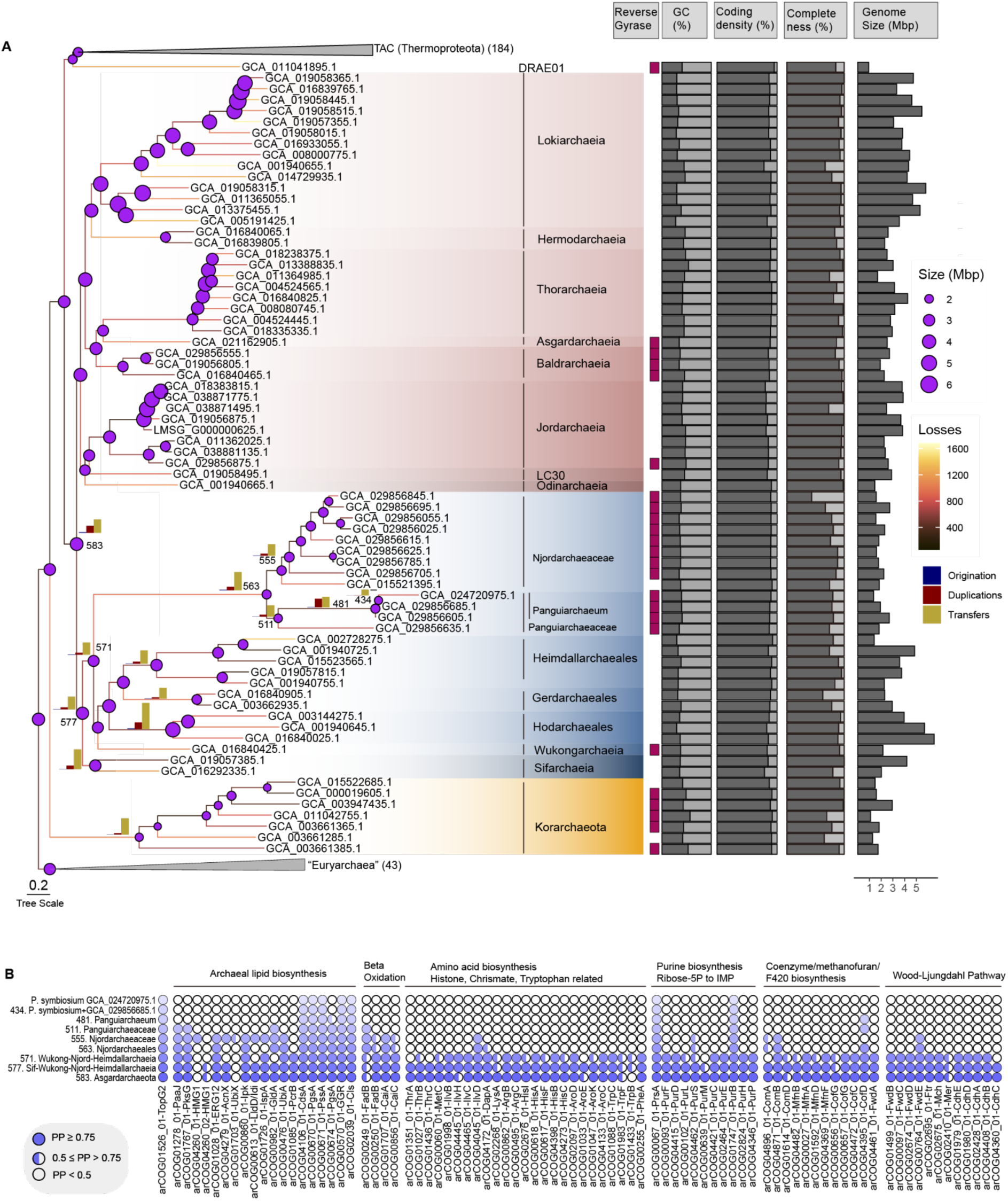
Gene-tree-aware ancestral reconstruction of *Panguiarchaeum* lineages. Njordarchaeia, in particular *Panguiarchaeum*, might have experienced genome reduction from their common ancestor with a larger ancestor Wukongarchaeia and Heimdallarchaeia (0.53-0.84 Mpb on the stem: node 571 to node 563). A. Inferred species of asgard archaeal, thermoproteotal and korarchaeotal lineages from Figure 1B. The colours on the branches represent the number of gene losses, and the bar plots on the branches of interest show the number of originations, duplications and transfers. The size of circles at nodes represents the inferred ancestral genome size. The number of taxa in each collapsed clade is shown by the number in parenthesis next to the clade name. B. The presence probability of genes of interest at the ancestral nodes leading to *Panguiarchaeum*. Node numbers of ancestral nodes are shown and referred to in Panel A. Full circles indicate a presence probability (PP) ≥ 0.75, half circles show a PP< 0.75 but ≥ 0.5, while PP<0.5 is shown in open circles (See Supplementary Data 12).

Njordarchaeia encodes reverse gyrase, a topoisomerase typically encoded by hyperthermophiles (Forterre 2002). Consistent with previous work (Eme et al. 2023) and proposal of a thermophilic ancestry of Asgard archaea (Lu et al. 2024), we inferred the presence of reverse gyrase (Presence probability, PP = 0.96) in the last common ancestor of Asgard archaea. This gene family was, however, subsequently lost in Heimdallarchaeia (**Supplementary Data 13**) but not in Wukongarchaeia and Njordarchaeia. Interestingly, phylogenetic and reconciliation analyses suggest a later re-acquisition of a reverse gyrase in the *Panguiarchaeum* clade (**Supplementary Figures 33-34; Supplementary Information 3.1.2**). The MAG/genome size of putative hyperthermophilic Asgard archaea encoding reverse gyrase (mean: 2.08 Mbp, n=18) is significantly smaller than that of Asgard archaea representatives without reverse gyrase (mean: 3.54 Mbp, n=45) (p=5.669×10-7, two-tailed Wilcoxon rank sum test). Genome reduction has previously been reported in cases of thermophilic adaptation, perhaps due to a selective advantage for smaller cell volumes (and so genome) at high temperatures (Sabath et al. 2013; Pierpont et al. 2024). We therefore hypothesise that the initial reductive evolution of Njordarchaeia might have been associated with their adaptation to high-temperature environments. The further genome reduction in *Panguiarchaeum*, including the loss of essential genes for biosynthetic pathways, might be associated with a transition to a host-associated lifestyle (McCutcheon and Moran 2011).

### *Panguiarchaeum* co-occurs with Thermoproteota taxa

To investigate the inferred dependence of *Panguiarchaeaum* on other community members, we assessed the extent to which members of this group co-occur with other microbes using network inferences. First, we tested the inference of our co-occurrence networks with the known host-symbiont pair *Nanoarchaeum equitans* and *Ignicoccus hospitalis* (Huber et al. 2002), and the proposed symbiosis between *Huberarchaeum crystalense* and *Altiarchaeum hamiconexum* (Schwank et al. 2019). We did not detect any associations between the former, mainly due to insufficient sampling of *N. equitans* which is present in a single sample only (**Supplementary Figure 35-36, Supplementary Information 4**). However, for *Huberarchaeum*-*Altiarchaeum,* we found a high correlation (Spearman’s rho = 0.931) and a robust detection frequency (100%) of this suggested interaction starting at a sample size of 40 (**Supplementary Figure 36F; see Methods**). Next, we identified 120 samples containing taxa annotated as family *Panguiarchaeaceae* in GTDB (Rinke et al. 2021), a lineage equivalent to the order Njordarchaeales proposed in this manuscript, from 248,559 NCBI metagenomes community profiles in the ‘Sandpiper’ website (https://sandpiper.qut.edu.au). Community profiles of these 120 samples were subjected to various data transformations (**see methods**) and network inferences. Across all original networks, i.e. independent of filtering criteria applied before network inference, Njordarchaeales taxa were associated with Thermoproteota taxa. The frequency of association between these taxa was up to ∼4 times higher than that between Njordarchaeales and any other taxa or random number controls. On phylum, class, and order level Njordarchaeales were associated with Thermoproteota (73 out of 256), Thermoprotei_A (33 of 73), and Sulfolobales (23 of 33) (**Supplementary Data 14-15**), respectively, in agreement with abundance-based co-occurrence patterns (**Figure 6)**. Within the order Njordarchaeales, there was only one OTU (OTU_804; **Supplementary Data 14**) assigned to the genus *Panguiarchaeum*, that found association partners in the network. This genus had 29 associations in total, whereby the taxonomic annotation of its partners was similar to the associations detected for the order, i.e. Thermoproteota (15 of 29), Thermoprotei_A (9 of 15), and Sulfolobales (4 of 9), and *Desulfurococcaceae* (3 of 4) (**Supplementary Data 15**). These results indicate that members of the Njordarchaeales, in particular the genus *Panguiarchaeum*, might interact with members of the phylum Thermoproteota, in particular with Sulfolobales, an order comprising hosts of thermophilic Nanoarchaeota (Huber et al. 2002; Podar et al. 2013; Munson-McGee et al. 2015; Wurch et al. 2016; St John et al. 2019; Kato et al. 2022; Sakai, Nur, et al. 2022).

**Figure 6.**
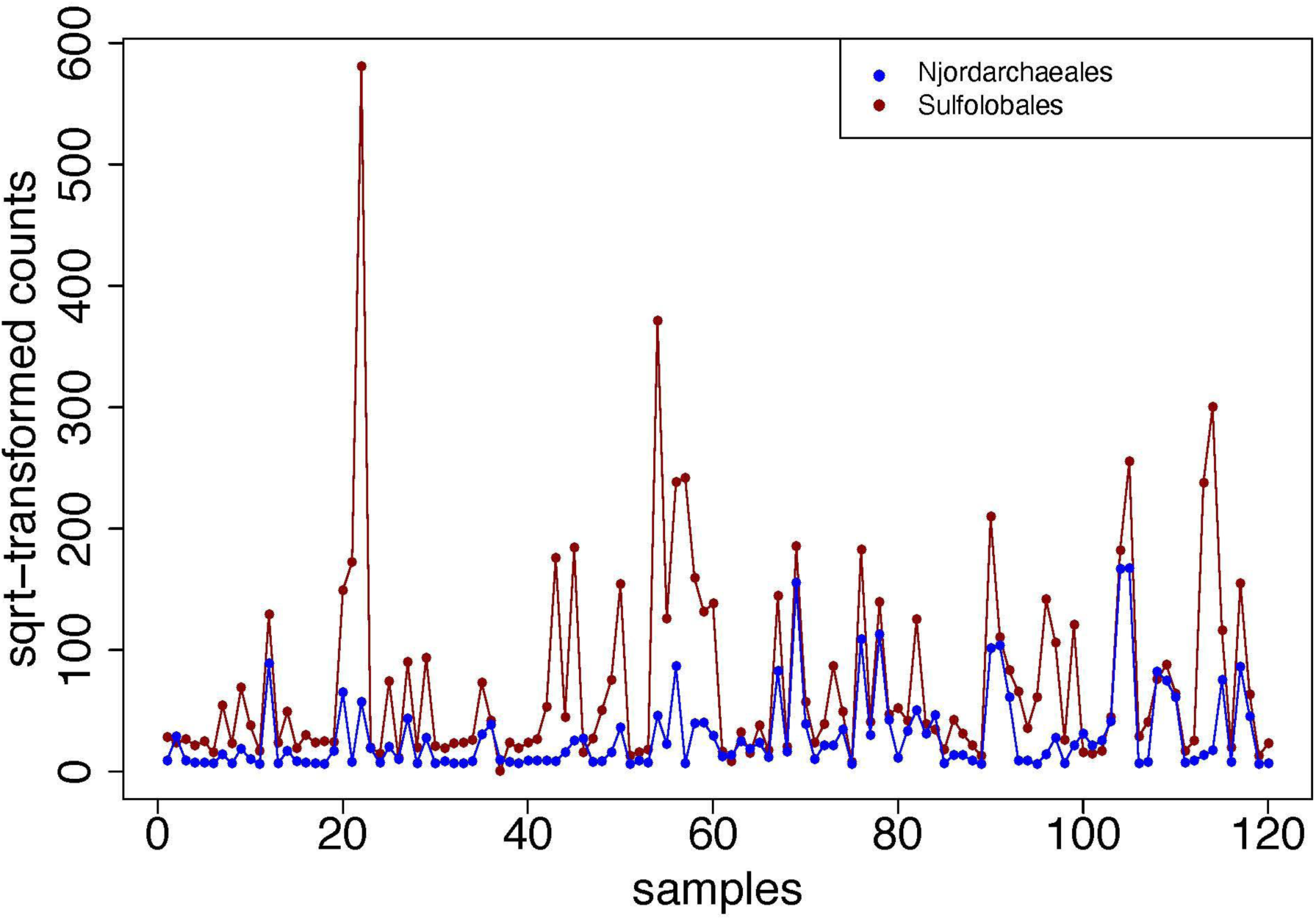
Abundance-based co-occurrence of Njordarchaeales and Sulfolobales. For plotting, the counts of both orders Njordarchaeales (including Panguiarcheaceae) and Sulfolobales, were summed in each sample and plotted. For visualisation purposes, the counts were square root transformed.

## Discussion

Reconstructing the tree that best depicts the evolutionary relationship between major archaeal lineages is important for our understanding of archaeal genome evolution, the evolution of archaeal symbioses and eukaryogenesis. Our comprehensive phylogenetic and comparative genomic analyses reveal that *Panguiarchaeum* together with GCA_029856635.1 forms a family within the Njordarchaeia as part of the Asgard archaea. Thus, the previous placement of the *Panguiarchaeum* clade as a sister group to Korarchaeota appears to be due to phylogenetic artefacts (Qu et al. 2023) including the use of problematic marker gene families in concatenations and insufficient modelling of compositional biases and site heterogeneity similar to what has been shown for Njordarchaeales (Eme et al. 2023). In contrast, our analyses did not provide support for the recent suggestion that phylogenetic placement of Njordarchaeia has been affected by metagenomic contamination (Zhang et al. 2025). While some Njordarchaeaia MAGs do show evidence of low/moderate levels of contamination (**Figure 2**), we found no evidence that marker genes were affected in any case. Our analyses instead suggest that disagreement among marker genes as to the placement of Njordarchaeia is a result of poor-fitting phylogenetic models, not MAG chimerism. These results emphasise the importance of evolutionary model fit in analyses of prokaryotic evolution, particularly for datasets containing pervasive compositional heterogeneity.

The placement of *Panguiarchaeum* within the Asgard archaea is supported by the presence of various ESPs shared with *Njordarchaeaceae*. However, metabolic reconstructions reveal that members of *Panguiarchaeum* have various auxotrophies and encode fewer ESPs than other Asgards. This is indicative of genome reduction potentially associated with a host-associated lifestyle, with co-occurrence analyses pointing towards an archaeal host. Notably, *Ca*. P. syntrophicum, the first cultivated representative of the Asgard archaea (i.e. a member of the Lokiarchaeia/Prometheoarchaeia), appears to syntrophically grow on amino acids with partner organisms such as *Halodesulfovibrio* and *Methanogenium*^13^. However, this symbiotic interaction does not seem to be obligate and has not resulted in reductive genome evolution. Our inferences of the energy metabolism of *Panguiarchaeum,* and perhaps Njordarchaeaceae as well, which seems to resemble that of Odinarchaeia and *Thermococcus*, indicates the potential for syntrophic growth (Schut et al. 2013; Topçuoğlu et al. 2016; Yu et al. 2018; Spang et al. 2019). For example, *Panguiarchaeum* encodes 3 different [NiFe]-hydrogenases, including a putative membrane-bound group 4g [NiFe]-hydrogenase, and might use electrons harnessed from small organic acids/simple carbohydrates to produce H_2_. While speculative in the absence of experimental data, it may be hypothesised that the hydrogen could support the growth of a H_2_-ultilising archaeon, perhaps, in return for essential cellular building blocks such as lipids and vitamins (Ver Eecke et al. 2012; Topçuoğlu et al. 2016; Imachi et al. 2020; Yu et al. 2024).

Likewise, it is interesting to note that our ancestral reconstructions infer a complex ancestor of Asgard archaea as was observed previously (Eme et al. 2023). Further, our analyses predict that the Asgard archaeal ancestor has encoded a Wood-Lundgal pathway (WLP) (**Figure 5, Supplementary Data 13**), i.e. a pathway that can serve as electron sink supporting both autotrophic and heterotrophic growth modes (Ragsdale and Pierce 2008; Schuchmann and Müller 2014; Schuchmann and Müller 2016). However, this pathway seems to have been lost in various members of the Asgard archaea (Spang et al. 2019; Liu et al. 2021) (**Supplementary Data 10-13**); specifically, our analyses indicate that genes for proteins involved in the WLP were lost on the branch leading to the Njordarchaeia after their split from Heimdall- and Wukongarchaeia (**Figure 5**). The absence of the WLP as electron sink may have strengthened the dependency of members of this group on syntrophic H_2_-ultilising partners, such as Thermoproteota (Huber et al. 2000; Sakai, Nakamura, et al. 2022; Leung et al. 2024) that could serve as external electron sinks. It is tempting to speculate that such a syntrophic lifestyle may have permitted the loss of genes involved in various biosynthetic pathways in the ancestor of the *Panguiarchaeum* clade (**Figure 4,5**), increasing the dependency of its members on partner organisms.

This scenario on the evolution of a potentially obligate symbiont lineage among the Asgard archaea points to interesting parallels with the evolutionary trajectory leading to the origin of eukaryotes. Many current models on eukaryogenesis hypothesise syntrophy as a key driver for the establishment of an intricate symbiotic relationship between the archaeal ancestor of eukaryotes (Martin and Müller 1998; Spang et al. 2019; Imachi et al. 2020; López-García and Moreira 2020; Vosseberg et al. 2024), i.e. a likely sister lineage of the Hodarchaeales (Williams et al. 2020), and the alphaproteobacterial partner that evolved into the mitochondrion (Lane and Martin 2010; Poole and Gribaldo 2014; Ettema 2016; Martijn et al. 2018; Muñoz-Gómez et al. 2022). Similar to *Panguiarchaeum*, the eukaryotic host lineage also appears to have lost ancestral archaeal genes during eukaryogenesis, i.e. the transition from the First to the Last Eukaryotic Common Ancestor. For example, glycolysis and iron-sulphur cluster biosynthesis in eukaryotes are implemented by pathways with patchwork ancestries that include both archaeal, alphaproteobacterial and other bacterial contributions (Freibert et al. 2017), while the archaeal TCA cycle was replaced by that of the mitochondrion in eukaryotes (Santana-Molina et al. 2025). Similarly, the evolution of *Panguiarchaeum* was shaped by the loss of genes involved in the TCA cycle and several biosynthetic pathways. However, reductive evolution in *Panguiarchaeum* has also led to the loss of several genes encoding ESPs found in its close relatives. In contrast, the eukaryotic lineage expanded an already large protein repertoire and its ESPs via gene duplication and genome expansion, as well as acquisitions from mitochondrial and other bacterial genes along its stem (Pittis and Gabaldón 2016; Eme et al. 2017; Tria et al. 2021; Vosseberg et al. 2021; Vosseberg et al. 2024). Recent analyses have demonstrated that Asgard archaeal genomes often contain substantial proportions of horizontally-acquired genes of bacterial origin (Wu et al. 2022), and we hypothesise that part of the difference in evolutionary trajectories of *Panguiarchaeum* and eukaryotes may result from the nature of the symbiotic partner, either archaeal (*Panguiarchaeum*) or bacterial (eukaryotes) as well as environmental parameters (e.g. temperature). To address these hypotheses, prospective efforts would benefit from a better sampling of symbiotic members of the Asgard archaea including both syntrophic and/or genome-reduced representatives, combined with their co-cultivation with partner organisms and physiological characterisation.

## Materials and Methods

### Marker gene inspection and ranking analysis

#### Marker gene inspection

To accurately place Njordarchaeales and Panguiarchaeals in the archaeal tree of life, we first downloaded the NM57 A64 dataset from Eme et al., 2023 (Eme et al. 2023) and assigned a cluster of orthologs (COG) family to each sequence using hmmsearch (Finn et al. 2011) v3.3.2 (settings: -E 1e-5). The best hit for each protein sequence was selected based on the lowest e-value and highest score, and the COG families representing the majority of sequences in each marker gene were selected to annotate each marker. These COG families were then compared to 185 marker gene families used in four other previous studies (Williams et al. 2020; Moody et al. 2022) to create a non-redundant marker set comprising 112 unique COG families for further inspection. Potential orthologous sequences assigned to these 112 unique families were identified in 966 archaeal and 1325 bacterial genomes/MAGs using hmmsearch v3.3.2 (settings: -E 1e-5) and used for subsequent phylogenetic analyses. Since gene fission of DNA-directed RNA polymerase subunits A and B were reported in some archaea (Werner 2007), we used a custom HMM profile of COG0085 and COG0086 to detect “split” subunits and concatenate these subunits before alignment (https://doi.org/10.5281/zenodo.15477107).

We then aligned sequences assigned to these 112 marker gene families using mafft v7.453 (Katoh and Standley 2013) and removed poorly aligned amino acid sites with BMGE 1.12 (Criscuolo and Gribaldo 2010) (settings: -m BLOSUM30 -h 0.55). The maximum likelihood trees for these marker genes were inferred using IQ-tree v2.1.2 (Minh et al. 2020) (-m LG+G -B 1000). We decorated the tips of these marker gene trees with annotations from KEGG orthology (Aramaki et al. 2020), PFAM (Bateman et al. 2004), COG (Galperin et al. 2021) and archeal COG (arCOG) (Makarova et al. 2015) information. We manually inspected all single protein phylogenies to identify marker gene families that did not meet archaeal monophyly and excluded those families from subsequent analyses. Furthermore, we identified and removed paralogues and sequences that experienced horizontal gene transfer (HGT) events from bacteria to archaea. In total, this yielded 100 COG marker gene families.

We performed a second round of inspection and curation based on corresponding arCOG families of these 100 COG families except COG0085 and COG0086 (See above). In this round, we only included the marker gene families if homologs were present in at least 60% of archaeal taxa and were duplicated in less than 20% of the genomes of major archaeal lineages (e.g., Asgard archaea, Thermoproteota and Halobacteriota), which resulted in 90 unique families with arCOG identifiers. Sequences were realigned with mafft-linsi v7.453, and poorly aligned sites were removed using BMGE as described above. The initial ML trees were inferred using IQ-tree v 2.1.2 (Minh et al. 2020) (-m LG+G -B 1000). These trees were then used as guide tree to perform phylogenetic inference based on the best-fitting model (Kalyaanamoorthy et al. 2017) (settings: -mset LG -madd LG+C10,LG+C20,LG+C30,LG+C40,LG+C50,LG+C60,LG+C10+R+F,LG+C20+R+F, LG+C30+R+F,LG+C40+R+F,LG+C50+R+F,LG+C60+R+F --score-diff all) with 1000 ultrafast bootstraps. We inspected the single gene trees to identify and remove sequences which seem to represent paralogues and/or were affected by long-branch attraction (LBA) artefacts or HGT.

#### Ranking analysis

We used a previously developed marker gene ranking procedure to rank each of the 90 markers based on the extent to which they recovered established archaeal phylum- or order-level lineages (Dombrowski et al. 2020; Moody et al. 2022). In brief, the number of splits (i.e., the occurrence of a certain taxon failing to group within its expected taxonomic clade, **Supplementary Data 16**) was counted across all bootstrap trees for each marker gene. We then defined the highest ranking (top 50%, 43 markers) marker sets based on the number of splits per phylogenetic cluster and the total number of splits normalised by the number of genomes within each tree.

### Concatenation, site filtration and species tree inference

#### Concatenated gene trees using taxon set 1: i.e. 966 taxa

The top 50% marker gene sequences were aligned with mafft-linsi v7.453 (Katoh and Standley 2013) and poorly aligned amino acid sites were removed with BMGE 1.12 (Criscuolo and Gribaldo 2010) (settings: -m BLOSUM30 -h 0.55). Genomes with more than 50% gaps in the alignment were further removed. The trimmed sequences of these 45 markers were then subsequently concatenated into a supermatrix comprising 966 taxa. Two additional supermatrixes were constructed by either excluding DNA topoisomerase VI subunit A (arCOG04143) alone or together with DNA topoisomerase VI subunit B (arCOG01165) for the following reasons (See also **Supplementary Information 1.1**):

1. The mixing of orthologues of arCOG04143 provides spurious support for the relationship between Njordarchaeia and Asgard archaea.
2. The interaction between DNA topoisomerase VI subunits might lead to co-evolution of some amino acid residues.

Concatenated gene trees both with and without DNA topoisomerase VI subunits were inferred using IQ-Tree v2.1.2 (Minh et al. 2020) under LG+C60+G+F model with posterior mean sites frequency approximation (Wang et al. 2018) with guide trees inferred under LG+G+F model.

#### Concatenated gene trees using taxon set 2: i.e. 303 taxa

To resolve the relationship between Njordarchaeales, Panguiarchaeales, Korarchaeota and Asgard archaea, we next excluded divergent sequences from DPANN MAGs/genomes and downsampled the Thermoproteota (previously TACK), and Halobacteriota, Thermoplasmatota, Hydrothermarchaeota, Methanobacteriota, Methanobacteriota_A and Methanobacteriota_B lineages (previously “Euryarchaea”) based on the predicted genome completeness and contamination (completeness-5 X contamination) (Parks et al. 2017) to create a dataset of 303 taxa by one-per-Thermoproteota/Asgardarchaeota-family and one-per-”Euryarchaea”-order citeria. Marker gene sequences were realigned with mafft-linsi v7.453 (Katoh and Standley 2013) and trimmed with BMGE 1.12 (Criscuolo and Gribaldo 2010) (settings: -m BLOSUM30 -h 0.55). We concatenated three supermatrices by including all top 50% ranked markers and excluding arCOG04143 alone or with arCOG01165. Trees were inferred under the LG+G+F and LG+C60+G+F model, as well as LG+EDM0256LCLR+G+F model using IQ-Tree v2.1.2 (Minh et al. 2020) and IQ-Tree v2.2.2.7 (Minh et al. 2020) respectively (settings: -B 1000 -alrt 1000, see also 71 taxa). Focusing on the supermatrix of 303 archaeal taxa, excluding DNA topoisomerase subunits, we progressively removed 10% to 50% of fastest evolving sites according to the empirical Bayesian rate under LG+C60+G+F model (settings –rate), and 5%, 10% to 50% most heterogeneous sites using Alignment_pruner.pl (https://github.com/novigit/davinciCode/blob/master/perl). Trees were inferred under either LG+C60+G+F or LG+EDM0256LCLR+G+F models (settings: -B 1000 -alrt 1000).

#### Concatenated gene trees using taxon set 3: i.e. 71 taxa

A smaller dataset of 71 taxa was created based on the 303 taxa set (set 2) for Bayesian phylogenetic analyses. In particular, we kept one genome of “Euryarchaea” for each GTDB R207 class as an outgroup and one genome of thermoproteotal (TACK) lineages for each GTDB R207 order considering predicted genome completeness and contamination (completeness-5xcontamniation) (Parks et al. 2017). Marker gene sequences were realigned with mafft-linsi v7.453 (Katoh and Standley 2013) and trimmed with BMGE 1.12 (Criscuolo and Gribaldo 2010) (settings: -m BLOSUM30 -h 0.55). We concatenated only one supermatrix without both DNA topoisomerase subunits (arCOG04143 and arCOG01165). ML trees were inferred under LG+G+F and LG+C60+G+F model using IQ-Tree v2.1.2 (Minh et al. 2020) (settings: -B 1000 -alrt 1000).

Additionally, we performed a comprehensive phylogenetic analysis using a recently developed CAT-PMSF pipeline (Szánthó et al. 2023). In particular, we used three different guide trees representing different hypotheses regarding the relationship of Njordarchaeia to Korarchaeota and Asgard archaea, namely, Korarchaeota topology: Njordarchaeia-sister-to-Korarchaeota; Asgard archaea topology: Njordarchaeia-sister-to-Wukong-Heimdallarchaeia and Asgard archaea sister topology: Njordarchaeia-sister-to-Asgard-Archaea to test if used guide trees affect topology inference. We then used these 3 different starting topologies in a subsequent bayesian analysis with the CAT model in combination with either Poisson or LG exchangeabilities and a 4 discrete categories gamma rate model. We ran four different Markov chains for each topology representing different hypotheses under the Poisson+CAT+G model in Phylobayes v 1.8 (Lartillot et al. 2013). Additionally, four different Markov chains were run for both Asgard archaea and Korarchaeota topology under the LG+CAT+G model. These chains were run until the effective sample size of all parameters was above 100 and/or the relative divergence of all parameters was below 0.3, or visual inspection with Tracer 1.7 (Rambaut et al. 2018) indicated convergence (**Supplementary Data 17**). Next, we extracted posterior mean site-specific stationary distributions of amino acids for each chain using readpb_mpi (Lartillot et al. 2013), which were converted to site-specific frequency profile using the python script convert-site-dist.py (https://github.com/drenal/cat-pmsf-paper/blob/main/scripts/convert-site-dists.py) (Szánthó et al. 2023). The site-specific frequency profile was then used to infer ML trees using IQ-Tree v2.1.2 (settings: - fs .sitefreq -B 1000 -alrt 1000).

We next applied the EDCluster algorithm (Schrempf et al. 2020) on one of the LG+CAT+G4 chains that yielded the highest likelihood under LG+CAT-PMSF model to infer EDM with various components (8, 16, 32, 64, 128 and 256) using log centred log-ratio transformation (LCLR). ML trees were then inferred under different components of EDM models using IQ-Tree v2.2.2.7 (e.g., settings: -m LG+EDM0256LCLR+G+F -B 1000 -alrt 1000) and universal distribution mixture models (UDM) with 128, 256 and 512 components (Schrempf et al. 2020). The model fit between Poisson+CAT-PMSF, LG+CAT-PMSF, LG+G+F and LG+C60+G+F were assessed using a recently developed parametric bootstrap method (Giacomelli et al. 2025), which compares the model adequacy in describing across-site compositional heterogeneity.

### Compositional constrained analyses

Constrained searches were performed for Asgard archaea (-sister) and Korarchaeota topology based on the unconstrained ML tree under the same model using IQ-Tree v2.1.2 (Minh et al. 2020) via the -g and -wslr flags. AU tests were performed using consel v 1.20 (Shimodaira 2001) with 10000 bootstrap replicates. Site-wise log-likelihood differences were calculated for Asgard archaea and Korarchaeota topology under LG+G+F, LG+C60+G+F, Poisson+CAT-PMSF and LG+CAT-PMSF models to investigate the attraction of Njordarchaeia towards Korarchaeota. The effective amino acids (K_eff_) values of mixture models and site-specific frequency profile were calculated using the convert-site-dists-to-k_eff.py script (Schrempf et al. 2020; Szánthó et al. 2023). The site-wise log-likelihood differences were then grouped into different bins according to partitions, evolving rates and the effective amino acids.

### RED values scaling and classification of ranks

Relative evolutionary distance (RED) metric on the species trees was calculated with methods described in the PhyloRank v1.1.12 tool (Parks et al. 2018). Class, family and order rank nodes in the 303 archaeal sets were extracted from our phylogenetic tree based on LG+EDM0256LCLR+G+F model, as well as, from GTDB R220 reference tree (Rinke et al. 2021). Linear regression analysis of the RED values of our phylogenetic tree and GTDB R220 tree was performed using the Scipy package. The taxonomic boundaries used in GTDB R220 were scaled onto our tree by the fitted slope and intercept.

### Amino acid composition

We used a custom Python script to estimate the relative frequency of each amino acid in our 71 taxa dataset for the concatenation without DNA topoisomerase subunits (https://doi.org/10.5281/zenodo.15477107). A principal component analysis (PCA) was performed using Scikit-learn (Pedregosa et al. 2011), and two axes were plotted along with eigenvectors using Seaborn (Waskom 2021). We identified RYPEI(WKVL)/QSNT(DGC) as thermophilic enriched/depleted amino acid residues in the concatenation using a binomial test (Baker et al. 2024) and modified the GFmix (Muñoz-Gómez et al. 2022) model to account for the variation of the ratio RYPEI(WKVL)/QSNT(DGC) at every branch. The likelihood of different tree topologies under the GFmix-RYPEI(WKVL)/QSNT(DGC) model was then calculated with LG+C60+G+F and LG+EDM0256LCLR+G+F, with weights of mixtures model, branch length, and alpha shape parameters estimated using IQ-Tree v2.1.2 (Minh et al. 2020).

### Gene calling and Annotations

We annotated 966 archaeal genomes/MAGs using an in-house annotation pipeline (https://doi.org/10.5281/zenodo.15477107). In brief: Gene calling was performed using Prokka (Seemann 2014) (v1.14.6, settings: –kingdom Archaea –addgenes – increment 10 –compliant –centre UU –norrna –notrna). The generated protein files were searched against COG database (Galperin et al. 2021) (NCBI_COGs_Oct2020.hmm), arCOG database (Galperin et al. 2021) (All_Arcogs_2018.hmm), PFAM (Bateman et al. 2004) (Release 34.0), TIGRFAM (Release 15.0), KEGG orthology (Aramaki et al. 2020) (KO) profile (downloaded April, 2019), the Carbohydrate-Active enZymes (CAZy) database (Lombard et al. 2014) (downloaded from dbCAN2 in September 2019), the Transporter Classification Database (Saier et al. 2021) (TCDB; downloaded in November 2018), the hydrogenase database (Søndergaard et al. 2016) (HydDB; downloaded in November 2018), the MERPOS database (Rawlings et al. 2018) (Release 12.4) and NCBI_nonredudant database (NCBI_nr; downloaded in November 2018). Additionally, we used Interproscan version 5.61.93.0 to scan for protein domains (Jones et al. 2014) (settings: --iprlookup --goterms).

COG (settings: -E 1e-5), arCOG (settings: -E 1e-5), PFAM (settings: -E 1e-5), TIGRFAM (settings: -E 1e-5), KO (settings: -E 1e-5), and CAZy (settings: -E 1e-10) identifiers were assigned using hmmsearch v3.3.2. The TCDB, HydDB and MERPOS database were searched using BLASTp v2.12.0 (Altschul et al. 1997) (settings: -outfmt 6, -evalue 1e-20), and NCBI_nr database was searched using DIAMOND v2.0.6 (Buchfink et al. 2015) (settings: –more-sensitive –e-value 1e-5 –seq 50 –no-self-hits – taxonmap prot.accession2taxid.gz). Asgard COG (Liu et al. 2021) (asCOG) database was searched with psi-blast v2.12.0 (settings: -show_gis -outfmt 6 -dbsize 100000000 -comp_based_stats F -seg no -evalue 1e-20). The best hit of each protein for all other database searches was selected based on the lowest e-value and highest score, and results were summarised for Njordarchaeales MAGs. Additionally, 1325 bacterial genomes/MAGs and 137 eukaryotic genomes and/or largely complete transcriptomes (Santana-Molina et al. 2025) (**Supplementary Data 18**) were searched against COG, arCOG, PFAM and KO databases using the same pipeline. Njordarchaeia MAGs annotation results were summarised in **Supplementary Data 7-11 and 19-21**.

### Metabolic comparison

Metabolic comparisons were based on the annotation results described above. We counted the occurrence of each gene across each MAG/genome and reported the presence and absence pattern using Pandas (The pandas development team 2024) in Python v3.9.0 (Van Rossum and Drake 2009). This data was then summarised per GTDB R207 class level whenever possible. The Occurrence of ESPs was compared using the results from asCOG searches. The presence and absence pattern of genes across each database (COG_DB and ARCOG_DB) was analysed with PCA and t-distributed Stochastic Neighbor Embedding (t-SNE) using scikit-learn (Pedregosa et al. 2011).

### Contamination assessment of representative Njordarchaeia MAGs and its impact on phylogenetic analyses

Metagenomes of the representative MAGs and similar samples were downloaded from public database (Dombrowski et al. 2018; Qu et al. 2023). These metagenomes include samples from hydrothermal vents in Guaymas Basin (Dombrowski et al. 2018) and a hot spring in Yunan, China (Qu et al. 2023). We used the bowtie2 v2.5.3 (Langmead and Salzberg 2012) to map reads to the MAGs, and samtools v1.18 to process the mapped profiles. We used Anvi’o v8 (Eren et al. 2021) to perform hierarchical clustering based on sequence composition and coverage patterns across different metagenomes. Clusters were identified based on the similarity of coverage profile across different samples while taking into account sample similarity. Marker genes used in our phylogenetic analyses and homologs from marker sets in a recent study (Zhang et al. 2025) were mapped back to the contigs using sequences selected for phylogenetic analyses and annotations described above.

### Single gene tree inferences for ancestral reconstruction

The predicted protein sequence of 303 archaeal genomes/MAGs was searched against the arCOG database using hmmsearch v3.3.2 (Finn et al. 2011) (settings: -- tblout –domtblout –notextw). We used the best hits (the lowest evalue and highest bitscore, cutoff: e-value < 1E-3) to identify and split potential fused proteins. Specifically, we first subtracted the position of the first domain hit from the full protein using bedtools subtract (Quinlan and Hall 2010) (v2.26.0). Secondly, we investigated the presence of a secondary domain hit that is assigned to a different arCOGs than the first domain. Subsequently, we repeated these two steps until all the proteins were investigated for up to four domains. Finally, the individual domains from the original protein were extracted using the positional domain information from hmmsearch output using bedtools getfasta function. To keep track of unsplit and split domains from the original protein, we assigned ‘a0’ notation for unsplit proteins and ‘a1’ to ‘a4’ for split domains (primarily to quarterly) in the generated trees.

We then combined all protein sequences and removed sequences with ambiguous amino acids using a custom Python script (remove_seq_with_specific_char.py, https://doi.org/10.5281/zenodo.15477107). To improve the phylogenetic signal in the single gene trees, we carefully performed three rounds of alignments before the final phylogenetic inference. First, we aligned sequences using mafft or mafft-linsi, for sequences with greater than or less than 1000 sequences, respectively. Alignments were then trimmed using trimAl 1.2rev59 (Capella-Gutiérrez et al. 2009) with – gappyout flag. Trimmed sequences with greater than or equal to 50% gaps were removed using a custom Python script (faa_drop.py, https://doi.org/10.5281/zenodo.15477107). Second, we realigned the arCOG gene families (with more than or equal to 4 sequences) using MAFFT as described above and trimmed the alignment with BMGE (v1.12; settings: -m BLOSUM30 -b 2 -h 0.55) (Criscuolo and Gribaldo 2010). We then inferred a phylogenetic tree using IQ-Tree v2.1.2 (settings: -m LG+G) (Minh et al. 2020). To alleviate the effect of long-branching sequences on phylogenetic inferences, we identified sequences/clusters of sequences on long branches of the inferred tree using a custom script (cut_gene_tree.py, https://doi.org/10.5281/zenodo.15477107) with a branch cutoff of two (Davín et al. 2025). Single sequences were discarded, and clusters with more than four sequences were separated into different gene families. We appended the notation “_ [0-3]” to keep track of the separation of sequences into a new cluster. Finally, we aligned and trimmed the sequences as described in the second step. For each family, we selected the best-fit model using the model test with IQ-Tree v2.1.2 (settings: -m MF -mset LG -madd LG+C10, LG+C10+G, LG+C10+R, LG+C10+F, LG+C10+R+F, LG+C10+G+F, LG+C20, LG+C20+G, LG+C20+F, LG+C20+G+F, LG+C20+R, LG+C20+R+F, LG+C30, LG+C30+G, LG+C30+R, LG+C30+F, LG+C30+R+F, LG+C30+G+F --score-diff all -T 2). For the selected best-fitting model with profile mixtures, we used the posterior mean site frequency (PMSF) approximation (Wang et al. 2018) to infer the bootstrap tree distribution (settings: -s trimmed_aln.faa -m BEST-FITTING MODEL -T 2 -wbtl -B 1000 -pers 0.2 -nstop 500) using a guide tree inferred from LG+G model (settings: -T 2) (Minh et al. 2020). For the best-fitting non-mixture models, bootstrap tree distribution was inferred using IQ-Tree v2.1.2 with the settings: -T 1 -wbtl -B 1000 -pers 0.2 -nstop 500. In total, we generated 6077 single gene tree distributions in the analysis for ancestral reconciliations.

### Gene tree and species tree reconciliations

We used the amalgamated likelihood estimation (ALE, v1.0) approach to reconcile single gene trees against the species tree. First, we generated the ALE objects from the bootstrap trees that approximate tree uncertainty using ALEobserve. Using genome completeness estimation from CheckM v2 (Chklovski et al. 2023) and ALEml_undated (Szöllősi et al. 2015) program, we first reconciled the gene trees against species tree using the default parameters, which assumes origination at any node of the species tree was uniform. Based on the reconciliation results from this, we next used an additional reconciliation approach described by Coleman et al., 2021 (Coleman et al. 2021) that assumes the origination probabilities at the root of the species tree (O_R) are different to those for all other internal nodes. We first inferred the O_R for each of the 21 arCOG functional categories by maximising the total reconciliation likelihood over all gene families in that category using two Python scripts (setup_OR_estimation.py and O_R_optimisation.py, https://doi.org/10.5281/zenodo.15477107). Subsequently, we used the category-specific probabilities for origination at the root to reconcile the single gene trees and species trees and estimated the presence probabilities/copies of each family in the internal nodes of the tree. The presence probabilities of both approaches for each family for the nodes of interest are provided in **Supplementary Data 13**. Ancestral genome size was predicted using locally weighted smoothing regression based on the relationship between the genome size of extant taxa and their arCOG gene family number. The uncertainty of the prediction of genome size was taken into account using the bootstrap resampling approach (1000 times).

### Phylogenetic analyses of DNA topoisomerase subunit A and DNA topoisomerase B

The homologs of DNA topoisomerase subunit A and B, i.e. COG1389 and COG1697, were identified and retrieved from 966 archaeal, 1325 bacterial and 137 eukaryotic genomes/MAGs or largely complete transcriptomes. Sequences of each gene family were first aligned with mafft-linsi v7.453, and poorly aligned alignment sites were trimmed with BMGE v1.2 (Criscuolo and Gribaldo 2010) (Settings: -m BLOSUM30 -h 0.55). The inference of phylogenetic trees for individual maker gene families was conducted as follows: an initial ML tree was inferred under LG+G with IQ-Tree v 2.1.2 (Minh et al. 2020). Single gene trees were manually inspected to remove putative LBA artefacts, and the remaining sequences for each gene were realigned and trimmed as described above; finally, an ML tree was inferred for each gene under LG+C60+G+F+PMSF (Wang et al. 2018) model (settings: -B 1000 -alrt 1000) using guide tree inferred from LG+F+G.

### Phylogenetic analyses of Actin homologs

The actin homologs were identified from 966 archaeal, 1325 bacterial and 137 eukaryotic genomes/MAGs or largely complete transcriptomes using IPR00400. The cell shape determining protein MreB homologs, a distant homolog to actin, in 966 archaeal genomes were identified using COG1077 and dereplicated using cd-hit v4.7 (-c 0.75) (Fu et al. 2012). These sequences were combined and aligned with mafft-linsi v7.453 (Katoh and Standley 2013). Poorly aligned sequences were trimmed using BMGE (Criscuolo and Gribaldo 2010) (settings: -m BLOSUM30 -h 0.55). An initial ML tree was inferred under LG+G using IQ-Tree v 2.1.2 (Minh et al. 2020), and sequences showing potential instances of LBA artefacts were removed; the remaining sequences were then realigned and trimmed as described above; finally, an ML tree was then inferred under LG+C60+G+F+PMSF model (settings: -B 1000 -alrt 1000) using a guide tree inferred from LG+F+G. 20% of the most heterogeneous sites were removed using alignment_pruner.pl and an ML tree was inferred in the same way under LG+C60+G+F+PMSF using a guide tree inferred from LG+F+G

### Phylogenetic analyses of ribosomal protein L28

The ribosomal protein L28/Mak16 homologs (i.e. PF01778) were retrieved from 966 archaeal, 1325 bacterial and 137 eukaryotic genomes/MAGs or largely complete transcriptomes (evalue: 0.0001). Sequences were first aligned with mafft-linsi v7.453, and gappy columns were removed using trimal 1.2rev59 (Capella-Gutiérrez et al. 2009) with -gappyout flag. An initial ML tree was inferred under LG+G using IQ-Tree v2.1.2 (Minh et al. 2020), and sequences showing potential instances of LBA artefacts were removed; remaining sequences were then realigned and trimmed as described above; finally, an ML tree was inferred under LG+C60+G+F+PMSF model (settings: -B 1000 -alrt 1000) using the guide tree inferred from LG+G.

### Phylogenetic analyses of oligosaccharyltransferase 3/6 like (OST3/6 like) and translocon-associated protein (TRAP) complexes subunit beta

The OST3/6-like homologs and TRAP beta were retrieved from 966 archaeal, 1325 bacterial and 137 eukaryotic genomes/MAGs or largely complete transcriptomes using PF04756 and PF05753 respectively, from the annotation results described above. These sequences were dereplicated using cd-hit v4.7 (Fu et al. 2012) (-c 0.7). Sequences of each gene family were first aligned with mafft-linsi v7.453, and gappy columns were removed using trimal 1.2rev59 (Capella-Gutiérrez et al. 2009) with - gappyout flag. An initial ML tree was inferred under LG+G using IQ-Tree v2.1.2 (Minh et al. 2020) for each gene family, and sequences showing potential instances of LBA artefacts were removed; remaining sequences for each gene family were then realigned and trimmed as described above; finally an ML tree was then inferred for each gene under LG+C60+G+F+PMSF model (settings: -B 1000 -alrt 1000) using guide tree inferred from LG+G.

### Phylogenetic analyses of the Snf7 family

The Snf7 family homologs were identified and retrieved from 966 archaeal and 137 eukaryotic genomes/MAGs based on the presence of the domain IPR005024. Eukaryotic sequences were dereplicated using cd-hit v4.7 (Capella-Gutiérrez et al. 2009) (-c 0.7). We created two datasets for this family, i.e. a dataset with only archaeal sequences and another with archaeal and eukaryotic sequences. Sequences of each dataset were first aligned with mafft-linsi v7.453 (Katoh and Standley 2013), and poorly aligned sites were removed using BMGE v1.1.2 (Criscuolo and Gribaldo 2010) (settings: -m BLOSUM30 -h 0.55). The inference of phylogenetic trees was conducted as follows: an initial ML tree was inferred under LG+G using IQ-Tree v 2.1.2 (Minh et al. 2020), and sequences showing potential instances of LBA artefacts were removed; remaining sequences were then realigned and trimmed as described above; finally, an ML tree was then inferred funder the LG+C60+G+F+PMSF model (settings: -B 1000 -alrt 1000) using a guide tree inferred from LG+G. We also removed the 20% most heterogeneous sites from the alignment with both archaeal and eukaryotic sequences included using Alignment_pruner.pl (https://github.com/novigit/davinciCode/blob/master/perl) and inferred an ML tree using the same approach.

### Phylogenetic analyses of CDP-archaeol synthase (CarS), archaetidylinositol phosphate synthase (PgsA) and phosphatidylserine synthase (PssA)

Sequences identified as CarS (K19664), PgsA (K17884), and PssA (K17103) were retrieved from 966 archaeal, 1325 bacterial and 137 eukaryotic genomes/MAGs or largely complete transcriptomes, respectively, from the annotation results described above. Sequences of each gene family were first aligned with mafft-linsi v7.453 (Katoh and Standley 2013), and gappy columns were removed using trimal 1.2rev59 (Capella-Gutiérrez et al. 2009) with the -gappyout flag. The inference of phylogenetic tree for individual gene families was conducted as follows: an initial ML tree was inferred under LG+G using IQ-Tree v2.1.2 (Minh et al. 2020) for each gene family, and sequences showing potential instance of LBA artefacts were removed; remaining sequences for each gene were then realigned and trimmed as described above; finally an ML tree was then inferred for each gene under LG+C60+G+F+PMSF model (settings: -B 1000 -alrt 1000) using guide tree inferred from LG+G.

### Phylogenetic analyses of [NiFe] group hydrogenase, large subunit

Backbone sequences for phylogenetic analyses of [NiFe] group 3 and 4 hydrogenases were obtained from a previous study (Spang et al. 2019). Njordarchaeia homologs were added to the backbone sequences and aligned with mafft-linsi v7.453. Poorly aligned sites were removed using BMGE v1.1.2 (Criscuolo and Gribaldo 2010)(settings: -m BLOSUM30 -h 0.55). A phylogenetic tree was inferred under the LG+C60+G+F+PMSF model guide trees inferred from the LG+G+F model using IQ-Tree v2.1.2 (Minh et al. 2020) (settings: -B 1000 -alrt 1000). Phylogenetic tree was visualised using ggtree 3.2.0 (Xu et al. 2022).

### Network inferences

The input dataset for network prediction was obtained from OTU community profiles derived from 248,559 NCBI metagenomes (Woodcroft et al. 2024) from the ‘Sandpiper’ website (https://sandpiper.qut.edu.au), which were annotated with the GTDB R214 taxonomy (Parks et al. 2022). First, we used the known association between *Huberarchaeum* and *Altiarchaeum* to confirm that our network association approach is able to detect symbiotic interactions. We selected all samples that contain taxa annotated as *Huberarchaeum* (67 samples), respectively. We transformed the coverage of the taxa in these samples to counts by multiplying with 100 and formatted the data into a table listing files in rows and all OTUs in columns. To reduce the scarcity of the dataset, we filtered for OTUs present in 10%, 20%, 30%, 40%, 50%, 60%, 70%, 80%, 90% of the samples. To estimate the minimum number of samples required for robust, true positive prediction of *Huberarchaeum*-*Altiarchaeum* associations, we drew random samples and applied two different filter criteria. Numbers of 10, 20, 30, 40, 50, and 60 samples were drawn; the random data sets were filtered for OTUs detected in at least 50% and 80% of all random samples, respectively. For each condition, ten networks were inferred. Statistics were run across all ten networks within one condition. These networks were inferred using SpiecEasi v1.1.3 (Kurtz et al. 2015). Across all networks, we checked for associations between *Huberarchaeum* and *Altiarchaeum*. In all networks inferred (>30 samples), the expected association was robustly detected. However, below a number of 30 samples, the association was detected only in 50% of the samples. This clearly indicates a minimum number of > 30 samples (equaling 40% of samples) that is necessary for reliable inference of association. We also attempted to confirm the known association between *Nanoarchaeum* and its host *Ignicoccus* (order Sulfolobales). Our dataset contained several OTUs, assigned to the genus *Nanoarchaeum* (otu_658, otu_3077) and the genus *Ignicoccus* (otu_3007, otu_678, otu_3062, otu_3063, otu_3922, otu_4367). However, some of the taxa were present in only very few samples (**Supplementary Figure 35,36)**, and subsequently in the inferred network, no association between any of these taxa was predicted. Interestingly, some OTUs showed strong, significant negative correlations (**Supplementary Figure 35,36**), suggesting a high specificity of the symbiotic association on OTU level, although larger datasets are needed to confirm this assumption.

Next, we selected all samples that contain taxa annotated as “*Panguiarchaeaceae”* in GTDB R220 (Rinke et al. 2021; Parks et al. 2022), a taxonomic rank that is equivalent to the order Njordarchaeales proposed in this manuscript (**Figure 1BC**), resulting in a subset of 120 samples. To reduce the scarcity of the dataset, we filtered the dataset before network inference. We used 5 different occupancy thresholds based on shoulder theory and commonly applied criteria to prepare 5 different datasets from which separate networks were inferred. OTUs had to be present in (1) at least 24 samples as well as in at least (2) 40%, (3) 50%, (4) 60%, and (5) 70% of the entire dataset to be included in the network. Across all networks, we checked for associations involving OTUs of our target lineage in a comparative manner. To exclude random associations involving our target lineage, we prepared random networks and selected only those associations that were unlikely to be found randomly. For each filter criterion, 999 random networks were drawn based on the original network inferred from the filtered data. Then we compared the taxonomic distribution of OTUs associated with OTUs annotated as Njordarchaeales (equivalent to “*Panguiarchaeaceae”* in GTDB R220) between the random networks and the original network. Association frequencies were compared based on phylum, class, order, and family level. Next, we explored associations at the genus level. A single OTU within the genus *Panguiarchaeum* was detected in higher numbers across samples, that is OTU_804 (43 samples), for which we investigated association partners. All analyses were performed in R v4.4.1 (R Core Team R 2024).

## Data availability

All genomic data/transcriptomic data analysed in this study are available at NCBI (Supplementary Data 16, 18) and are deposited in the data repository (https://doi.org/10.5281/zenodo.15477107). Data generated in this study including single gene and concatenated phylogenies (i.e., sequence files, alignments, and treefiles) have also been deposited in our data repository at Zenodo (https://doi.org/10.5281/zenodo.15477107) under the following license CC BY 4.0.

## Code availability

Workflows for annotations, phylogenies, reconciliation, and custom scripts (bash/python) to analyse and parse annotation data, generate single gene trees and perform ancestral gene content reconstruction are deposited in the repository (https://doi.org/10.5281/zenodo.15477107).

## Acknowledgements

This project has received funding from the European Research Council (ERC) under the European Union’s Horizon 2020 research and innovation programme (grant agreement No. 947317, ASymbEL to A.S. and grant agreement No. 714774, GENECLOCKS to G.J.S.). Furthermore, this work was supported by the Gordon and Betty Moore Foundation (GBMF9741 to T.A.W., A.S., and G.J.S.) and the Simons Foundation with the Moore–Simons Project on the Origin of the Eukaryotic Cell, 735929LPI (https://doi.org/10.46714/735929LPI) (to A.S. and coworkers). Our research is funded by the John Templeton Foundation (63451 to L.Sz., G.J.Sz., T.A.W. and A.S.; the opinions expressed in this publication are those of the authors and do not necessarily reflect the views of the John Templeton Foundation. We also want to thank Tara Mahendrarajah for valuable feedback on the manuscript.

## Author contributions

W.H. T.A.W. and A.S. conceived the study; W.H. analysed and annotated genomes and generated accompanying data tables; W.H. and A.S. analysed genomic data; W.H. T.A.W. and A.S. performed and analyzed phylogenetic analyses, C.R. and M.P. performed co-proportionality analysis; W.H., Z.H., T.W., C.R., M.P., G.J.S, L.S, T.E., and A.S. interpreted data; W.H., T.W., and A.S wrote and all authors edited and approved the manuscript.

