## Supplementary material for "Phylogenomic analyses reveal that Panguiarchaeum is a clade of genome-reduced Asgard archaea within the Njordarchaeia"

### Table of Contents

**1. The placement of Njordarchaeia in archaeal phylogeny** **3**
1.1 Complex evolutionary history of DNA topoisomerase VI subunits A and B 3
1.2 Taxonomic rank considerations 4
1.3 Addressing site-heterogeneity with the CAT-PMSF approach 5
1.4 Conflicting signals in analyses under the LG+C60+G+F model are likely due to inadequate modelling of site compositional heterogeneity 6
**2. Occurrence and phylogeny of Eukaryotic signature proteins (ESPs)** **7**
2.1 Occurrence of ESPs 7
2.2 Phylogeny of ESPs 7
2.2.1 Ribosomal protein L28e/Mak16 homologs 7
2.2.2 Actin homologs 7
2.2.3 N-glycosylation (OST3/OST6 and TRAP Beta): 8
2.2.4 SNF7 proteins 8
**3. Metabolism** **9**
3.1 Informational processing and repair systems 9
3.1.1 DNA replication and repair 9
3.1.2 Evolution of reverse gyrase 9
3.1.2 Transcription 10
3.1.3 Translation 11
3.2 Metabolic features 11
3.2.1 Central carbon metabolism 11
3.2.2 Peptides/amino acids degradation 13
3.2.3 Redox balance and energy conservation 13
3.2.4 Purine and pyrimidine biosynthesis 14
3.2.5 Purine and pyrimidine salvage 15
3.2.6 Amino acid biosynthesis 15
3.2.7 Lipid biosynthesis 16
3.2.8 Vitamin and cofactor biosynthesis 18
**4 Positive controls in the network inference** **18**
**5. Supplementary Figures** **21-57**
**6. Supplementary References** **57**

### 1. The placement of Njordarchaeia in archaeal phylogeny

#### 1.1 Complex evolutionary history of DNA topoisomerase VI subunits A and B

DNA topoisomerase (topo) VI is a heterotetrameric type IIB topo formed by two Top6A and two Top6B subunits (McKie et al. 2022). It was initially isolated from Sulfolobus shibatae (Bergerat et al. 1997) and later identified in many archaea, some eukaryotes and some bacteria (Forterre et al. 2007). In contrast to archaea, eukaryotic cells possess, in addition, a topoVI-like complex which comprises two proteins, i.e. SPO11 and Top6B-like, homologous to Top6A and Topb6B subunits, respectively. This complex plays a crucial role in the initiation of meiosis by inducing double-strand breaks (DSB) to initiate homologous recombination and to establish conditions for chromosome segregation.

Our phylogenetic analyses of the Top6A/SPO11 family homologs from 966 archaeal, 1325 bacterial and 137 eukaryotic metagenome-assembled genomes (MAGs), genomes and largely complete transcriptomes revealed two distinct Top6A/SPO11 clades containing Asgard archaea. Among these two clusters, cluster 1 (K10878 clade) shared a close relationship to eukaryotic SPO11 homologs, with Heimdallarchaeia sequences branching basal to eukaryotic homologs (Supplementary Figures 2-3), which is compatible with the previously proposed Heimdallarchaeia ancestry of Eukaryotes (Eme et al. 2023). Njordarchaeia possessed only one homolog of Top6A clustering within Asgard archaea, supporting the Asgard archaea topology. Some members of Odinarchaeia, Jordarchaeia and Thorarchaeia, however, encode a paralog of Top6A, which branched at the base of a clade composed of thermoproteotal members. Korarchaeota encoded long-branching Top6A homologs, which formed a clade with several thorarchaeial homologs and are distinct from the K10878 homologs (Supplementary Figure 3).

Similarly, our phylogenetic analyses of Top6B family enzymes recovered Njordarchaeia together with Sifarchaeia basal to other Heimdallarchaeia and Eukaryotes, supporting the Asgard archaea topology (Supplementary Figure 4). We furthermore recovered a second asgard archaea cluster comprising divergent Top6B paralogs from Odinarchaeia, Jordarchaeia and Thorarchaeia.

Thus, the Top6A and 6B phylogenies,  seem to share a similar evolutionary history which, however, is not in full agreement with the species topology.  For example, in both Top6A and Top6B families, the homologs from Sulfolobales, Thermofiales, and Thermoproteales branched sister to a clade comprising Asgardarchaeaota and Eukaryote. By contrast, Top6A from Bathyarchaeia, Marsarchaeales and Geothermarchaeales branched within the second asgard archaea clade, i.e., K03166 (Supplementary Figures 2,3) albeit with low support, indicating a complicated evolutionary history of Top6A within the thermproteotal lineages. In turn, due to the presence of distinct paralogs in different asgard archaea lineages and complicated evolutionary history, Top6A subunits are not an ideal marker gene to address the species phylogeny and the relationship of Njordarchaeaia with Asgard archaea,

Korarchaeota and other thermoproteotal members. The reconstruction of species trees based on gene concatenations is based on the premise that individual gene families are single-copy markers and follow the same evolutionary history. Mixing of orthologues and paralogues can confound phylogenetic signals and skew phylogenetic results (see below). For example, because many Asgard archaea and some Korarchaeatoa encode two distinct paralogs of elongation factor 2 (EF2), with one branching closer to eukaryotes (Narrowe et al. 2018), this marker was previously excluded from concatenated datasets used to infer the archaeal phylogeny (Spang et al. 2015a; Da Cunha et al. 2017; Zaremba-Niedzwiedzka et al. 2017; Spang et al. 2018).

Given the complex gene history of DNA topoisomerase, we assessed the impact of Top6A (arCOG04143) and Topo6B (arCOG01165) on the species phylogeny inferred from the 50% highest ranking makers (Supplementary Data 4). Notably, our phylogenetic analyses indicated that *Panguiarchaeum* is a divergent clade within Njordarchaeia (Figure 1BC, Supplementary Figures 5-7). However, including different paralogs of Top6A (arCOG04143) supported different hypotheses for the placement of the Njordarchaeaia as a whole. To be more specific, including the K10878 paralogs of the Top6A (arCOG04143) in the concatenation leads to a recovery of Asgard archaea topology (Supplementary Figures 2,5, 7), while keeping the K03166 paralogs leads to a recovery of Korarchaeota topology based on the LG+C60+F+G+PMSF analyses on the expanded set 1 (Supplementary Figures 2,3,6). This confirms the confounding impact of mixing orthologues and paralogues in the phylogenetic analyses. In this regard, it is notable that a constraint analysis, based on the LG+C60+G+F analyses on the streamlined set 2 (see methods), revealed that DNA topoisomerase VI subunits A (arCOG04143) and B (arCOG01165) ranked as the first and third most influential markers in determining the relationship of Njordarchaeia with Asgard archaea (Supplementary Figure 8).

#### 1.2 Taxonomic rank considerations

“Njordarchaeota” were first proposed based on “GB_154”, a MAG referred to as *Candidatus* Njordarchaeum guaymaensis (Xie et al. 2022). This MAG was not included in the phylogenetic analyses due to the presence of a closely related (ANI: 98.39%) MAG, i.e. GCA_029856705.1, which had a slightly higher completeness based on CheckM (Parks et al. 2015). The GCA_029856705.1 MAG was placed consistently in the *Njordarchaeaceae* clade (Figure 1BC, Supplementary Figures 5-7, 10, 12-13). A subsequent study placed both *Njordarchaeaceae* and *Panguiarchaeaceae* together as “Njordarchaeales” within Heimdallarchaeia (Eme et al. 2023). However, the placement of this lineage within Heimdallarchaeia was only recovered with the inclusion of DNA topoisomerase subunit A (arCOG04143, Supplementary Figures 6, 7A), which is not an ideal marker to investigate relationship between Asgard/Njord/Panguiarchaea and Korarchaeota (see above). Finally, another recent study proposed the “Panguiarchaeales” as an order-level lineage within Korarcheia (Qu et al. 2023).

To resolve the taxonomic inconsistencies and normalise the rank, we applied a similar approach as integrated in a previous publication (Tamarit et al. 2024), which normalised the taxonomic boundaries in the GTDB reference tree to a phylogenetic tree inferred under a more complex phylogenetic model. Our analyses indicate a significant correlation (Pearson r: 0.96, p-value:0 ) between the relative evolutionary divergence (RED) values in the GTDB reference phylogenomic tree and the RED in our phylogenomic trees based on 256 mixture categories (Supplementary Figure 9A). Based on this correlation, we scaled the RED value used to delineate taxonomic ranks in the GTDB R220. Our analyses show that previously proposed "Njordarchaeales", "Njordarchaeota", and "Panguiarchaeales" likely form a putative class-level lineage, the Njordarchaeia (RED value: 0.2403). Further, we propose that this class contains the putative order-level lineage Njorarchaeales and two family-level lineages, i.e. *Panguiarchaeaceae* (previously "Panguiarchaeales") and *Njordarchaeaceae* (previously "Njordarchaeales"), based on RED values (Supplementary Figure 9B), and distinct metabolic features and inferred lifestyles (See below). However, better sampling of high-quality genomes of Asgard archaea, especially of the Wukongarchaeia, Njordarchaeia and Heimdallarchaeia will be needed to fully resolve the taxonomy of members of this clade.

#### 1.3 Addressing site-heterogeneity with the CAT-PMSF approach

Because site removal and recoding analyses result in a significant loss of potentially valuable information (Aouad et al. 2019; Sorokin et al. 2019; Fan et al. 2020; Martijn et al. 2022), we next analysed our full datasets using an approach that not only accounts for across-site substitutional heterogeneity but also adapts to the substitutional complexity of the dataset.

Specifically, we used the recently developed CAT-posterior mean site frequencies (PMSF) pipeline (Szánthó et al. 2023) to assess the support for the Asgard archaea and Korarchaeota topology. PMSF approaches require a guide tree, and the choice of a guide tree was previously shown to have an impact on the quality of phylogenetic inference (Wang et al. 2018; Wang et al. 2019). To account for this, we conducted CAT-PMSF analysis based on both LG and Poisson exchangeability matrices using different guide trees (with Asgard archaea topology and Korarchaeota topology, see Methods). Our CAT-PMSF phylogenetic inferences based on posterior site distributions extracted from different chains based on either LG or Poisson exchangeability matrix consistently yielded topologies supporting the placement of Njorarchaeia within the Asgard archaea (Supplementary Figures 12-13) regardless of the topology in guide trees. In line with this, approximately unbiased (AU) tests based on CAT-PMSF models from all chains confidently and consistently rejected the Korarchaeota topology ($p<0.05$; Supplementary Data 5). By contrast, the AU test based on the LG+C60+G+F model rejects none of the topologies (Supplementary Data 5). However, the simpler model LG+F+G rejected the Asgardarchaeta topology, likely due to the poor modelling of site heterogeneity, whose effects are the strongest a more constrained sites (Supplementary Data 5, Figure 1D). This provides robust analytical evidence that the Korarchaeota topology is a phylogenetic artefact due to the failure to model site-wise compositional heterogeneity accurately.

#### 1.4 Conflicting signals in analyses under the LG+C60+G+F model are likely due to inadequate modelling of site compositional heterogeneity

Profile mixture models such as C60 use a finite set of stationary distributions with different weights (Si Quang et al. 2008), whose performance is associated with the composition of effective amino acids of used sets of stationary distribution and their weights (Schrempf et al. 2020). To compare the performance of C60 in site-heterogeneity modelling in this dataset, we estimated empirical stationary distribution models (EDM) with different components using EDCluster from one chain based on the LG+CAT-PMSF model that gave the highest likelihood score. We inferred different trees based on different EDM models, LG+G+F and LG+C60+G+F models, using the streamlined set 2. Our results demonstrated that Njordarchaeia placement strongly depends on the number of components used (Supplementary Figures 10,11). In particular, the site homogenous model (LG+G+F, Supplementary Figure 10A) and EDM models with few components (4-8) favoured the Korarchaeota topology (Supplementary Figure 10BC). By contrast, EDM models with moderate components (16-32) preferred the sisterhood of Njordarchaeaia to the Wukong-Heimdallarchaeia clade (Supplementary Figure 10DE). Sifarchaeia branched, however, sister to the Jordarchaeia-Odinarchaeia clade. Interestingly, both LG+C60+G+F (Supplementary Figure 10J) and LG+EDM064+G+F models (Supplementary Figure 10F) supported the sisterhood of Njordarchaeia with the other Asgard archaea. Finally, the use of models with a higher number of components (128 and 256) as well as LG+CAT-PMSF recovered the sisterhood of Njordarchaeia and the Wukong-Heimdallarchaeia clade with strong support (>90/>90) (Supplementary Figure 10GHIK). These latter models also provided a better fit to the dataset based on Akaike and Bayesian Information Criteria/model adequacy test (Supplementary Figure 11A, Figure 1E). The LG+F+G showed the lowest model fit because it assumes a constant amino acid frequency at all sites. However, under different biogeochemical and selective constraints, empirical alignments often showed different compositions in different sites. This suggests that the effective amino acids per site in our alignment are lower than the prediction of LG+F+G ($K_{eff}$: 15.52, calculated from the empirical alignment). By contrast, the LG+EDM(4-64)+G+F and LG+C60+G+F accounted for across-site heterogeneity and showed improvement in model fits compared to LG+F+G (Supplementary Figure 11A). Additionally, more weights are put under the constrained distributions in the better fitting models (i.e., LG+EDM0128+G+F and LG+EDM0256+G+F), which is more similar to LG+CAT-PMSF (Supplementary Figure 11B). Similarly, the average number of substitutions inferred based on different models increased with their model fit (Supplementary Figure 11C). Taken together, this shows that LG+F+G underestimates the probability of convergent evolution. Models such as LG+C60+G+F, despite accounting for site-wise compositional heterogeneity, might still underestimate the probability of homoplasy between Njordarchaeia and Korarchaeota, comprising putative hyperthermophilic lineages.

In sum, our thorough analysis supports the Asgard archaea topology, highlighting the necessity to carefully assess phylogenetic inferences and the fit of evolutionary models to complex datasets.

### 2. Occurrence and phylogeny of Eukaryotic signature proteins (ESPs)

#### 2.1 Occurrence of ESPs

An important feature of the genomes of Asgard archaea is that they collectively encode more ESPs than other archaea (Spang et al. 2015a; Liu et al. 2021). To investigate the distribution of ESPs in *Panguiarchaeaceae* and *Njordarchaeaceae*, we queried the archaeal amino acid sequences against a recently developed Asgard Cluster of Orthologs database (Liu et al. 2021) and retrieved homologs with an updated accession list for previously identified ESPs (Eme et al. 2023). Phylogenies of actins, ribosomal L28e, TRAP beta, OST3/OST6 and Snf7 proteins were inferred to study the relationship between *Panguiarchaeaceae* and *Njordarchaeaceae* and other Asgard archaea. Our analyses based on the occurrence of asCOG accessions revealed that Njordarchaeales MAGs likely have a reduced repertoire of ESPs (Figure 3, Supplementary Data 7). Nevertheless, the clustering based on the nearest point algorithm using the occurrence of asCOGs indicated that the Njordarchaeia carry more ESPs than many other archaea and are very different from Korarchaeota (Figure 3, Supplementary Data 7). Notably, a non-asgard archaeal cluster was recovered in the dendrogram (Figure 3, denoted in star), likely reflecting a shared common ancestry.

#### 2.2 Phylogeny of ESPs

##### 2.2.1 Ribosomal protein L28e/Mak16 homologs

Eukaryotic Mak16 homologs (PF01778) are predicted to be involved in the maturation of 5.8S rRNA and the large subunit rRNA, whose distribution in archaea is restricted to members of the Hodarchaeales, “Njordarchaeales” and Wukongarchaeia (Eme et al. 2023). We detected MAK16 homologs in 7 representatives of the Njordarchaeia , which clustered sister to homologs of Wukongarchaeia and Hodarchaeales in agreement with the species tree (Supplementary Figure 15). In turn, this further supports the hypothesis of Njordarchaeia being a clade within the Asgard archaea.

##### 2.2.2 Actin homologs

Asgard archaea typically encode so-called lokiactins, i.e. conserved actin homologs, which might function in the formation of cytoskeleton (Spang et al. 2015a; Zaremba-Niedzwiedzka et al. 2017; Imachi et al. 2020; Rodrigues-Oliveira et al. 2023). All Njordarchaeia MAGs encode a single actin homolog (IPR004000), in contrast to other Asgard archaea such as Wukongarchaeia, Hodarchaeales and Lokiarchaeia, which, besides a conserved lokiactin, encode more distant actin-related proteins (Supplementary Data 9, Supplementary Figure 16,17). Phylogenetic analyses of actin family proteins revealed that Njordarchaeia homologs are distinct from the crenactin homologs found in Korarchaeota (Guy and Ettema 2011) and instead form a clade with some bathyarchaeial homologs sister to *bona fide* asgard archaea lokiactins (Supplementary Figure 16). Indeed, the removal of 20% of the most heterogenous sites recovered the monophyly of Njordarchaeia lokiactins with those of other Asgard archaea to the exclusion of the Bathyarchaeia (Supplementary Figure 17).

##### 2.2.3 N-glycosylation (OST3/OST6 and TRAP Beta):

Previous studies identified various asgard archaea proteins related to N-glycosylation, including various ESPs (Zaremba-Niedzwiedzka et al. 2017; Eme et al. 2023). Here, we focused on the phylogenetic analyses of two related proteins, the beta subunit of translocon-associated protein (TRAP) and OST3/OST6-like of the oligosaccharyltransferase (OST) complex. These complexes are thought to be involved in the biogenesis of N-glycosylated proteins (Pfeffer et al. 2017). We only identified two homologs of OST3/OST6-like proteins from Njordarchaeia representatives (e-value: 0.0001), which, in phylogenetic analyses, branched sister to Wukongarchaeia with medium support (62/80) and a Caldarchaeales MAG with maximal support (Supplementary Figure 19), respectively.

By contrast, homologs of TRAP-beta have a wider distribution among Njordarchaeia MAGs. Our analysis recovered the TRAP-beta homologs of Njordarchaeia sister to a copy from Sifarchaeotum subterraneus and clustering within the Heimdallarchaeia (Supplementary Figure 18) again supporting the hypothesis that Njordarchaeia are Asgard archaea.

##### 2.2.4 SNF7 proteins

Eukaryotic SNF7 domain proteins constitute a subunit of eukaryotic ESCRT-III complex (Tang et al. 2015). SNF7 homologs have previously been identified in both members of the Thermoproteota and Asgard archaea. In particular, most Asgard archaea encode two paralogs of SNF7 domain proteins which are related to eukaryotic VPS20/32/60 and VPS2/24/46 families, respectively (Spang et al. 2015b; Zaremba-Niedzwiedzka et al. 2017; Hatano et al. 2022). While cell division protein B homologs belonging to the SNF7 protein family are present in various Thermoproteota, they have not been identified in Korarchaeota. The herein investigated Njordarchaeia MAGs encode homologs of both VPS20/32/60 and VPS2/24/46 families, though homologs of the latter family are more common (Supplementary Figure 20). Phylogenetic analyses of these proteins resolved the CdvB protein as a divergent clade relative to the VPS20/32/60 and VPS2/24/46 clades (Supplementary Figure 20), in agreement with previous analyses (Spang et al. 2015c; Zaremba-Niedzwiedzka et al. 2017; Lu et al. 2020; Hatano et al. 2022). Our phylogenetic analysis of an untreated SNF7 family protein alignment revealed that the VPS20/32/60 homologs of Njordarchaeia formed a clade basal to the other VPS20/32/60 and VPS2/24/26 homologs (Supplementary Figure 20A). In contrast, upon removal of 20% of the most site-heterogeneous sites, homologs of the Njordarchaeia branched within the Heimdall-Sif-Wukongarchaeia cluster (Supplementary Figure 20B). Similarly, the phylogenetic analysis without eukaryotic sequences recovered the VPS20/32/60 homologs of Njordarchaeia within rather than basal to the VPS20/32/60 and VPS2/24/46 members (Supplementary Figure 20C). This suggests that compositional biases in Njordarchaeia homologs caused the artificial placement of VPS20/32/60 basal in the initial phylogeny. Taken together, SNF7 family proteins of the Njordarchaeia support their assignment to the

Asgard archaea. Our analyses suggest that *Panguiarchaeum* might have lost SNF7 family homologs (Supplementary Data 13), in agreement with their reduced metabolic potential and putative symbiotic lifestyle (see below).

### 3. Metabolism

#### 3.1 Informational processing and repair systems

##### 3.1.1 DNA replication and repair

We identified most genes encoding for proteins involved in DNA replication and cell division in Njordarchaeia MAGs (Supplementary Figure 24, Supplementary Data 8,9). In particular, Njordarchaeia MAGs encode two DNA polymerases: DNA polymerase B1 (PolB1, arCOG00328), which is widespread among different archaeal lineages (Supplementary Figure 24), and DNA polymerase II (PolC, arCOG04447 and arCOG04455). Additionally, Njordarchaeia MAGs encode at least two copies of ORC1-type DNA replication initiation protein 1 (Orc1/Cdc6, arCOG00467), which is common among archaea except in Methanococci and Methanpryi (Supplementary Figure 24, Supplementary Data 10) (Raymann et al. 2014). They also encode other replication-related proteins commonly found in archaea, such as a putative DNA helicase (Mcm2, arCOG00439), archaeal single-stranded DNA-binding replication protein A (RPA1, arCOG01510) and DNA-dependent ligase (Lig, arCOG01347). Finally, several different DNA topoisomerases were present in Njordarchaeia MAGs, including a reverse gyrase (TopG2, arCOG01526) and DNA topoisomerase VI subunit A and subunit B (Top6A, arCOG01165 and Top6B, arCOG04143). Although the distribution of reverse gyrase is sparse in archaea, the presence of reverse gyrase in Njordarchaeia is in line with their occurrence in hot environments, i.e., hydrothermal vents (Eme et al. 2023) or hot springs (Qu et al. 2023), suggesting Njordarchaeia members are potential hyperthermophiles. Furthermore, DNA topoisomerase IA (TopA, arCOG01527) and DNA gyrase B (GyrB, arCOG04371) are present in some *Njordarchaeaceae* MAGs (Supplementary Figure 24, Supplementary Data 8).

##### 3.1.2 Evolution of reverse gyrase

Our reconciliation analyses suggest that the reverse gyrase was present in the last Asgard archaea common ancestor (Supplementary Data 12) with high support (PP=0.96). We further assessed the amount of vertical evolution of this protein family along the species tree by using a verticality metric (Coleman et al. 2021). In line with the monophyly of Wukongarchaeia and *Njordarchaeaceae* recovered in the reverse gyrase phylogeny (Supplementary Figure 33A), our results suggest that this protein likely evolved vertically within the Sif-Wukong-Njord-Heimdallarchaeia lineages, combined with loss in some representatives (verticality of 577: 0.99, 571: 0.95, 502: 0.99, 563: 0.95) (Supplementary Figure 34). However, the reverse gyrase homologs of *Panguiarchaeum* formed a distinct clade (Supplementary Figure 33), suggesting a later horizontal acquisition of reverse gyrase in this clade (verticality of 481: 0.34, loss frequency: 1.01, transfer frequency: 0.69).

This, together with previous work, suggests that the evolutionary history of this protein family is complex (Brochier-Armanet and Forterre 2007; Catchpole and Forterre 2019; Eme et al. 2023) and is characterised by transfers and losses across archaeal lineages (Supplementary Figures 33,34). Similar to Eme et al. 2023 (Eme et al. 2023), we did not recover a monophyletic clade of all homologs of the Asgardarchaeaota. Based on this and the presence of reverse gyrase in only Njordarchaeia, Wukongarchaeia, Baldrarchaeia, Asgardarchaeia and Jordarchaeia, we cannot fully exclude the alternative scenario suggesting that reverse gyrase was acquired independently by distinct Asgard archaea rather than encoded by their common ancestor (Eme et al. 2023). More genomes will help to further assess the evolution of reverse gyrase.

##### 3.1.2 Transcription

Njordarchaeia MAGs encode most subunits of the RNA polymerase, including the core subunits that are shared and conserved among bacteria, archaea and eukaryotes, i.e., subunit A1 (RpoC/Rpo3: arCOG04257), subunit B (RpoB/Rpo2: arCOG01762), subunit D (RpoA/Rpo1: arCOG04241), subunit K (Rpo6/RpoZ: arCOG01268), subunit L (RpoL: arCOG04111), and the auxiliary subunits that are shared between archaea and eukaryotes, i.e., subunit H (RPB5, arCOG04258), subunit E (RPB7, arCOG00675), subunit N (RPB10, arCOG04244) and subunit P (RPO12, K03059). Moreover, subunit F (Rpo4, arCOG01016), subunit G (RPB8, arCOG04271) and Rpo13 (arCOG05938) appeared absent in Njordarchaeia MAGs as in many other archaeal lineages (Supplementary Figure 25). Subunit G and Rpo13 are sparsely distributed among thermoproteotal lineages such as Sulfolobales, Thermoproteales, Asgard archaea (e.g., Jordarchaeia) and Korarchaeota and may not be essential to a functional DNA-directed RNA polymerase (Koonin et al. 2007). Though subunit F appeared largely absent in Njordarchaeia MAGs while present in many other Archaea, it is unclear whether this subunit is essential. E.g. a RpoF- lacking Thermococcus kodakarensis mutant could grow though it did exhibit a temperature-sensitive phenotype (Hirata et al. 2008). Hodarchaeales and Njordarchaeia representatives were found to encode a fused version of RNA polymerase subunit A (Supplementary Figure 25) (Zaremba-Niedzwiedzka et al. 2017; Eme et al. 2023). The patchy distribution of fused RNA polymerase subunit A in archaeal lineages of Korarchaeota, Baldrarchaeia and Pacearchaeota is consistent with the previous hypothesis that gene fission and fusion of RNA polymerase subunit A occurred several times independently across different archaeal lineages (Zaremba-Niedzwiedzka et al. 2017). Transcription initiation and elongation factors common to archaea were also found in Njordarchaeia members, such as (TFIIB/SUA7, arCOG01981) and transcription elongation factor TFIIS (RPB9/TFS, arCOG00579, arCOG00580). *Panguiarchaeum* members appeared to have a reduced number of transcription factors compared to *Njordarchaeaceae* MAGs and other archaea (Supplementary Figure 25). For example, transcription elongation factor Spt5 (NusG/Spt5, arCOG01920) and transcription elongation factor (NusA, arCOG01760 and arCOG01761) appeared absent in *Panguiarchaeum* MAGs. In addition, transcription elongation factor Elf1 (arCOG04136) and transcription-termination-related protein Eta (arCOG00554) that are commonly found in other asgard archaea lineages seemed absent from Njordarchaeia

MAGs. Although comparative genomic analyses suggested that the gene loss of transcriptional regulators in some endosymbiotic bacteria such as *Buchnera* might occur (Chong et al. 2019), it remains to be determined if this represent true biological signal or is due to the incompleteness of MAGs, considering that many other DPANN archaeal lineages except Huberarchaeia encode a nearly complete set of transcriptional factors (Supplementary Figure 25, Supplementary Data 10).

##### 3.1.3 Translation

Most genes involved in translation, such as ribosomal proteins, are present in Njordarchaeia representatives (Supplementary Figures 26,27, Supplementary Data 8-10). To be more specific, many ribosomal proteins specific to Theromproteota, Korarchaeota and Asgard archaea lineages, such as L13E, L38E and S30, are also present in Njordarchaeia MAGs. They encode all archaeal tRNA synthetases as well (Supplementary Figure 27). In contrast to the absence of some transcriptional factors, most known translation initiation, elongation and termination factors are present in Njordarchaeia MAGs, such as translation initiation factor 1 (InfA, arCOG01179), translation initiation factor 6 (EIF6, arCOG04176), and peptide chain release factor (eRF1, arCOG01742). Diphthamide is a post-translationally modified histidine amino acid residue commonly present in the elongation factor 2 of archaea and eukaryotes. A recent study found that most Asgard archaea, Korarchaeota and Geoarchaeales lacked diphthamide biosynthesis gene (Narrowe et al. 2018), and many dph-genes-lacking organisms encode a second copy of the elongation factor 2 gene without key residues for histidine modification. Consistent with a previous analysis (Eme et al. 2023), genes for diphthamide biosynthesis appeared largely absent in Njordarchaeales MAGs, and most of the Njordarchaeia MAGs encode an additional copy of the elongation factor 2 gene (FusA, arCOG01559) (Supplementary Data 8,9).

#### 3.2 Metabolic features

##### 3.2.1 Central carbon metabolism

We investigated the central pathways for carbon metabolism and potential modes of energy conservation of representatives of the Njordarchaeia. We found that although Njordarchaeia members possessed similar metabolic profiles, members of the *Njordarchaeaceae* are metabolically more flexible than the *Panguiarchaeum* members. Njordarchaeia MAGs encode various numbers of carbohydrate degradation enzymes (CAZymes) (Supplementary Data 21). Interestingly, one genome (GCA_029856605.1) from the *Panguiarcheum* clade encoded several different glycoside hydrolases (GHs), including GH2 (beta-galactosidase/beta-glucuronidase), GH23 (transglycosylase), GH31 (alpha-glucosidase), GH33 (sialidases) and GH38 (alpha-mannosidase). Most *Panguiarchaeum* MAGs encode beta-galactosidase (arCOG07337.1, GH2), catalysing lactose into beta-galactose and glucose. Although we did not identify the gene encoding galactose mutarotase, which catalyses the interconversion between beta-galactose and alpha-galactose, a complete gene set encoding enzymes that catalyse the conversion between alpha-galactose and glucose was identified in Njordarchaeia MAGs. These include galactokinase (arCOG01029.1, GalK), UDP-glucose_4-epimerase (arCOG01376.1, arCOG01369.1, GalE), UDP-glucose pyrophosphorylase (arCOG00665.1, GalU), and phosphomannomutase/phosphoglucomutase (K15778, ManB), suggesting the potential to utilise galactose. We also investigated the presence of relevant sugar transporters in Njordarchaeia MAGs, which indicates that members of this group encode components of a putative ABC-type sugar transport system, including both a permease (UgpE, arCOG00159; UgpA, arCOG00157) and periplasmic component (UgpB, arCOG00151). However, the ATPase binding component protein appeared absent. Simple sugars could perhaps be taken up by passive diffusion.

The upper glycolytic pathway in Njordarchaeia MAGs is mostly complete, except for the key enzyme, the 6-phosphofructokinase (pfkA, arCOG03641 and arCOG03370), which appeared absent in all Njordarchaeia MAGs except GCA_029856705.1 (Supplementary Data 8). Notably, various nucleases such as exonuclease III (XthA, arCOG02207) and ATP-dependent RNA helicase (DeaD, arCOG00558), and genes encoding enzymes in nucleoside degradation pathway including, AMP phosphorylase (DeoA, arCOG02013), ribose 1,5-bisphosphate isomerase (arCOG01124) and ribulose 1,5-bisphosphate carboxylase (RbcL; RuBisCO, arCOG04443) were present in the *Njordarchaeaceae* MAGs but not in *Panguiarchaeum*. This suggests that *Njordarchaeaceae* members might be able to break down nucleic acids into nucleoside triphosphates, which can be fed into the lower glycolytic pathway in the form of 3-phosphoglycerate via the nucleoside degradation pathway. As most archaea, Njordarchaeia MAGs encoded 3-phosphoglycerate kinase (arCOG00496, Pgk) and pyruvate kinase (arCOG04120, PykF), which converts 1,3-bisphosphoglycerate to 3-bisphosphoglycerate and phosphoenolpyruvate to pyruvate, respectively and forms ATP. Additionally, in contrast with *Panguiarchaeum*, *Njordarchaeaceae* might be able to convert oxaloacetate to phosphoenolpyruvate using ATP-dependent phosphoenolpyruvate carboxykinase (PckA, arCOG06073). Njordarchaeia MAGs encoded a putative pyruvate ferredoxin oxidoreductase complex (PorA, arCOG01608; PorB, arCOG01601; PorD, arCOG01604; and PorG, arCOG01603) that may allow to catalyse the conversion of pyruvate to acetyl-CoA and reduce ferredoxin, which is common in anaerobic archaea (Bräsen et al. 2014). Acetyl-CoA could be further metabolised to acetate via the ADP-forming acetyl-CoA synthetase (AcdAB; arCOG01340 and arCOG01338) in Njordarchaeia members, producing ATP. The lack of pfkA in most Njordarchaeia representatives is puzzling since they harbour most enzymes of the glycolytic pathway and encode various GHs. Hence, it remains to be determined whether the absence of this gene is due to genome incompleteness or represents a true biological signal.

Furthermore, almost all Njordarchaeia MAGs encoded several putative enzymes of the tricarboxylic acid cycle (TCA) including furmase (FumA, arCOG04406; TtdA, arCOG04407), succinyl-CoA synthetase (SucC, arCOG01337; SucD, arCOG01339) and a 2-oxoglutarate/2-oxoacid ferredoxin oxidoreductase (KorA, K00174; KorB: K00175), which allows the conversion between malate and fumarate, and between succinate and oxoglutarate. However, the enzymes of the other steps in the TCA cycle were limited in the *Panguiarchaeum* MAGs. For example, citrate synthase (GltA, arCOG04237), malate dehydrogenase (Mdh, arCOG00246), and aconitase (AcnA, arCOG01697) were only identified in the *Njordarchaeaceae* MAGs. Given the incomplete nature of the TCA cycle and the absence of pfkA in most Njordarchaeia MAGs, the identified proteins might be involved in the fermentative processes.

##### 3.2.2 Peptides/amino acids degradation

Additionally, Njordarchaeia MAGs encode various putative peptidases, aminotransferases and proteases (Supplementary Data 20), among which we detected a signal peptide in a peptidase (MER0283003; C1A.UPA) based on SignalP (Almagro Armenteros et al. 2019). This suggests that they may be able to break down peptides, though most of the peptidases are likely located intracellularly. Amino acids or peptides could be transported through membranes by amino acid transporters (PotE, arCOG00009) or a peptide transport system such as an ABC-type antimicrobial peptide transport system (SalX and SalY, arCOG00922 and arCOG02312). Peptides can be degraded into amino acids by different peptidases and proteases and further converted to 2-oxoacids by aminotransferases such as aspartate/tyrosine/aromatic aminotransferase (arCOG01131). 2-oxoacids such as pyruvate could subsequently be metabolised by enzyme complexes such as pyruvate ferredoxin oxidoreductase complex or related 2-oxoacid:ferredoxin oxidoreductase / 2-oxoglutarate/2-oxoacid ferredoxin oxidoreductase (KorA, K00174; KorB: K00175) into acyl-CoA/acetyl-CoA and metabolised into small organic acids such as succinate/acetate by CoA synthetases (AcdAB; arCOG01340 and arCOG01338, SucCD, arCOG01337 and arCOG01339). Additionally, all Njordarchaeia MAGs encode a putative aldehyde:ferredoxin oxidoreductase (Aor, arCOG00706), which could convert an aldehyde to its corresponding acid while reducing ferredoxin. It was reported that the pyruvate ferredoxin oxidoreductase complex of Pyrococcus furious catalyses the formation of acetaldehyde from pyruvate in a CoA-dependent reaction (Ma et al. 1997). It seems possible that the 2-oxoacids generated from amino acids/peptide degradation could be converted to acetaldehyde which could be subsequently converted into acetate via the aldehyde:ferredoxin oxidoreductase.

##### 3.2.3 Redox balance and energy conservation

Njordarchaeia members encode various enzymes that may participate in the reduction and oxidation of NAD(P)H and NAD(P)+, as well as ferredoxins. Most of the representative MAGs investigated encode a putative glyceraldehyde-3-phosphate dehydrogenase (GapA, arCOG00493), pyruvate ferredoxin oxidoreductase, 2-oxoglutarate/2-oxoacid ferredoxin oxidoreductase (KorA, K00174; KorB: K00175), aldehyde ferredoxin oxidoreductase (Aor, arCOG00706) and in the case of some *Njordarchaeaceae* MAGs, malate dehydrogenase (Mdh: arCOG00246). All these proteins could couple the generation of reduced ferredoxin or NAD(P)H to the carbon metabolism. Additionally, Njordarchaeia MAGs harbour different hydrogenases, including group 3c [NiFe]-hydrogenases, group 3b [NiFe]-hydrogenases, and group 4g [NiFe]-hydrogenase (Supplementary Figures 28,29). These hydrogenases were reported to be able to couple the oxidation of ferredoxin/NAD(P)H to hydrogen evolution. In particular, group 4g [NiFe]-hydrogenases contained a Nuo-L-like subunit which is believed to be involved in proton gradient generation (Supplementary Figure 28B). The proton gradient could be harvested by a putative V-type ATP synthase (AtpABDEGIK; arCOG00868, arCOG00865, arCOG04101, arCOG00869, arCOG04102, arCOG04138 and arCOG02455). Moreover, some MAGs also encode putative $Na^+/H^+$-translocating membrane pyrophosphatases (HppA, arCOG04949), which may allow the translocation of sodium/hydrogen across the membrane.

Taken together, our analyses of the central carbon metabolism, amino acid/peptide degradation and energy conversation of the herein investigated Njordarchaeia MAGs indicated that they have the potential to ferment amino acids and perhaps also hydrocarbons (see main text) and can conserve energy via both substrate-level phosphorylation and the ATP synthase. *Njordarchaeaceae* members appeared more metabolically flexible than the *Panguiarchaeum* because they encoded more genes for carbohydrate metabolism.

#### 3.2.4 Purine and pyrimidine biosynthesis

While Njordarchaeia members lack both the oxidative and non-oxidative pentose phosphate pathway, most of them encode enzymes of a reductive ribulose-monophosphate pathway (RuMP), including a bifunction enzyme 3-hexulose-6-phosphate synthase/6-phospho-3-hexuloisomerase (phi/hps, K13831), and ribose-5-phosphate isomerase (RpiA), which allows the synthesis of ribose-5-phosphate from fructose-6-phosphate. The bifunctional enzyme phi/hps is common in archaea and was found in many different asgard archaea members (Bräsen et al. 2014; Padalko et al. 2024). Phosphoribosylpyrophosphate (PRPP), an important precursor for purine and pyrimidine biosynthesis, could be synthesized using phosphoribosylpyrophosphate synthetase (PrsA, arCOG00067). In contrast to *Njordarchaeaceae* MAGs, *Panguiarchaeum* MAGs encode only the adenylosuccinate lyase (PurB, arCOG01747) in the steps from PRPP to inosine monophosphate (IMP). Interestingly, most of the enzymes involved in synthesising deoxyadenosine triphosphate (dATP) and deoxyguanine triphosphate (dGTP) from IMP are present in Njordarchaeia MAGs. It is worthwhile to note that not all steps of the archaeal purine biosynthesis have been fully elucidated. For example, while bacteria use N5-carboxyaminoimidazole ribonucletoide (CAIR) synthase (PurK, arCOG01597) to catalyse the conversion of 5-AIR to N5-CAIR, this gene is absent in several hyperthermophilic organisms that live in an environment with high bicarbonate concentration (Brown et al. 2011) like *Ignicoccus hospitalis* (the host of *Nanoarchaeum equitans)*. Hence, it remains unclear if this step can happen non-enzymatically or whether it is performed by an unknown enzyme. Guanylate kinase (Gmk, K00942) is absent from all Njordarchaeales MAGs, similar to what was reported for other archaeal genomes (Dombrowski et al. 2020). Similarly, both the phosphoribosylaminoimidazole-succinocarboxamide formyltransferase/ IMP cyclohydrolase (PurH, arCOG02824) and IMP cyclohydrolase (PurO, arCOG04727) that catalyse the formation of IMP appear to be absent in Njordarchaeia MAGs. The de novo biosynthesis pathway in *Panguiarchaeum* MAGs is likely incomplete, given the absence of most genes from PRPP to IMP. In contrast to the *Njordarchaeaceae* MAGs, *Panguiarchaeaum* MAGs encode permease and ATPase components (NupOPQ: COG3845, COG4603 and COG1079) of an ABC-type guanosine uptake system, which might allow the transport of purine nucleosides into their cells.

Our analysis of the pyrimidine biosynthesis pathway revealed that most Njordarchaeia MAGs encode all the proteins necessary for the formation of deoxycytidine triphosphate (dCTP) and deoxythymidine triphosphate (dTTP) from carbamoyl phosphate. These proteins include aspartate carbamoyltransferase (PyrB, arCOG00911), dihydroorotase (PyrC, arCOG00689), dihydroorotate dehydrogenase (PyrD, arCOG00603), orotate phosphoribosyltransferase (PyrE, arCOG00029), and orotidine-5-phosphate decarboxylase (PyrF, arCOG00081), which catalyse the formation of uridine monophosphate (UMP) from carbamoyl phosphate. Moreover, almost all Njordarchaeia MAGs encode genes that further convert UMP to dCTP (PyrH, arCOG00858: uridylate kinase, Ndk, arCOG04313: nucleoside diphosphate kinase, PyrG, arCOG00063: CTP synthase and NrdD, arCOG04889: ribonucleoside-triphosphate reductase) and dCTP to dTTP (Dcd, arCOG04048: deoxycytidine deaminase, Tmk, arCOG01891: thymidylate kinase, ThyX, arCOG01883: flavin-dependent thymidylate synthase). NrdD, which catalyses the conversion of ribonucleotides into deoxyribonucleotides, is important for the growth of microorganisms under anaerobic conditions (Garriga et al. 1996; Crespo et al. 2017), and its presence coincided with the absence of cytochrome C oxidase in Njordarchaeia MAGs, suggesting an anaerobic lifestyle. However, the distribution of carbamoyl phosphate synthase (CarA: arCOG00064, CarB: arCOG01594) in Njordarchaeia MAGs, which converts glutamine to carbamoyl phosphate and initialises the first step of pyrimidine biosynthesis, appears to be restricted to GCA_029856705.1. Considering the presence of all other proteins in this pathway, it seems possible that the lack of carbamoyl phosphate synthase is due to genome incompleteness. The presence of a largely complete pyrimidine biosynthesis pathway in the putative symbiotic members of *Panguiarchaeum* differs from most DPANN cluster 2 members, such as members from Nanohaloarchaeota, Nanoarchaeales and Woesearchaeales and Pacearchaeales, which harbours an incomplete pyrimidine biosynthesis pathway (Supplementary Figure 23).

##### 3.2.5 Purine and pyrimidine salvage

Next, our analysis of genes encoding enzymes involved in the purine and pyrimidine salvage pathway revealed that most Njordarchaeia members encode adenine deaminase (AdeC, arCOG00693) and hypoxanthine phosphoribosyltransferase (Hpt, arCOG00040), which could catalyse the formation of IMP from adenine and the conversion from guanine to dGTP. The purine salvage pathway and potential transport system may compensate for the reduced purine biosynthesis potential. While *Panguiarchaeum* MAGs encode genes to convert cytidine to uracil (Cdd, arCOG04173 and Udp, arCOG01324), the uracil phosphoribosyltransferase (Upp, arCOG04128) which converts uracil to UMP seemed absent in *Panguiarchaeum* MAGs.

Altogether, our analyses of purine and pyrimidine biosynthesis and salvage indicated that members of *Njordarchaeaceae* possess a relatively complete repertoire for purine and pyrimidine synthesis. *Panguiarchaeum* members showed a reduced biosynthesis potential for purine and may depend on external sources.

##### 3.2.6 Amino acid biosynthesis

As mentioned above, Njordarchaeia MAGs encode various peptidases, most of which do not seem to have signal peptide components, suggesting that they may be involved in amino acid interconversions generating respective organic acids. For instance, the putative aspartate aminotransferase (AspB, arCOG01130) encoded by Njordarchaeia MAGs might convert aspartate and 2-oxoglutarate to glutamate and oxaloacetate and the glycine/serine hydroxymethyltransferase present in most Njordarchaeia MAGs may be involved in the interconversion of glycine and serine. Almost all Njordarchaeia MAGs (12 out 13) encode a putative serine-pyruvate aminotransferase (PucG, arCOG00082), which might transaminate serine and pyruvate to 3-hydroxypyruvate and alanine. Interestingly, although most Njordarchaeia MAGs encode 3-phosphoglycerate dehydrogenase (SerA, arCOG01754), homologs of phosphoserine phosphatase (SerB, arCOG01158) were not identified in *Panguiarchaeaum* clade but *Njordarchaeaceae*. While aminotransferase SerC (arCOG00083) appeared absent in Njordarchaeia MAGs, the function of this enzyme might be complemented by PucG which carries an aminotransferase class V domain (IPR000192), similar to *Methanocaldococcus* jannaschii that could use a broad-specificity class V aspartate aminotransferase to convert the phosphohydroxypyruvate product to phosphoserine (Helgadóttir et al. 2007).

Even though Njordarchaeia MAGs lack many genes involved in amino acid metabolism, they encode various systems to import amino acids/peptides. For example, an amino acid transporter PotE (arCOG00009), a putative ABC-type antimicrobial peptide transport system (SalXY), and a putative ABC-type dipeptide/oligopeptide/nickel transport system (DppBCD, arCOG00751, arCOG00749 and arCOG00181) was found present in most of the MAGs. Taken together, this suggests that Njordarchaeia might be able to acquire amino acids from the environment.

##### 3.2.7 Lipid biosynthesis

Our analysis of the lipid biosynthesis gene of the herein investigated Njordarchaeia MAGs revealed that the key genes for proteins involved in the biosynthesis of isoprenoid, i.e., the modified mevalonate pathway (MVA) (Hayakawa et al. 2018), are largely absent in *Panguiarchaeum* MAGs while present in the *Njordarchaeaceae* MAGs. These proteins include acetyl-CoA acetyltransferase (ACAT, arCOG01278), 3-hydroxy-3-methylglutaryl CoA synthase (HmgB, arCOG01767), hydroxymethylglutaryl-CoA reductase (HmgA, arCOG04260), mevalonate kinase (Mvk, arCOG01028), a putative aconitase X (AcnX, arCOG04279), UbiD-type decarboxylase (UbiD, arCOG01671), 3-polyprenyl-4-hydroxybenzoate decarboxylase (UbiX, arCOG01703), isopentenyl phosphate kinase (Ipk, arCOG00860), and isopentenyl diphosphate isomerase (Idi, arCOG00613). While one *Panguiarchaeum* MAG (GCA_029856605.1) encodes Mvk, which phosphorylates mevalonate and contains the archaeal mevalonta kinase domain (IPR022937), other enzymes in the MVA pathway appeared absent. Additionally, key genes for the eukaryotic-type MVA pathway (Pmk, K00938 and MvaD, K01597), the haloarchaea-type modified pathway (Pmd, K17942) and the thermoplasma-type modified pathway (M3K, K18689 and Bmd, K22813) seem to be absent in all Njordarchaeia MAGs (Supplementary Data 9). This suggests that Njordarchaeia uses the modified mevalonate pathway, which seems conserved among most archaea (Hayakawa et al. 2018). However, as pointed out earlier (Qu et al. 2023), the absence of a complete lipid biosynthesis pathway in *Panguiarchaeum* is surprising. Indeed, so far, incomplete lipid biosynthesis pathways have only been described in DPANN lineages. Specifically, many members of the proposed symbiotic DPANN archaeal lack a number of lipid biosynthesis genes (Castelle and Banfield 2018; Dombrowski et al. 2019; Dombrowski et al. 2020), and cultured representatives such as *N. equitans* and *Ca.* Nanohaloarchaeum antarcticus has been shown to acquire lipids from their respective hosts (Jahn et al. 2008; Ding et al. 2024). However, the absence of isoprenoid pathway is mostly found in the DPANN cluster 2 lineages (Supplementary Figure 23), and some cultivated Micrarchaea (Cluster 1 DPANN), such as *Microcaldus variisymbioticus* were reported to have complete MVA pathway (Sakai et al. 2022). To verify the absence of archaeal lipid biosynthesis proteins in Panguiarchaeaum, we conducted a six-frame translation query of *Panguiarchaeum* MAGs against MVA genes in other Njordarchaeia MAGs using blastX (settings: 1e-5). However, no significant homologs were found for *Panguiarchaeum* MAGs, except for the detection of a putative galactokinase (GalK, arCOG01029) using mevalonate kinase as query, consistent with a previous analysis (Qu et al. 2023).

Next, our analyses of genes involved in the biosynthesis of ether-linked archaeal lipids from dimethylallyl pyrophosphate (DMAPP) revealed a similar presence and absence pattern as the MVA pathway. Key genes for proteins synthesising 2,3-bis-Geranylgeranyl glycerol-1 phosphate appeared absent in *Panguiarchaeum* MAGs. These genes include geranylgeranyl pyrophosphate synthase (GGPS, arCOG01726), glycerol-1-phosphate dehydrogenase (GldA, arCOG00982), phosphoglycerol geranylgeranyltransferase (GGGPS, arCOG01085), and digeranylgeranylglyceryl phosphate synthases (DGGGP, arCOG00476). However, similar to *Njordarchaeaceae* members, *Panguiarchaeum* MAGs encode genes for protein involved in the activation and modification of polar head group, including CDP-archaeol synthase (CarS, arCOG04106), phosphatidylglycerophosphate synthase (PgsA, arCOG00670) and phosphatidylserine synthase (PssA, arCOG00673). The Njordarchaeia MAGs also encode putative digeranylgeranylglycerophospholipid reductases (GGR,arCOG00570), which could catalyse the hydrogenation of geranlygeranyl chains of unsaturated archaetidic acid, a reaction that remains unclear at which step it takes places in the pathway, allowing it to adjust membrane fluidity (Jain et al. 2014). Phylogenetic analyses of Njordarchaeia PssA homologs showed they formed a paraphyletic clade with Heimdallarchaeia (Supplementary Figure 31). Additionally, the analysis of genes involved in beta-oxidation, which could operate reversely to generate acyl-CoA, showed a similar pattern as the archaeal lipid biosynthesis genes in that *Panguiarchaeum* MAGs lack key genes for beta-oxidation (Figure 2). Missing genes include acetyl-CoA acetyltransferase (ACAT, arCOG01278), enoyl-CoA hydratase/ 3-hydroxyacyl-CoA dehydrogenase (ECH/HADH, K15016) and Acyl-CoA dehydrogenase (ACDH, K18244). All Njordarchaeia MAGs lack genes (COG2030, HCD, COG1064, ECR and COG1028, ARC) for a recently described fatty acid biosynthesis pathway in archaea (Schmerling et al. 2024). Although *Panguiarchaeum* MAGs might be able to generate glycerol-3-phosphate（G3P) via a G3P dehydrogenase (GpsA, K00057), genes encoding for PlsB/PlsC/PlsX/PlsY appeared absent in *Panguiarchaeum* MAGs. These genes, however, were reported to be present in some heimdallarchaeial members such as Hodarchaeales and Gerdarchaeales (Coleman et al. 2019).

Altogether, our analyses indicate that *Panguiarchaeum* lacks genes for de novo biosynthesis of isoprenoids and the geranylgeranyl chains, considering that most of the genes known to the archaeal lipid biosynthesis appeared absent. The *Panguiarchaeum* members might need to acquire lipids from their environment or rely on a potential partner. In the meantime, it is worthwhile to note that more MAGs/genomes, ideally circular, of *Panguiarchaeum* and culture representatives are needed to confidently determine the absence of glycerophospholipid pathway and lifestyle of members of this clade.

##### 3.2.8 Vitamin and cofactor biosynthesis

Njordarchaeia MAGs encode diverse genes that encode cofactor-dependent enzymes. For example, the pyruvate ferredoxin oxidoreductase encoded by Njordarchaeia MAGs contains thiamine-domain (IPR029601), the glyceraldehyde-3-phosphate dehydrogenase requires NAD(P) to function and the group 3c [NiFe]-dehydrogenase needs flavin adenine dinucleotide (FAD). In line with this, Njordarchaeia MAGs encode a complete set of genes involved in *de novo* thiamine biosynthesis, including archaeal ribulose 1,5-bisphosphate synthetase (Thi4, arCOG00574), 5-hydroxybenzimidazole synthase (ThiC, arCOG02741), hydroxymethylpyrimidine/phosphomethylpyrimidine kinase (ThiDN, arCOG00020) and thiamine monophosphate kinase (ThiL, arCOG00638) (Supplementary Data 8,9). In addition, although the riboflavin biosynthesis pathway appeared incomplete in Njordarchaeia, they encode an Ecf-type riboflavin transporter (arCOG05752) and riboflavin transporter (FmnP, arCOG03794), which might allow the uptake of riboflavin. Riboflavin could subsequently be converted to FAD via a CTP-dependent riboflavin kinase (arCOG01904) and archaeal riboflavin synthase (RibC2, arCOG01322). The biosynthesis of nicotinamide adenine dinucleotide (NAD) might also be feasible considering the presence of aspartate dehydrogenase (AspD, arCOG00254), quinolinate synthase (NadA, arCOG04459), nicotinate-nucleotide pyrophosphorylase (NadC, arCOG01482), nicotinamide mononucleotide adenylyltransferase (NadR, arCOG00972), and $NH_3$-dependent $NAD^+$ synthetase (NadE, arCOG00069). In turn, this may suggest that Njordarchaeia, including *Panguiarchaeum*, synthesize various cofactors.

### 4 Positive controls in the network inference

To assess the reliability of our network approach aiming at identifying ecologically meaningful associations among taxa, we subset the metagenomic community profiles derived from NCBI (248,559 metagenomes) annotated with GTDB R214 taxonomy (Parks et al, 2022) to the genera *Nanoarchaeum* and *Huberarchaeum*, respectively. Using the methodological pipeline described in the Methods, we checked to what extent we would identify known symbiont-host taxa: i.e. associations between *Nanoarchaeum equitans* and *Ignicoccus hospotalis* (Huber et al. 2002) and *Ca*. Huberarchaeum crystalense and *Ca.* Altiarchaeum hamiconexum (Schwank et al. 2019). Furthermore, we estimated the minimum number of samples needed for reproducible, reliable inference using random subsets of samples.

Specifically, we first obtained the two datasets from community profiles derived from 248,559 NCBI metagenomes (Woodcroft et al. 2024) from the ‘Sandpiper’ website (https://sandpiper.qut.edu.au), which were annotated with the GTDB R214 taxonomy (Parks et al., 2022). We selected all samples that contain taxa annotated as (i) Nanoarchaeum (30 samples) and (ii) Huberarchaeum (67 samples), respectively. We transformed the coverage profile of the taxa in these samples to counts by multiplying with 100 and formatted the data into a table listing files in rows and all taxa in columns (analogue to an OTU- or ASV-table).

To reduce the scarcity of the dataset, we filtered the dataset before network inference, depending on data distribution. For the Nanoarchaeum dataset, we filtered for taxa present in 10, 20, and 25 samples based on shoulder theory (reference). For the Huberarchaeum dataset, we filtered for taxa present in 10%, 20%, 30%, 40%, 50%, 60%, 70%, 80%, and 90% of the samples. Networks were inferred using SpiecEasi v1.1.3 (Kurtz Z et al, 2021). Across all networks, we checked for associations of Nanoarchaeum-Ignicoccus and Huberarchaeum-Altiarchaeum, respectively.
For estimating the minimum number of samples needed for reproducible, reliable network inference, we used the Huberarchaeum dataset, as, in contrast to the Nanoarchaeum dataset, it had enough samples for subsampling. To estimate the minimum number of samples required for robust, true positive prediction of Huberarchaeum-Altiarchaeum associations, we drew random samples and applied two different filter criteria. Numbers of 10, 20, 30, 40, 50, and 60 samples were drawn; the random data sets were filtered for taxa detected in at least 50% and 80% of all random samples, respectively. For each condition, ten networks were inferred. Statistics were run across all ten networks within one condition.
All analyses were performed in R v4.4.1 (Team, 2024).

In the dataset, there were two taxa annotated as Nanoarchaeum genus (taxa_658, taxa_3077); and six taxa annotated as Ignicoccus genus (taxa_3007, taxa_678, taxa_3062, taxa_3063, taxa_3922, taxa_4367). Some of the taxa were present in only very few samples (Supplementary Figure 35). In the inferred network, no association between any of these taxa was predicted. Furthermore, using correlation analysis, only a few of the taxa of the known symbiont-host systems were correlated (Supplementary Figure 36). The correlation may be too weak to be detected in the network inference. Interestingly, some of the significantly correlated taxa were (almost) perfectly negatively correlated. In other words, inside the dataset extracted, there was mutual exclusion rather than symbiosis (Supplementary Figure 36 C-E ).

In the dataset, on genus level, there were two taxa annotated as *Huberarchaeum* (taxa_128, taxa_1707) and four annotated as *Altiarchaeum* (taxa_127, taxa_208, taxa_559, taxa_836). Across all but one of the networks inferred after applying different filter conditions, we could detect one consistent association between a Huberarchaeum (taxa_1707) and an Altiarchaeum taxon (taxa_208). Specifically, only in the 90% network was the association not detected. The counts of both taxa were found to be correlated (Supplementary Figure 36F). While taxa_1707 and taxa_208 occurred in the dataset 60 and 61 times, respectively, the other taxa were detected in a maximum of 7 samples.

To check if occupancy, i.e. sample size is a limiting factor, we subset the dataset used for network inference (10, 20, 30, 40, 50, and 60 samples, respectively) and determined the minimum sample size allowing us to robustly detect the known associations found in the original network (taxa_1707-taxa_208). Before inferring the network of the randomly drawn samples, we filtered the random dataset for 50% and 80% occupancy, respectively. For the 80% occupancy networks, no associations were repeatedly found, independent of the number of samples used for network inference. For the 50% occupancy networks, the detection frequency of the expected association was dependent on the number of samples used for network inference (Supplementary Table 1). In all networks inferred from >30 samples, the expected association between taxa_1707 and taxa_208 was robustly detected. Below a number of 30 samples, the association was detected only in 50% of the samples.

This indicates a minimum number of > 30 samples, 40 samples, which is necessary for reliable inference of association.

Thus, our network inferences were able to detect one of the two positive controls, i.e the association between *Huberarchaeum* and *Altiarchaeum*. From the estimation of the minimum number of samples needed for correct network detection inferred from the *Huberarchaeum–Altiarchaeum* dataset, the minimum sample size appeared to be >30. Therefore, it is possible that the association between Nanoarchaeum equitans and Ignicoccus hospitalis was below the detection limit. It is known that Nanoarchaea other than N. equitans interact with other hosts, including other members of the Thermoproteota, which might further complicate the inference of an association. Finally, it is notable that in contrast to the counts of the *Huberarchaeum* and *Altiarchaeum* taxa in the dataset, the counts of the Nanoarchaeum and Ignicoccus taxa were not correlated.

**Supplementary Table 1. Dependence of detection frequency on sample size.** From the original dataset subset of different sizes (10-60 samples) were drawn. For each sampling size, ten subsets consisting of randomly drawn samples were drawn. Random datasets were filtered discarding taxa occurring in < 50% of the samples. Networks were inferred from filtered random subsets. The number of times the wet lab confirmed, expected association was detected in the networks was counted.

| Sample size for network inference | Detection frequency |
|---|---|
| 10 | 5/10 |
| 20 | 6/10 |
| 30 | 5/10 |
| 40 | 10/10 |
| 50 | 10/10 |
| 60 | 10/10 |

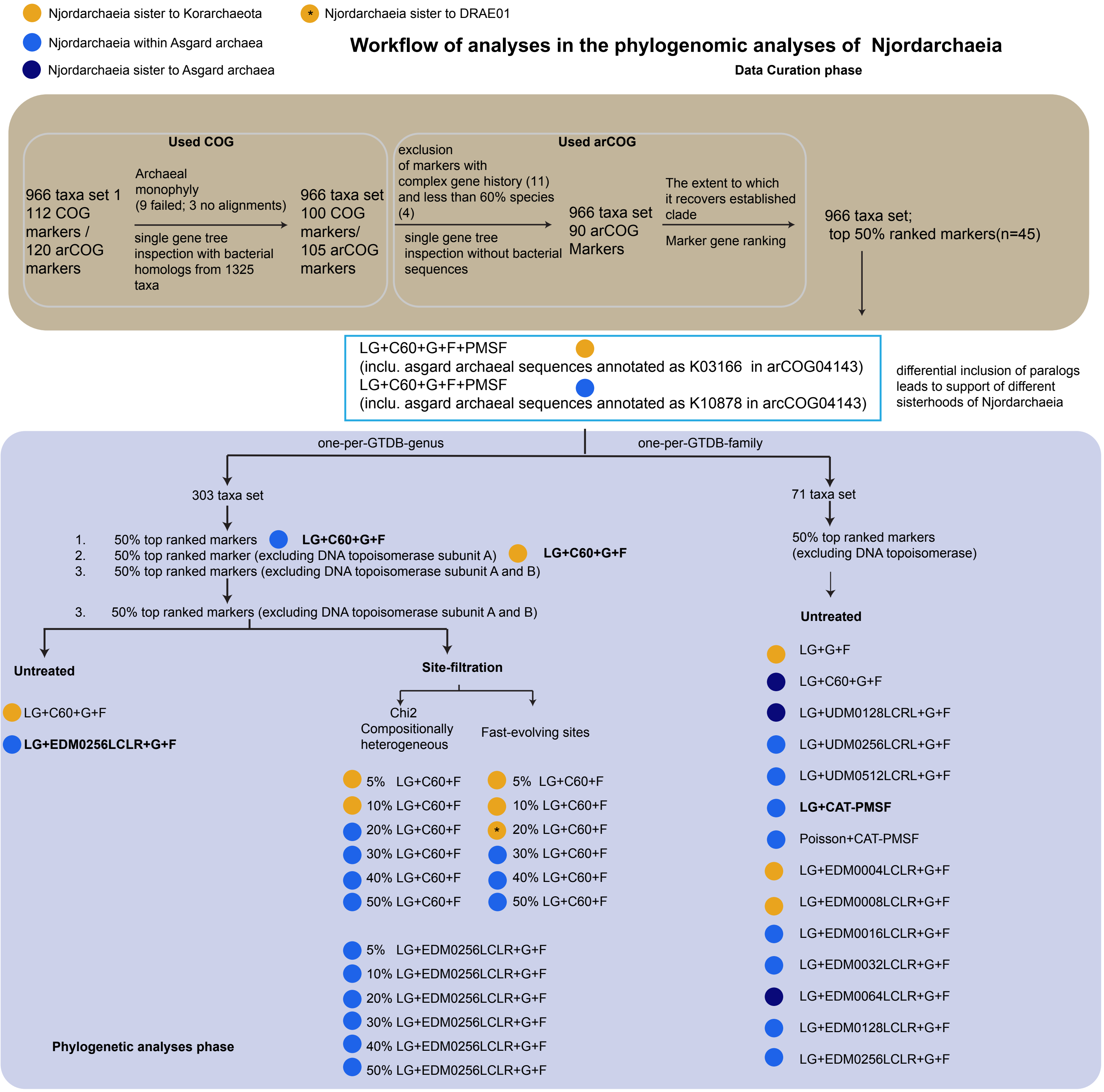


**Supplementary Figure 1.** Summary of phylogenetic workflow in this study. The analyses consist of two phases: the data curation phase, which consists of single gene tree inspection and evaluation of markers, and the phylogenetic analyses phase, which a variety of phylogenetic inferences was performed using taxa set1 (966 taxa), streamlined set 2 (303 taxa) and streamlined set 3 (71 taxa). DNA topoisomerase subunits were removed from the concatenation analyses due to the occurrence of paralogs in certain lineages with inconsistent placements (see, for example, **Supplementary Figures 2-8**). The placement of Njordarchaeia, including the Panguiarchaeum clade, recovered in the respective maximum-likelihood analyses, is shown in circles with the colour code in the figure inlet. Best-fit-ting models of untreated datasets are shown in bold font.

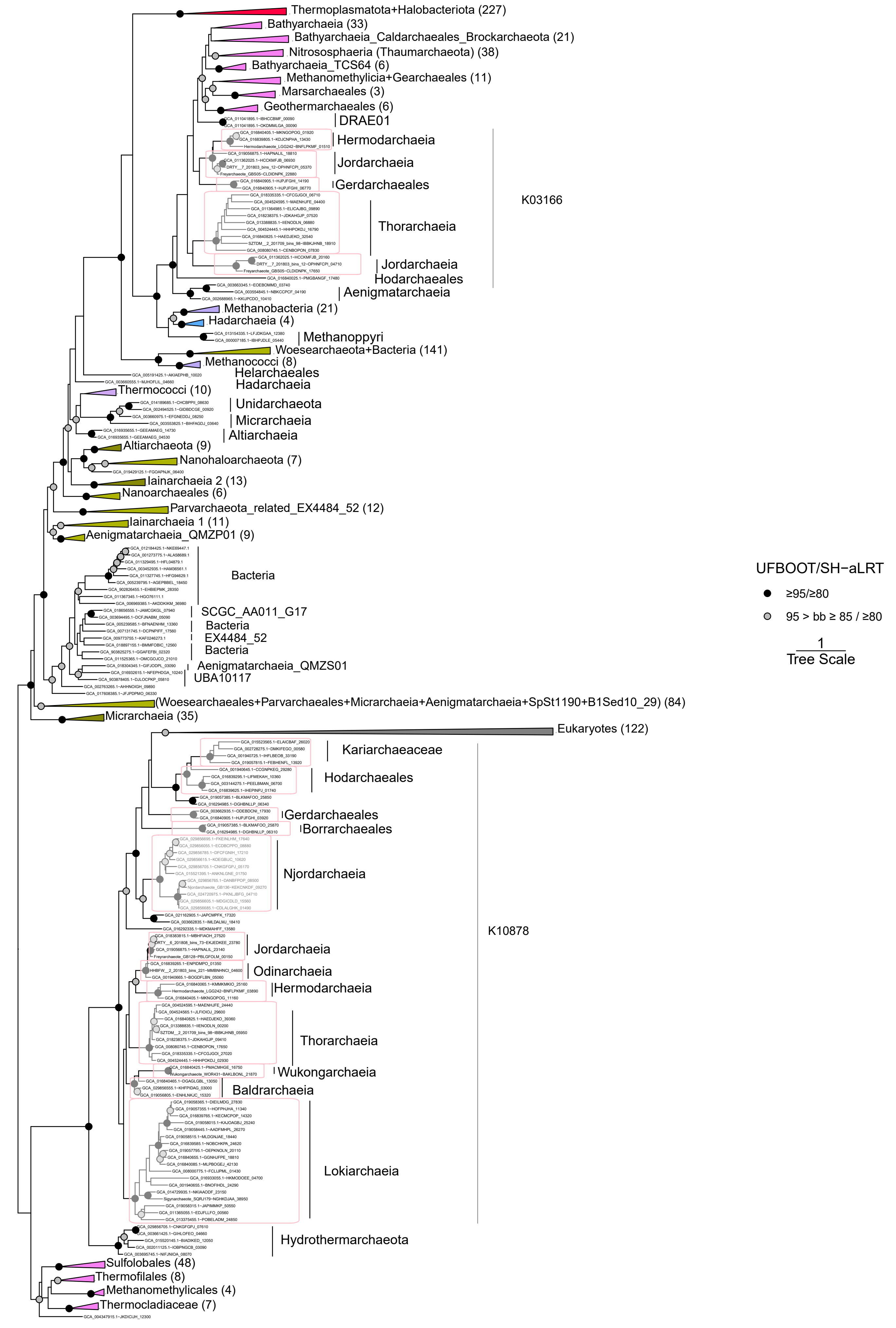


**Supplementary Figure 2. Phylogenetic analysis of DNA topoisomerase subunit A. (Top 6A).** The homologs were retrieved using COG1697 as described in the methods. Long branching sequences from Korarchaeota, Thorarchaeia and Odinarchaeia were removed. Note that Asgard archaea sequences are placed in two different clades, i.e., K03166, DNA topoisomerase VI subunit A, and K10878, meiotic recombination protein SPO11. The alignment includes 1041 sequenc-es and 204 amino acid sites. The phylogeny is manually rooted. The analysis used the LG+C60+G+F+PMSF model with the guide tree inferred using the LG+F+G model. Scale bar: Average substitution per site.

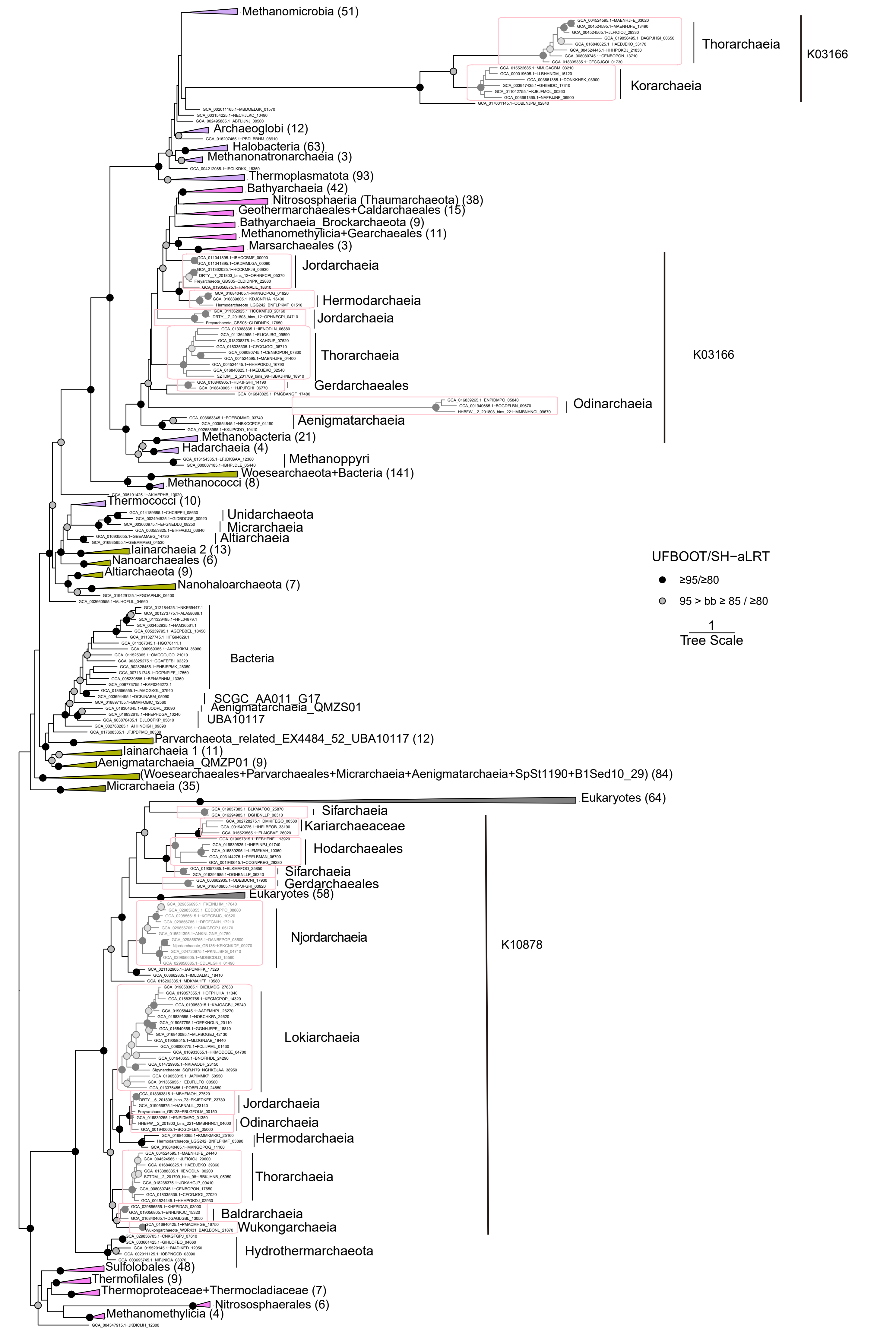


**Supplementary Figure 3. Phylogenetic analysis of DNA topoisomerase subunit A. (Top 6A).** The homologs were retrieved using COG1697 as described in the methods. Long branching sequences from Korarchaeota, Thorarchaeia and Odinarchaeia are included. Note that Asgard archaea sequences are placed in two different clades, i.e., K03166, DNA topoisomerase VI subunit A, and K10878, meiotic recombination protein SPO11; further, Korarchaeotal sequences branched close to the K03166 clade. The alignment includes 1065 sequences and 295 amino acid sites. The phylogeny is manually rooted. The analysis used the LG+C60+G+F+PMSF model with a guide tree inferred using the LG+F+G model. Scale bar: Average substitution per site.

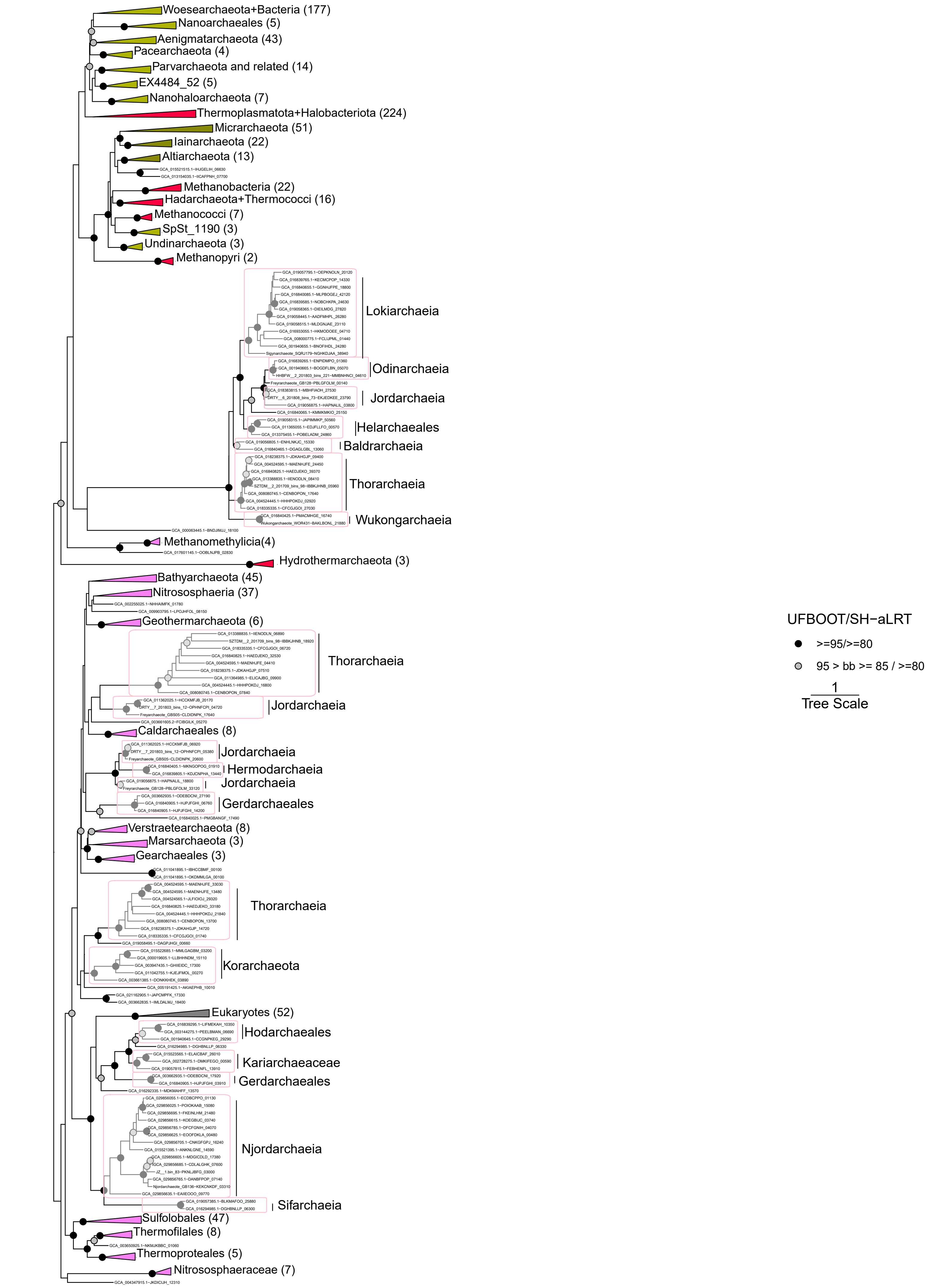


**Supplementary Figure 4. Phylogenetic analysis of DNA topoisomerase subunit B. (Top 6B).** The homologs were retrieved using COG1389 as described in the methods. Long branching sequences were excluded. Note that Asgard archaea sequences are placed in two different clades, as shown in the phylogeny of Top6A in Supplementary Figures 16,17. The alignment includes 966 sequences and 305 amino acid sites. The phylogeny is manually rooted. The analysis used the LG+C60+G+F+PMSF model, while the guide tree was inferred using the LG+F+G model. Scale bar: Average substitution per site.

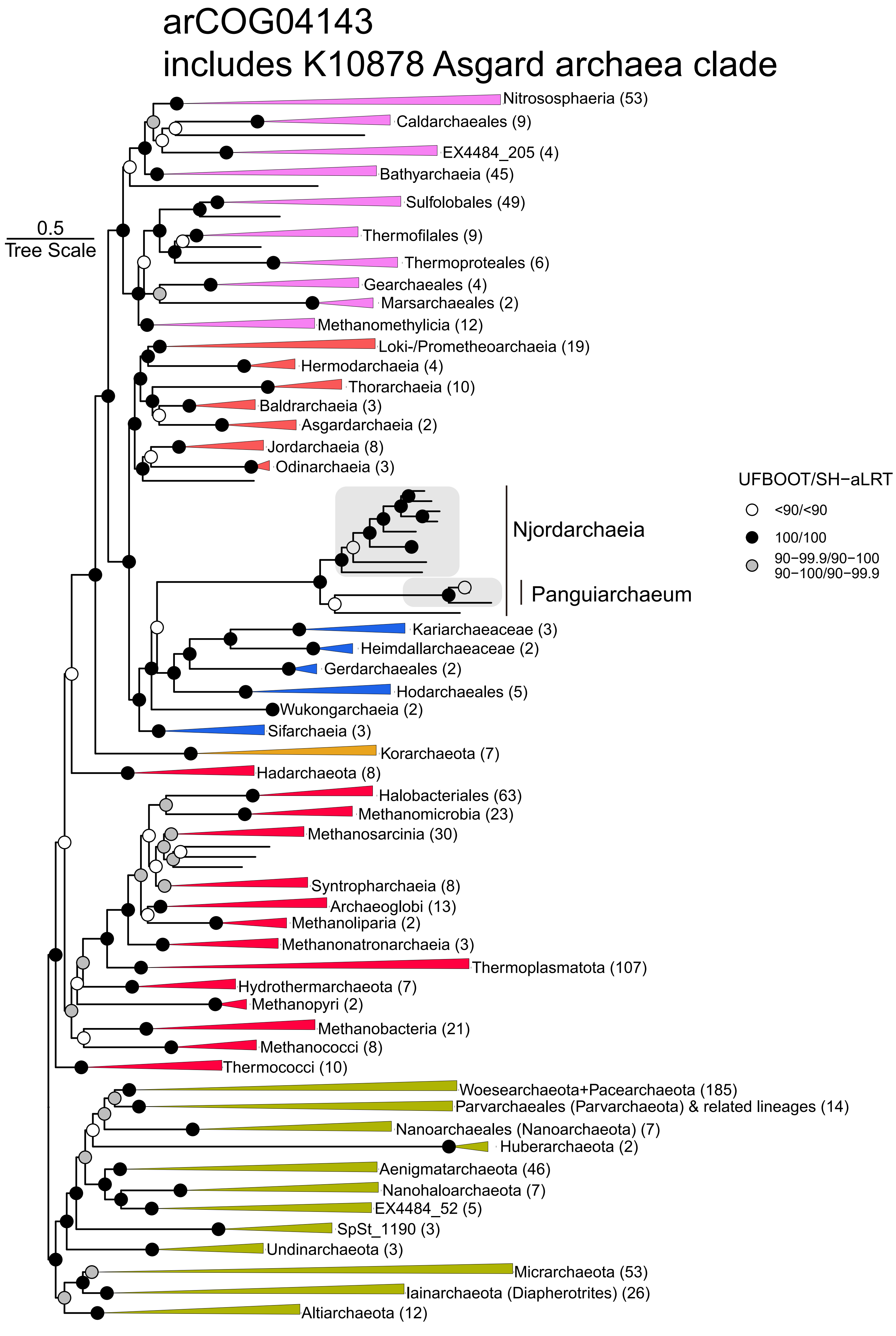


**Supplementary Figure 5. Maximum-likelihood phylogenetic analysis of the concatenated 50% top-ranked marker proteins (n=45), including DNA topoisomerase subunits, based on set 1 (966 archaeal taxa).** Note that the K10878 Asgard archaea sequences were included in this concatenation and led to the recovery of Njordarchaeia within Asgard archaea. The alignment includes 955 sequences and 10559 amino acid sites. The analysis used LG+C60+F+G+PMSF model with a guide tree based on LG+F+G. Scale bar: average substitutions per site. The number of taxa in each collapsed clade is shown by the number in parenthesis next to the clade name.

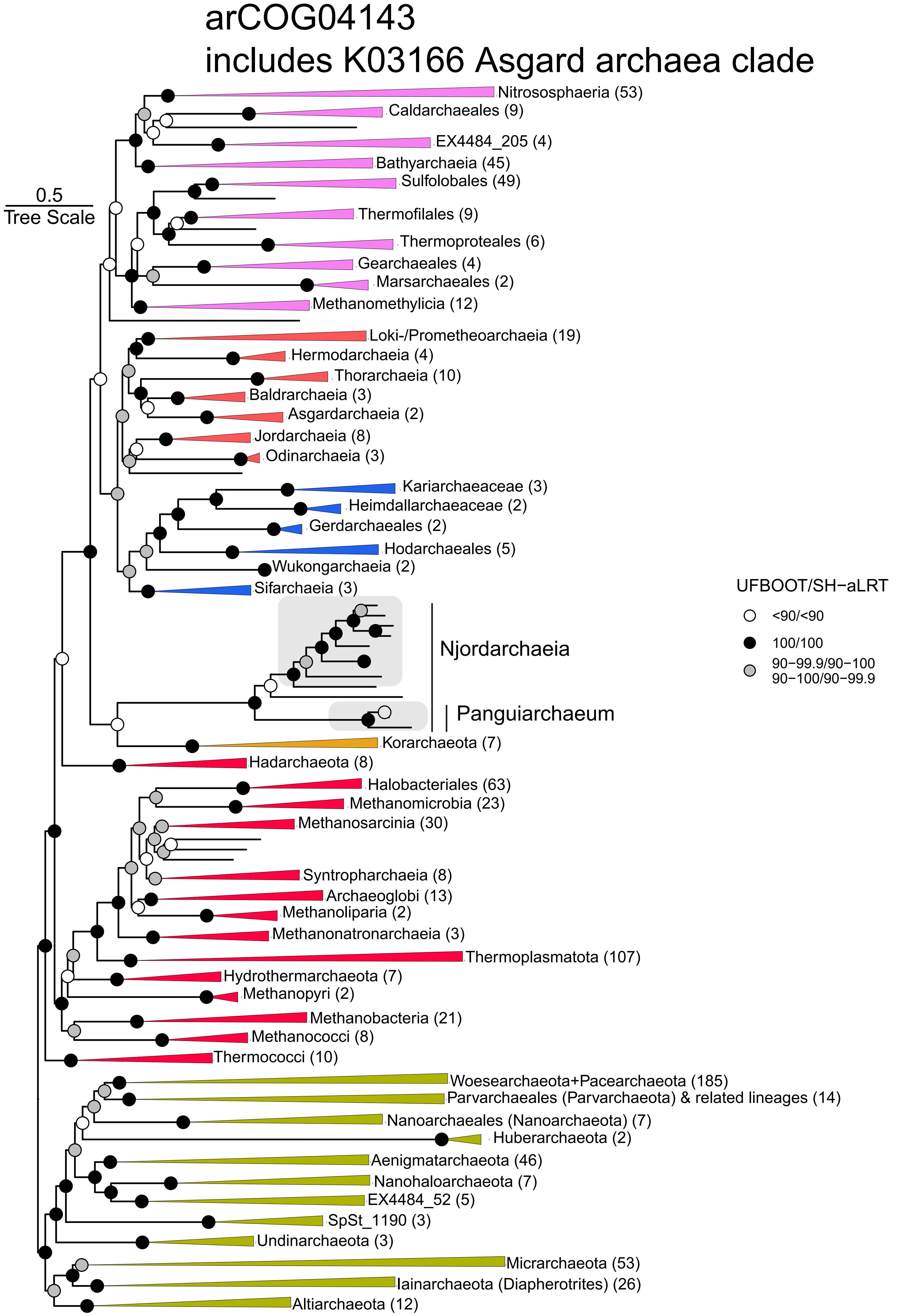


**Supplementary Figure 6. Maximum-likelihood phylogenetic analysis of the concatenated 50% top-ranked marker proteins (n=45), including DNA topoisomerase subunits, based on set 1 (966 archaeal taxa).** Note that the K03166 Asgard archaea sequences were included in this concatenation and led to the recovery of Korarchaeota topology. The alignment includes 955 sequences and 10573 amino acid sites. The analysis used LG+C60+G+F+PMSF model with a guide tree based on LG+F+G. Scale bar: average substitutions per site. The number of taxa in each collapsed clade is shown by the number in parenthesis next to the clade name.

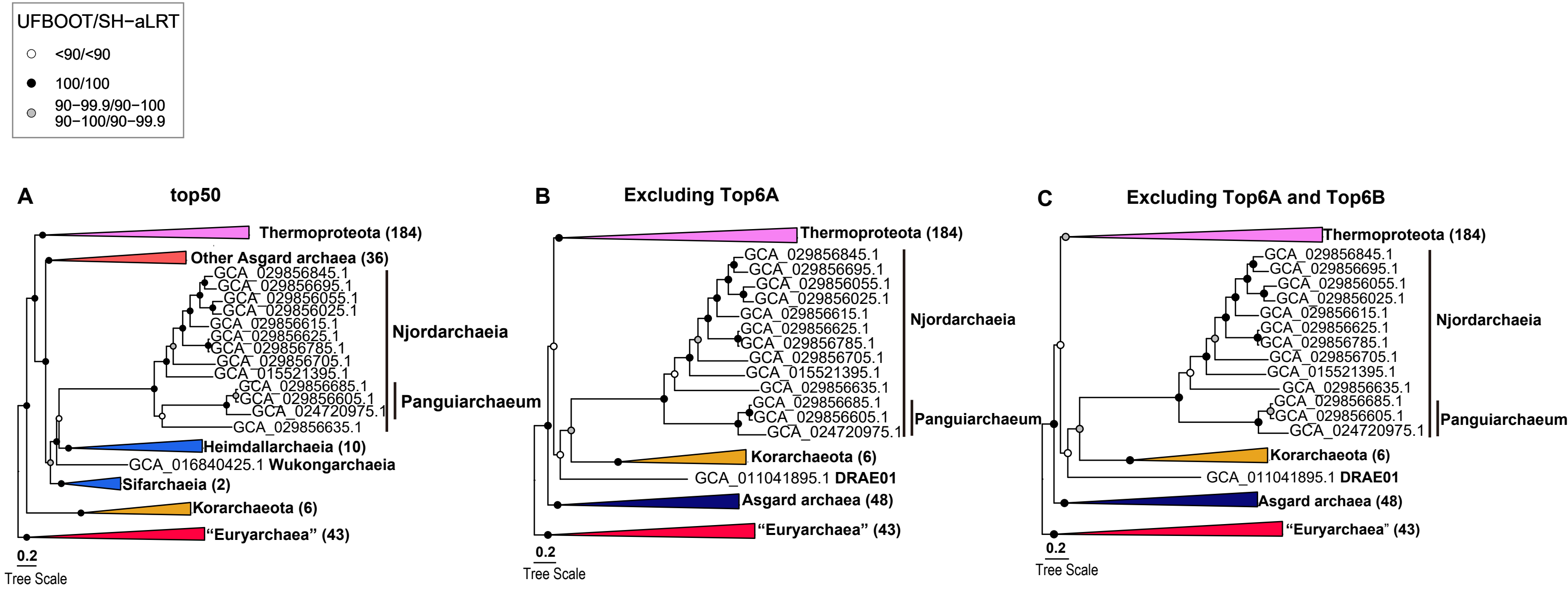


**Supplementary Figure 7. Maximum-likelihood phylogenetic analysis of the concatenated 50% top-ranked marker proteins based on set 2 (303 archaeal taxa), including DNA topoisomerase subunits (A, n=45, amino acid sites: 12687), excluding DNA topoisomerase subunit A (B, n=44, amino acid sites: 12372), and excluding DNA topoisomerase subunit AB (C, n=43, amino acid sites: 11971).** Note that this analysis reduces impacts from the guide tree and divergent DPANN archaeal sequences. The alignments contain 303 sequences. Phylogenetic analyses were performed using the LG+C60+G+F model. Scale bar: average substitutions per site. The number of taxa in each collapsed clade is shown by the number in parenthesis next to the clade name.

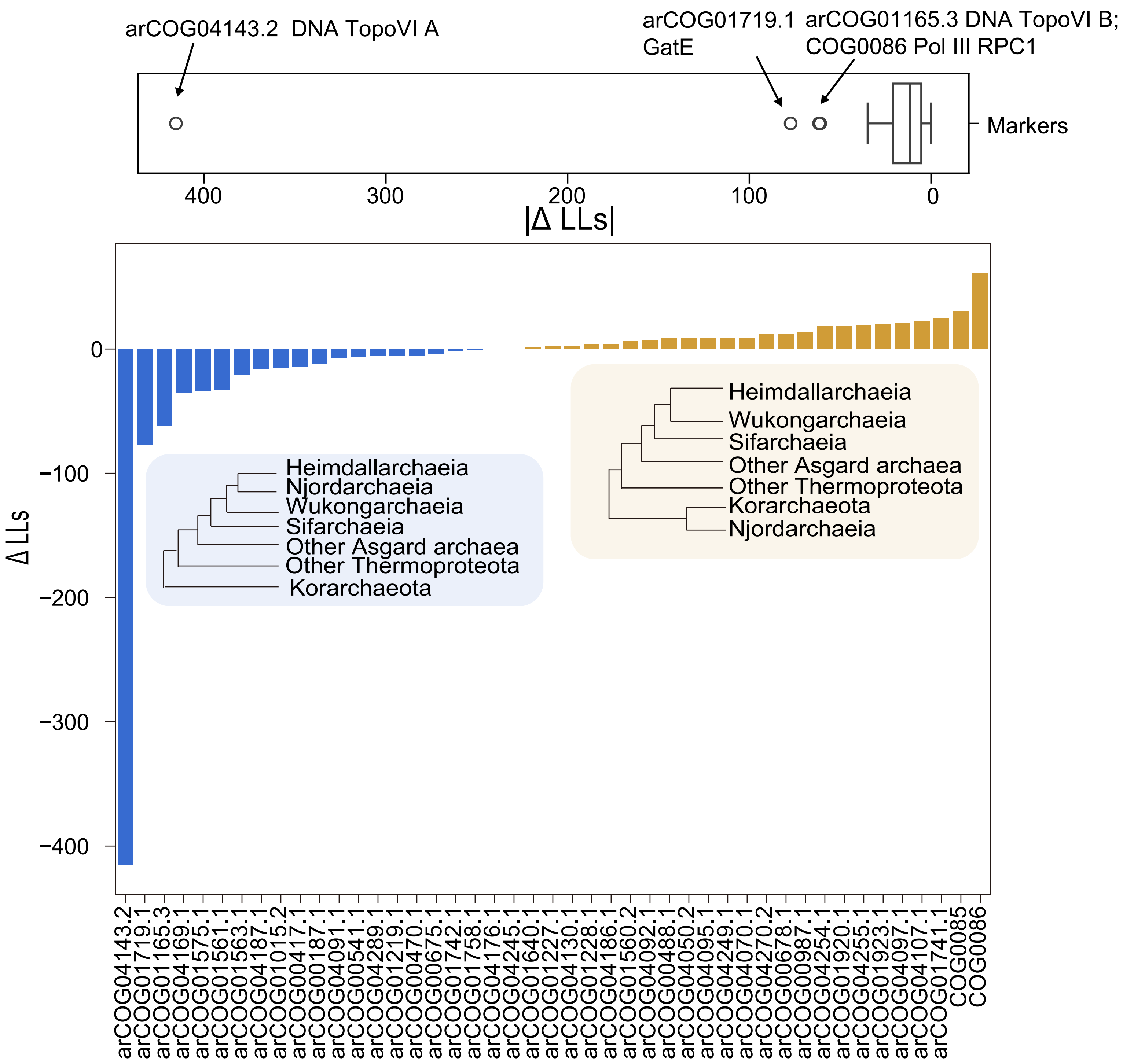


**Supplementary Figure 8. The gene-wise likelihood difference underlying the support for the Asgard archaea topology (Blue) and the Korarchaeota topology (Orange).** Note that arCOG04143 and arCOG01165 (DNA topoisomerase) ranked respectively as the first and third most influential markers on the phylogenetic inference. Constraint analysis was based on the concatenated top 50% ranked marker proteins and set 2, including 303 archaeal taxa. The distribution of the absolute likelihood difference (|ΔLLs|) of each marker is shown with a boxplot. The boxplot shows the median, the two hinges corresponding to the first and third quartiles, the whiskers corresponding to the 1.5X interquartile range from the hinges, and all individual points beyond the whiskers.

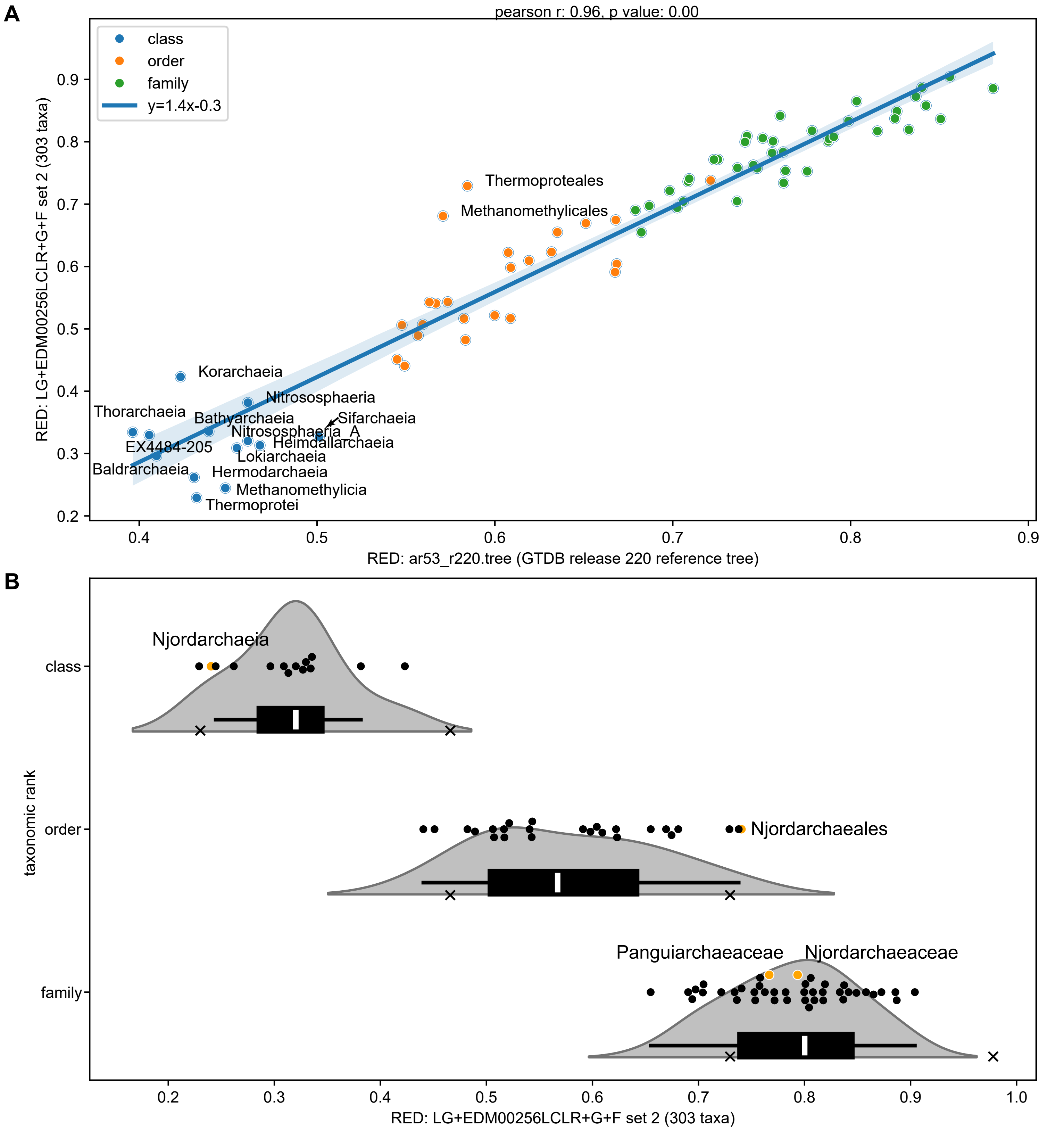


**Supplementary Figure 9. Rank normalisation using relative evolutionary divergence (RED) value. A**. Correlation of RED value of different class-, order- and family-level lineages in the inferred species tree in Figure 1B and GTDB release 220 reference tree, excluding Njordarchaeia lineages to be classified. The analysis shows the RED value of class-, order- and family-level lineages from Thermoproteota and Asgard archaea of these two trees are significantly correlated (Pearson r: 0.96, p-value: 0). Thermoproteales and Methanomethylicales are labelled for the deviation from the trend. **B**. Violin plot of RED values of class-, order- and family-level lineages in the inferred species tree. The density cure was shown with an inner box plot. White sticks denote the median. The lower and upper hinges of the inner boxplot correspond to the first and third quartiles. The upper/lower whiskers extend from the hinge to the largest/smallest value no further than 1.5 times the interquartile range. Individual data points are shown as scatter plots. Rank boundaries of GTDB release 220 are normalised to the inferred species tree and marked with crosses. RED values of Njordarchaeia, Njordarchaeales, Panguiarchaeaceae and Njordarchaeaceae are shown in orange. The RED value of Njordarchaeia was assigned to 0.2403, Njordarchaeales to 0.7421, Panguiarchaeaceae to 0.7667 and Njordarchaeaceae to 0.7935.

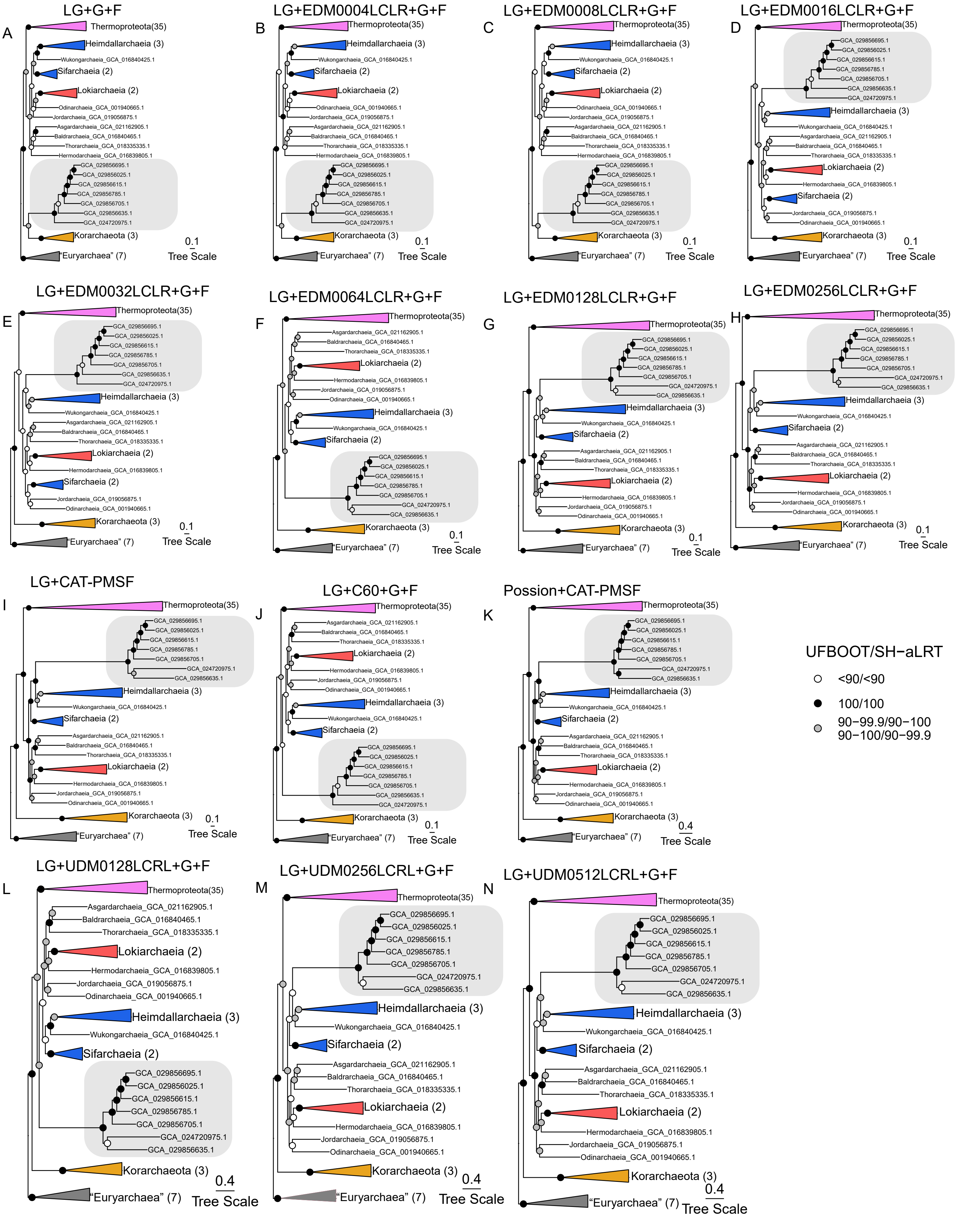


**Supplementary Figure 10. Maximum-likelihood (ML) phylogenetic analysis of the 50% top-ranked markers (n=43) and streamlined set 3 (71 taxa) based on various substitution models.** This alignment has 13345 amino acid sites. Scale bar: average substitutions per site. **A**. ML phylogeny based on site-homo-geneous substitution model (LG+G+F). ML phylogenetic tree inference using a custom site-heterogeneous substitution model with **B**. 4, **C**. 8, **D**. 16, **E**. 32, **F**. 64, **G**. 128 and **H**. 256 mixture categories. **I**. ML phylogenetic tree inference using LG+CAT-PMSF model. **J**. ML phylogenetic tree inference using LG +C60+G+F model. ML phylogenetic tree inference using **K**. LG+UDM0128+G+F, **L**. LG+UDM0256+G+F and **M**. LG+UDM0512+G+F models.

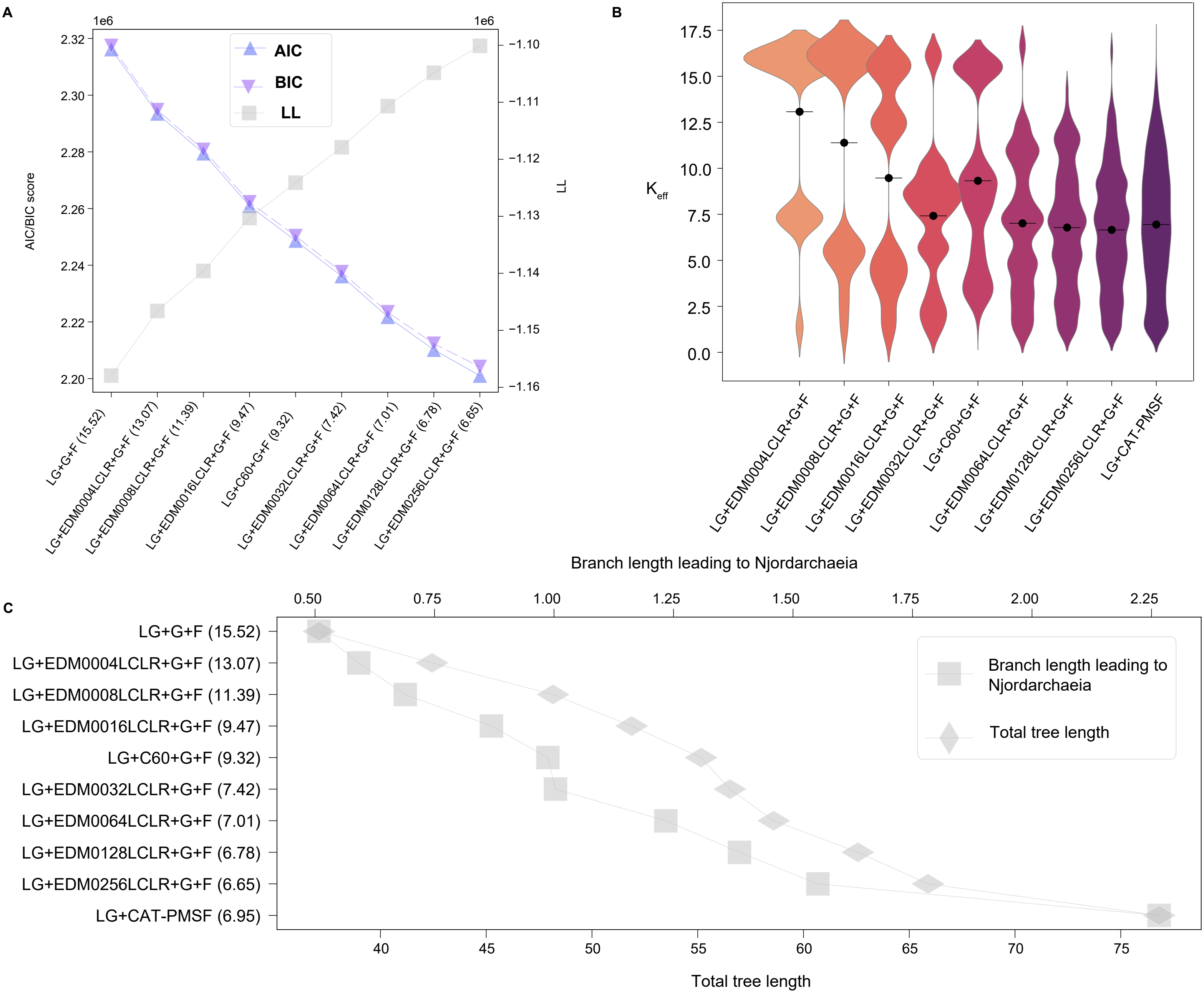


**Supplementary Figure 11. Performance of different substitution models on the alignment of 50% top-ranked marker proteins and 71 taxa. A**. Model fitness is measured using the Akaike information criterion (AIC) and Bayesian information criterion (BIC), as well as the likelihoods of different substitution models. The weighted effective amino acids are shown in parentheses. **B**. Distributions of the effective number of amino acids of the stationary distributions used by EDM models with different components, LG+CAT-PMSF and LG+C60+G+F model. The black bar denoted the weighted mean effective number of amino acids. **C**. The sum of all branch lengths (total branch length) and the length of the branch leading to the Njordarchaeia in the maximum-likelihood phylogenies is measured as the average number of inferred substitutions.

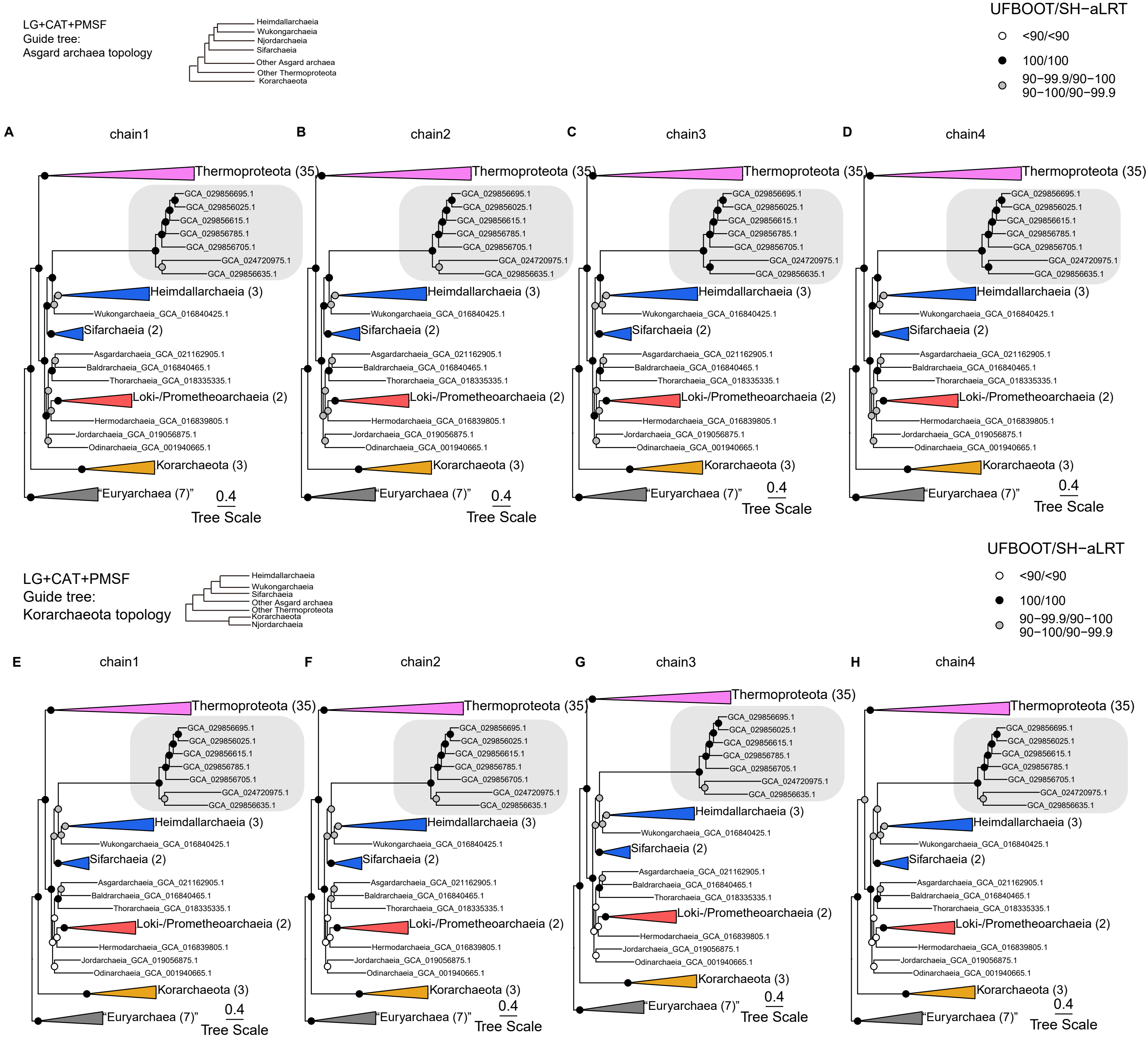


**Supplementary Figure 12. Maximum-likelihood phylogenetic analysis of the concatenated 50% top-ranked marker proteins based on set 3 (71 archaeal taxa), excluding DNA topoisomerase subunit AB (n=43).** Phylogenetic analyses were performed using the LG+CAT-PMSF model. Note that despite a different guide tree (**A-D**, Asgard archaea topology; **E-H,** Korarchaeota topology), all chains recovered the Asgard archaea topology with high support. Scale bar: average substitutions per site. The number of taxa in each collapsed clade is indicated by the number in parenthesis next to the clade name.

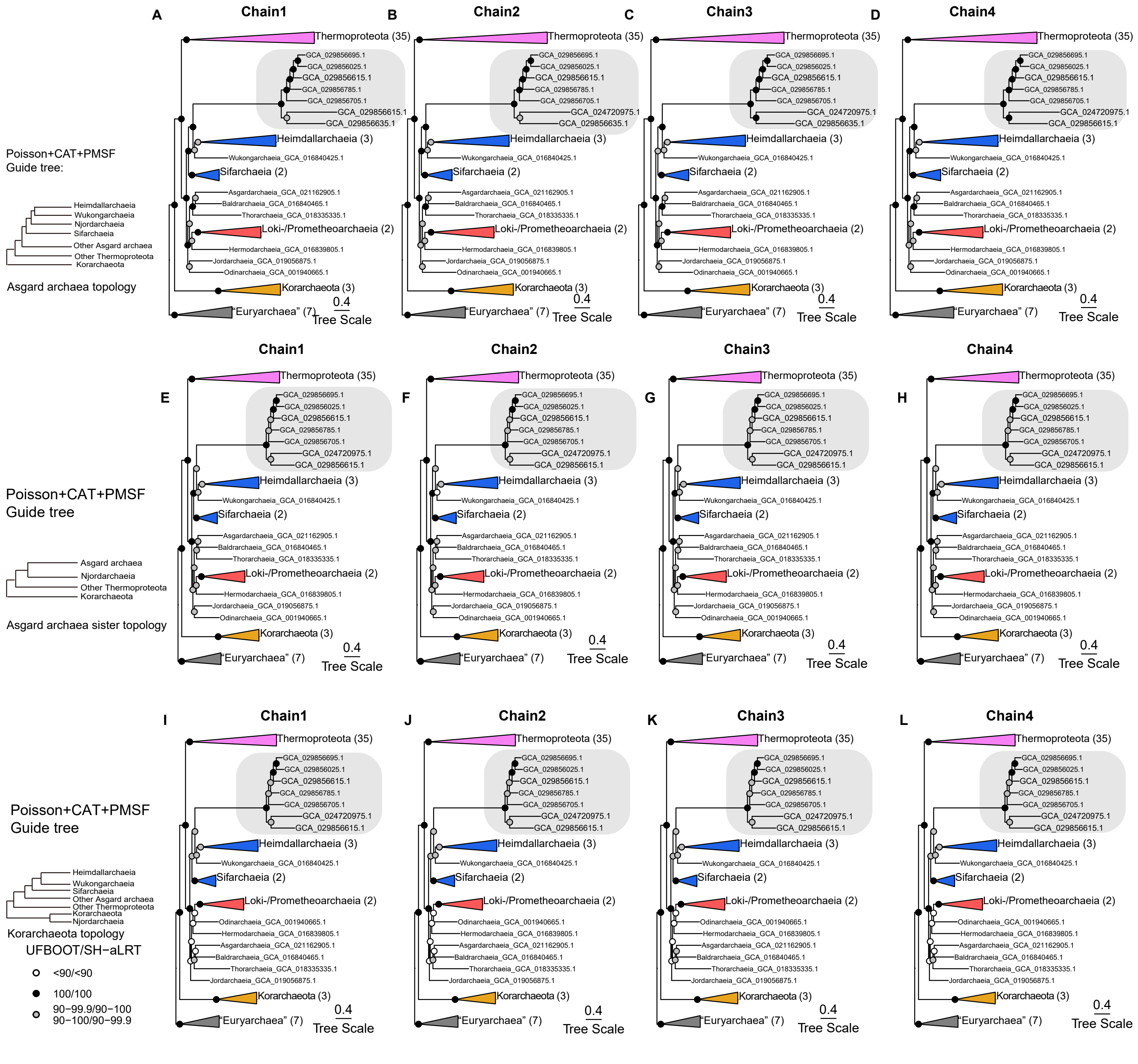


**Supplementary Figure 13. Maximum-likelihood phylogenetic analysis of the concatenated 50% top-ranked marker proteins based on set 3 (71 archaeal taxa), excluding DNA topoisomerase subunit AB (n=43).** Phylogenetic analyses were performed using the Poisson+CAT-PMSF model. Note that despite a different guide tree (**A-D**, Asgard archaea topology; **E-H**, Asgard archaea sister topology; **I-L**: Korarchaeota topology), site frequency extracted from all chains recovered the Asgard archaea topology with high support. Scale bar: average substitutions per site. The number of taxa in each collapsed clade is indicated by the number in parenthesis next to the clade name.

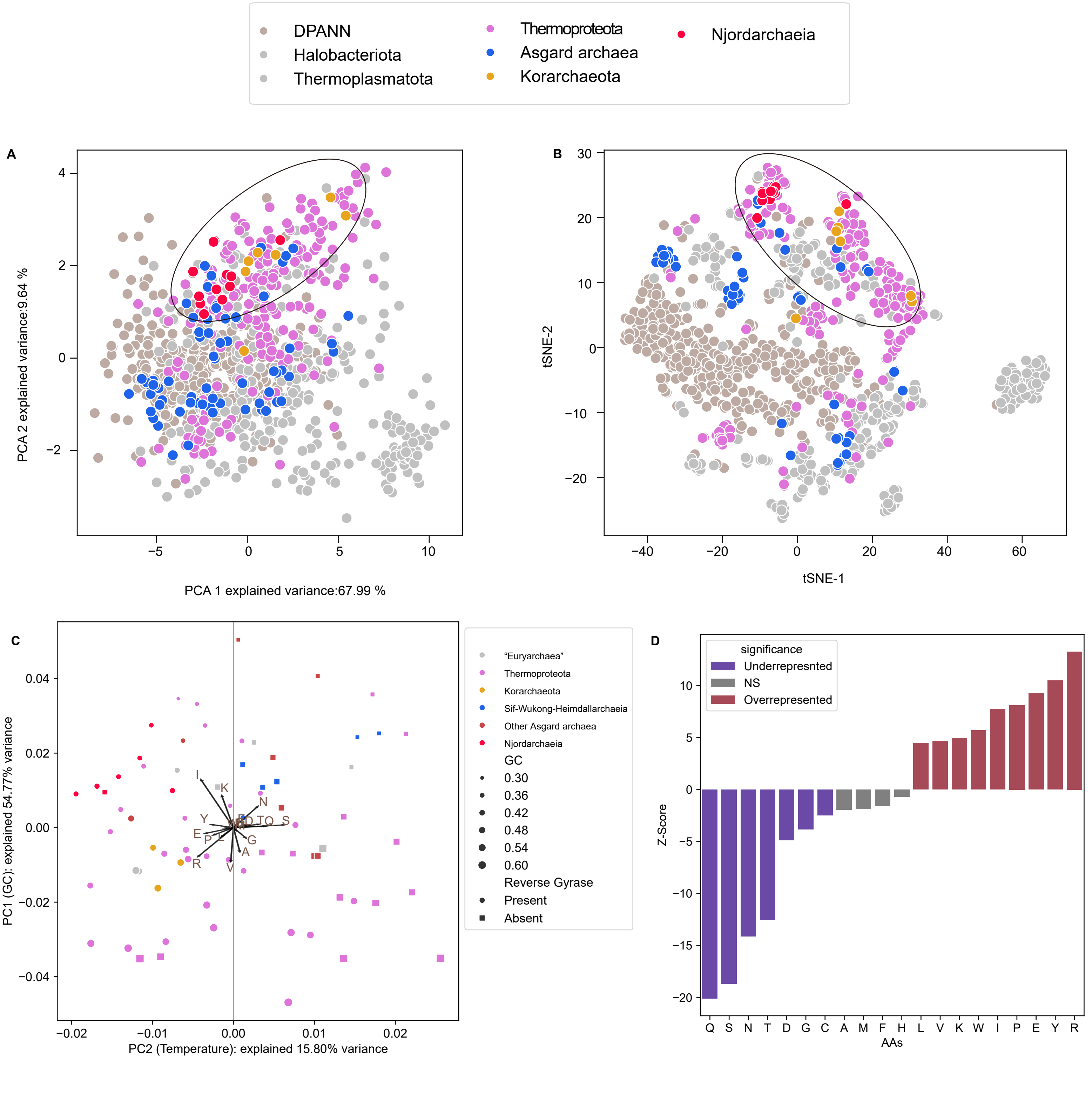


**Supplementary Figure 14. Amino acid composition of Njordarchaeia.** **A**. Principal component analysis (PCA) and **B**. t-distributed stochastic neighbour embed-ding (t-SNE) of the amino acid composition across Njordarchaeia (n=12), Halobacteriota (n=200), Thermoplasmatota (n=108), Thermoproteota (n=199), other Asgard archaea (n=67) and Korarchaeota proteome (n=7). **C**. Principal component analysis (PCA) of the amino acid composition of the 50% top-ranked markers. The correlation of each amino acid to a principal component (loadings) is shown with arrows. The presence and absence of reverse gyrase (i.e. arCOG01526) is indicated by a circle (present) and square (absent). The size of circles and squares corresponds to the GC content of the genomes. Note that axis 1 is associated with the GC content while axis 2 with the presence and absence of reverse gyrase. **D**. Enriched and depleted amino acids of (putative) thermophiles. A binominal test was conducted on the concatenation of 50% top-ranked marker proteins.

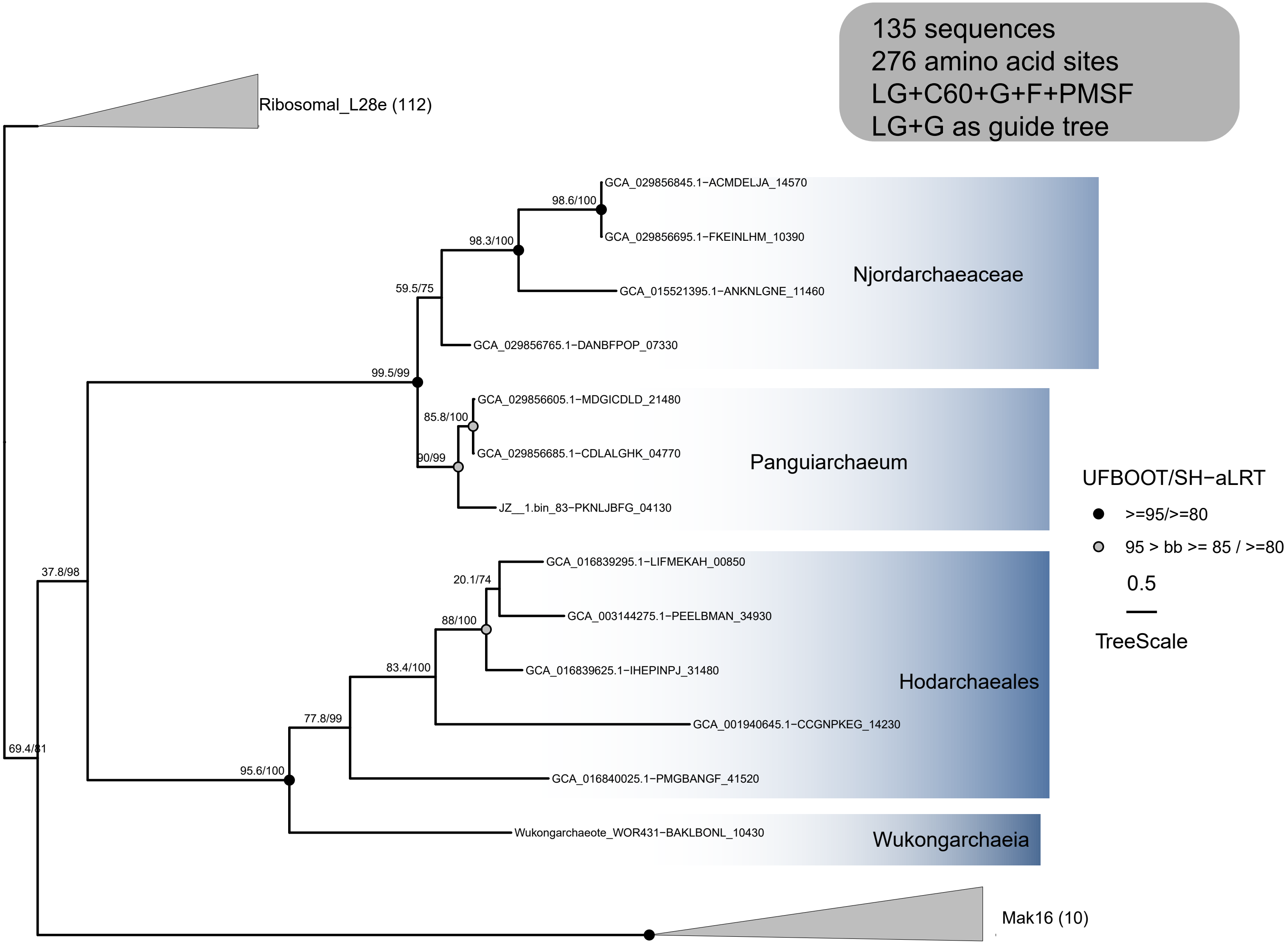


**Supplementary Figure 15. Phylogenetic analysis of ribosomal L28e/Mak16 homologs in Asgard archaea and Eukaryotes (LG+C60+G+F+PMSF).** The alignment comprises 276 amino acid positions and 135 sequences, including 13 Asgard archaea sequences, 10 Mak16 eukaryotic sequences and 112 ribosomal L28e sequences. The UFBOOT (left) and SH-aLRT support (right) values are shown on branches. The phylogenetic tree was rooted between the ribosomal L28e cluster and all other sequences. Scale bar: average substitution per site.

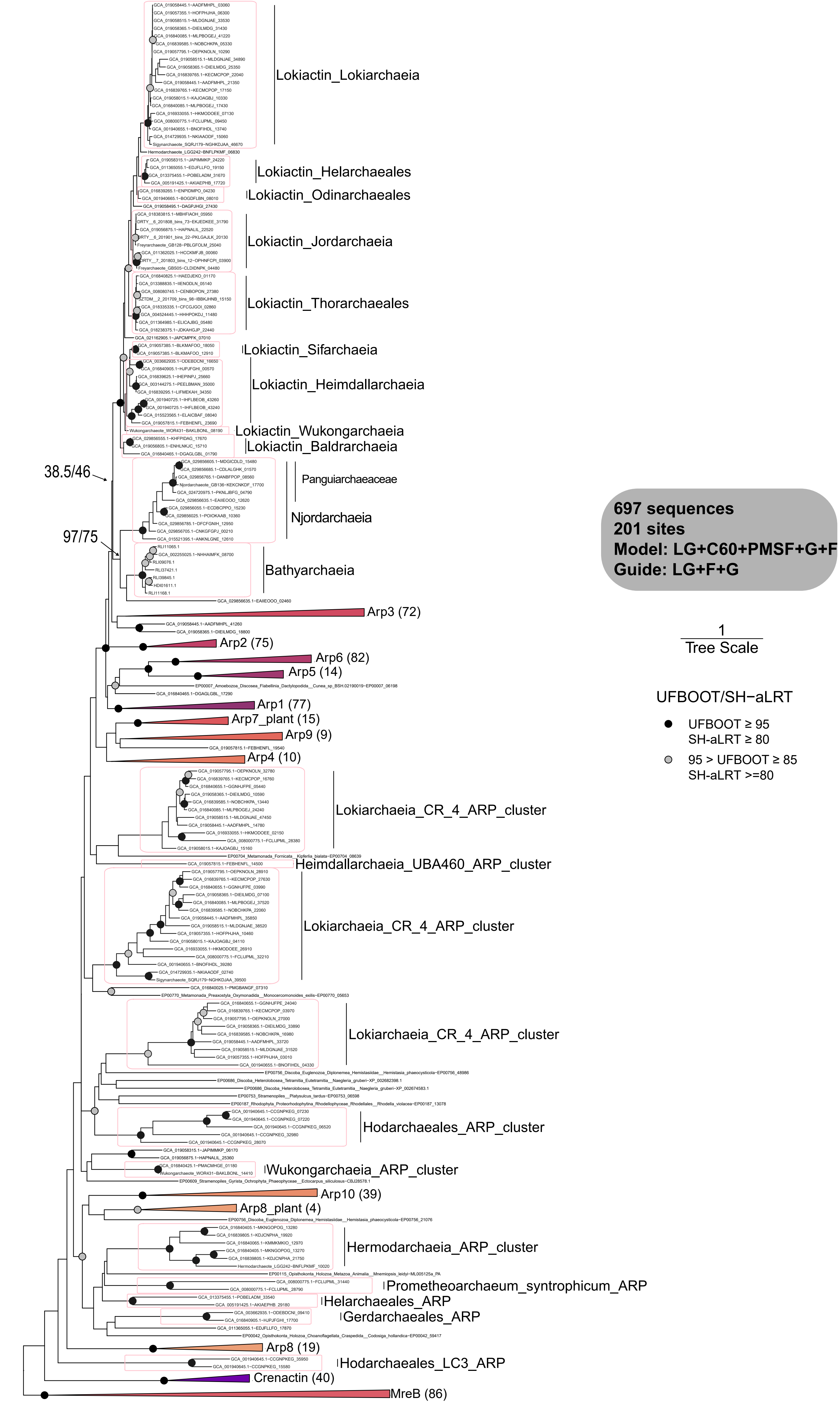


**Supplementary Figure 16. Phylogenetic analyses of Actin homologs in Asgard archaea, Thermoproteota and Eukaryotes (LG+C60+G+F+PMSF).** The alignment comprises 201 amino acid positions and 697 sequences. The UFBOOT (left) and SH-aLRT support (right) values are shown on branches. The phylogenetic tree is rooted between MreB sequences and the rest. The number of sequences is each clade is shown in parentheses. Scale bar: average substitution per site. ARP: Actin related protein. Note that bathyarchaeial sequences were selected from NCBI genbank database from comparison.

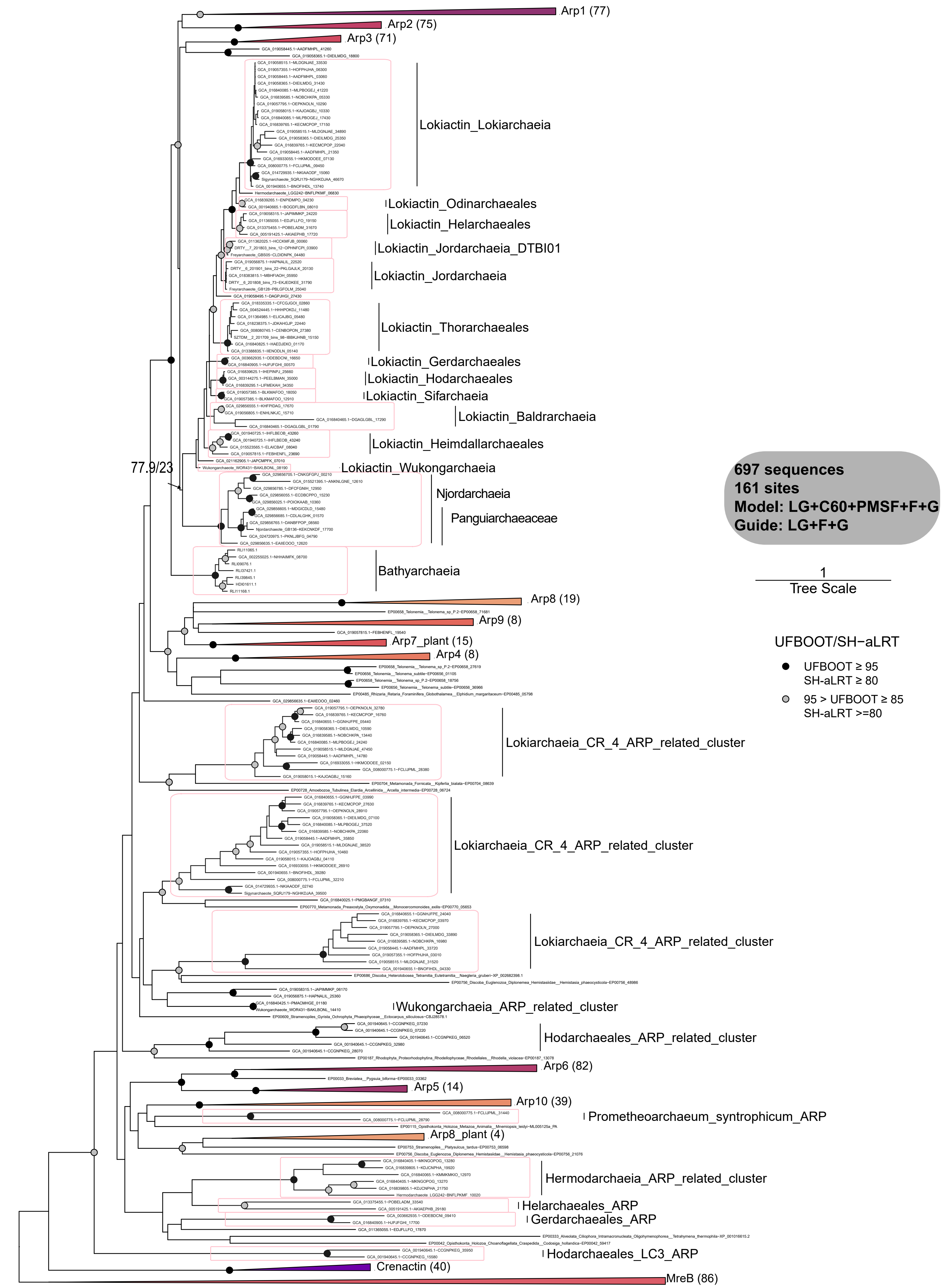


**Supplementary Figure 17. Phylogenetic analyses of Actin homologs in Asgard archaea, Thermoproteota and Eukaryotes (LG+C60+G+F+PMSF).** 20% of the most heterogeneous sites were removed from the alignment using chi2 test. The alignment comprises 161 amino acid positions and 697 sequences. The UFBOOT (left) and SH-aLRT support (right) values are shown on branches. The phylogenetic tree is rooted between MreB sequences and the rest. The number of sequences is each clade is shown in parentheses. Scale bar: average substitution per site. ARP: Actin related protein. Note that bathyarchaeial sequences were selected from NCBI genbank database from comparison.

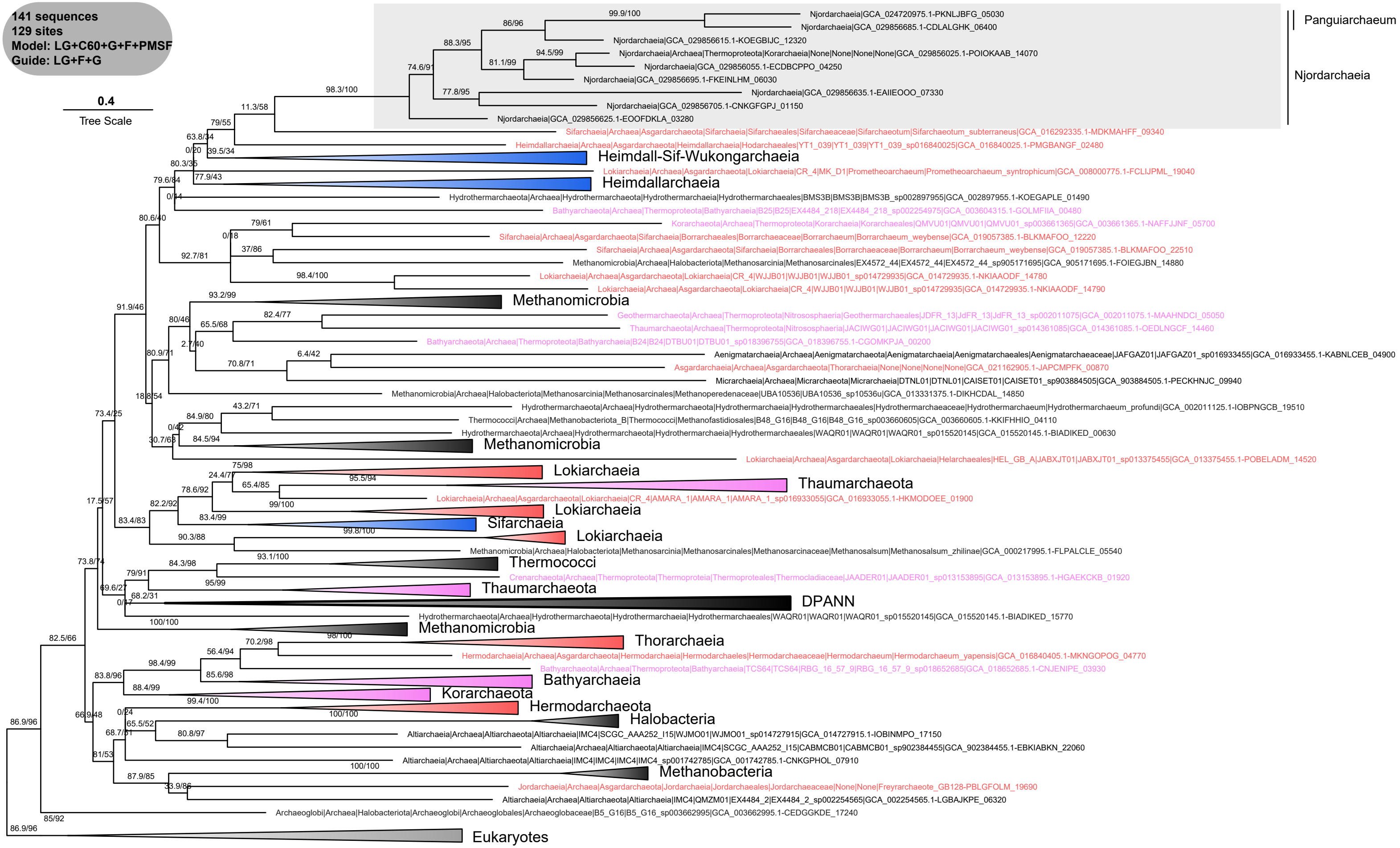


**Supplementary Figure 18. Phylogenetic analyses of translocon-associated protein (TRAP) complexes subunit beta.** The alignment includes 141 sequences and 129 amino acid sites. The phylogenetic inference was performed using the LG+C60+G+F+PMSF model with LG+F+G as a guide tree. The tree was rooted between eukaryotic sequences and the rest. Scale bar: average substitution per site.

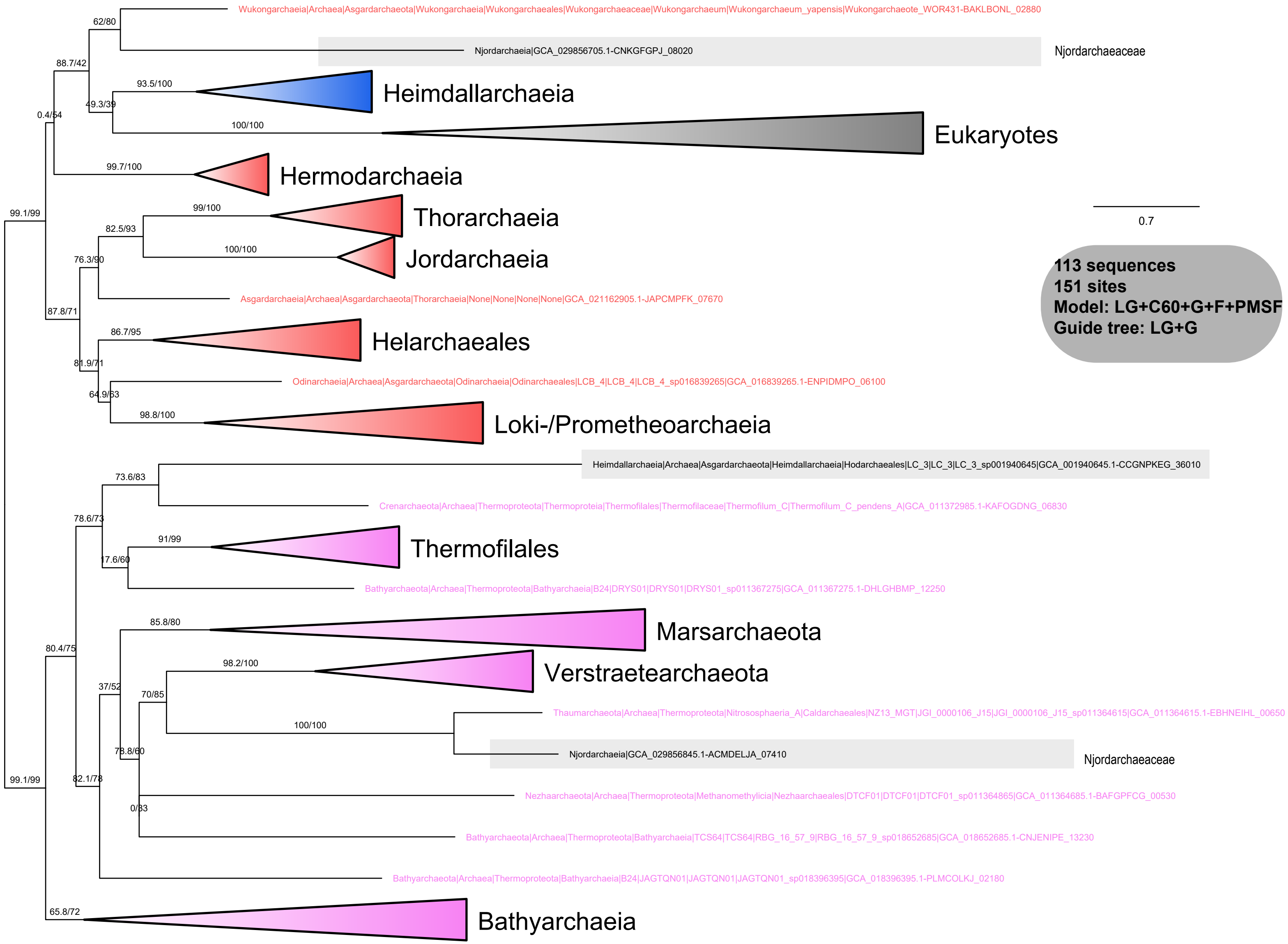


**Supplementary Figure 19. Phylogenetic analyses of OST3/OST6 proteins.** The alignment includes 113 sequences and 151 amino acid sites. The phylogeny is manually rooted at the thermoproteotal sequences. Note that Njordarchaeia sequences were shaded. The analysis used the LG+C60+G+F +PMSF model with a guide tree inferred using LG+G model. Scale bar: Average substitution per site.

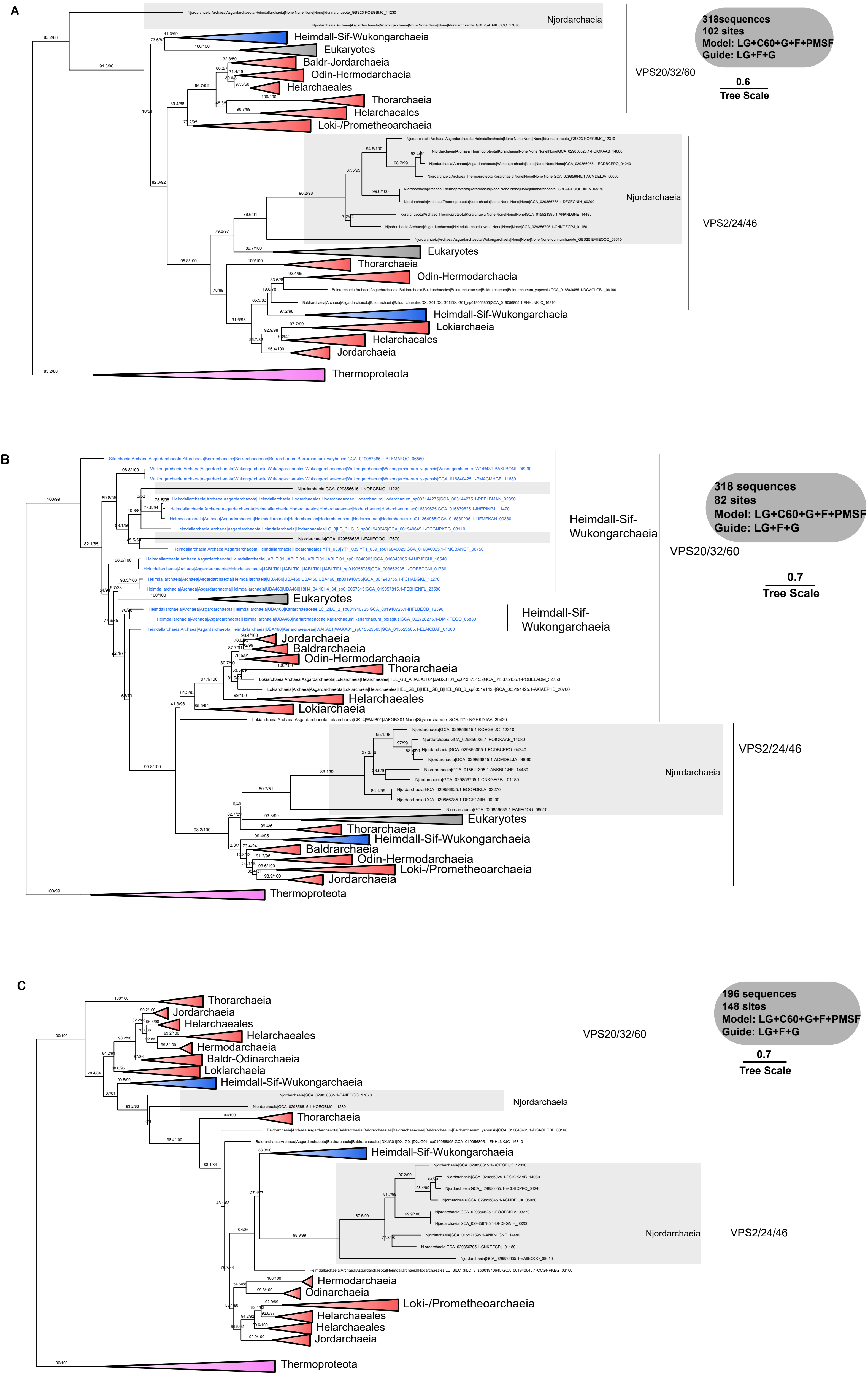


**Supplementary Figure 20. Phylogenetic analyses of SNF7 domain proteins. A**. Phylogenetic inference of SNF7 domain proteins from Thermoproteota, Asgard archaea and Eukaryotes (LG+C60+G+F+PMSF with LG+F+G for inferring the guide tree). The alignment includes 318 sequences and 102 amino acid sites. **B**. A second analysis was performed by removing 20% of the most heterogeneous sites, leaving 82 sites, using the LG+C60+G+F+PMSF model with LG+F+G for inferring the guide tree. **C**. Phylogenetic inference of SNF7 domain proteins from Thermoproteota and Asgard archaea. In this analysis, eukaryotic sequences were removed to alleviate potential long-branching attraction artefacts. This alignment includes 196 sequences and 148 amino acid sites. The analysis used the LG+C60+G+F+PMSF model and the LG+F+G model to infer the guide tree. The trees were manually rooted between Thermoproteota sequences and the rest. Scale bar: average substitution per site.

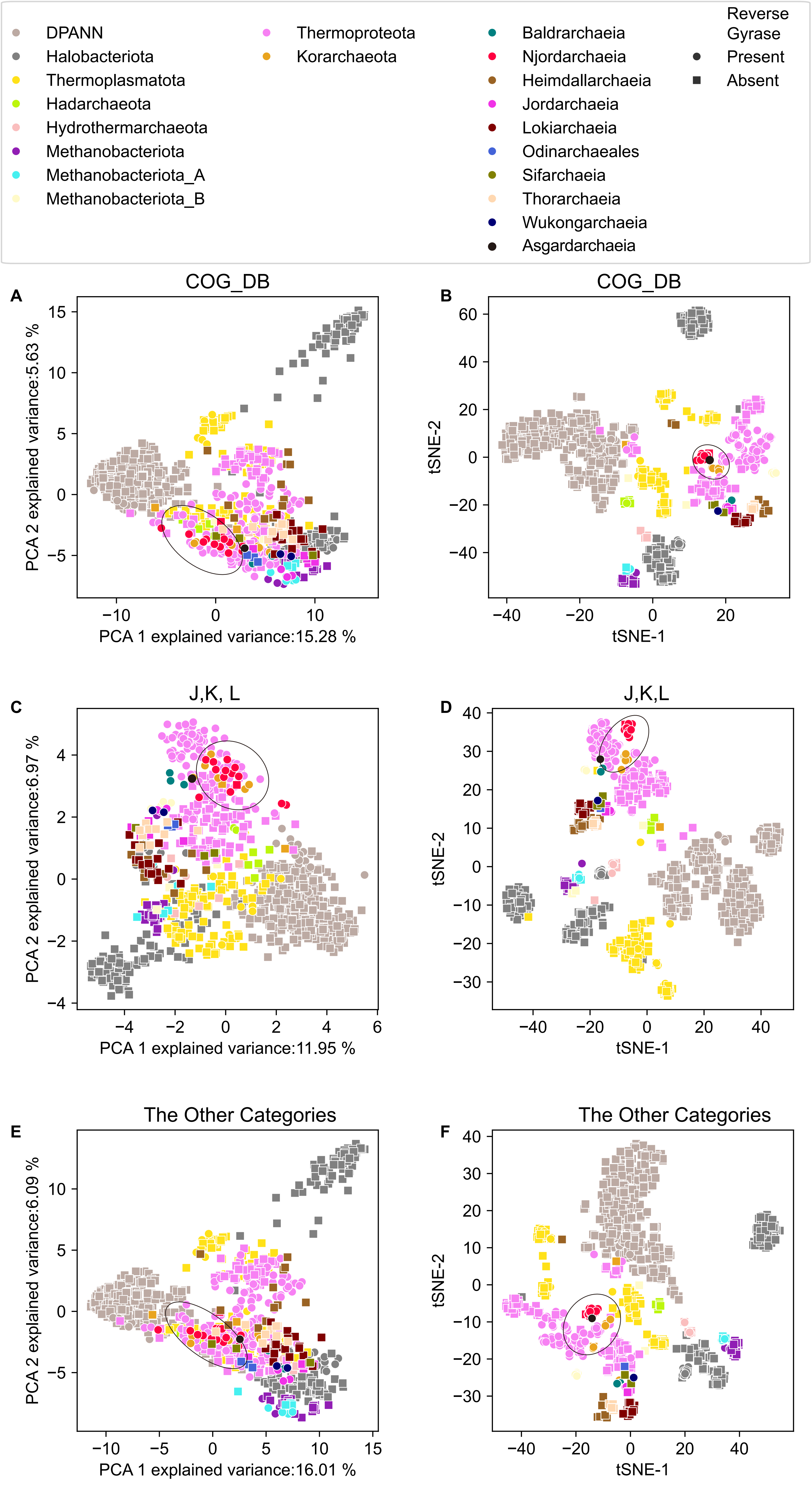


**Supplementary Figure 21. Comparison of presence and absence pattern of genes based on COG database.** Comparison based on the full COG database using PCA (**A**) and t-SNE algorithm (**B**). Comparison of presence and absence pattern of genes related to transcription (K), translation (J) and replication (L) using PCA (**C**) and t-SNE algorithm (**D**). Comparison of presence and absence pattern of genes related to other categories using PCA (**E**) and t-SNE algorithm (**F**).

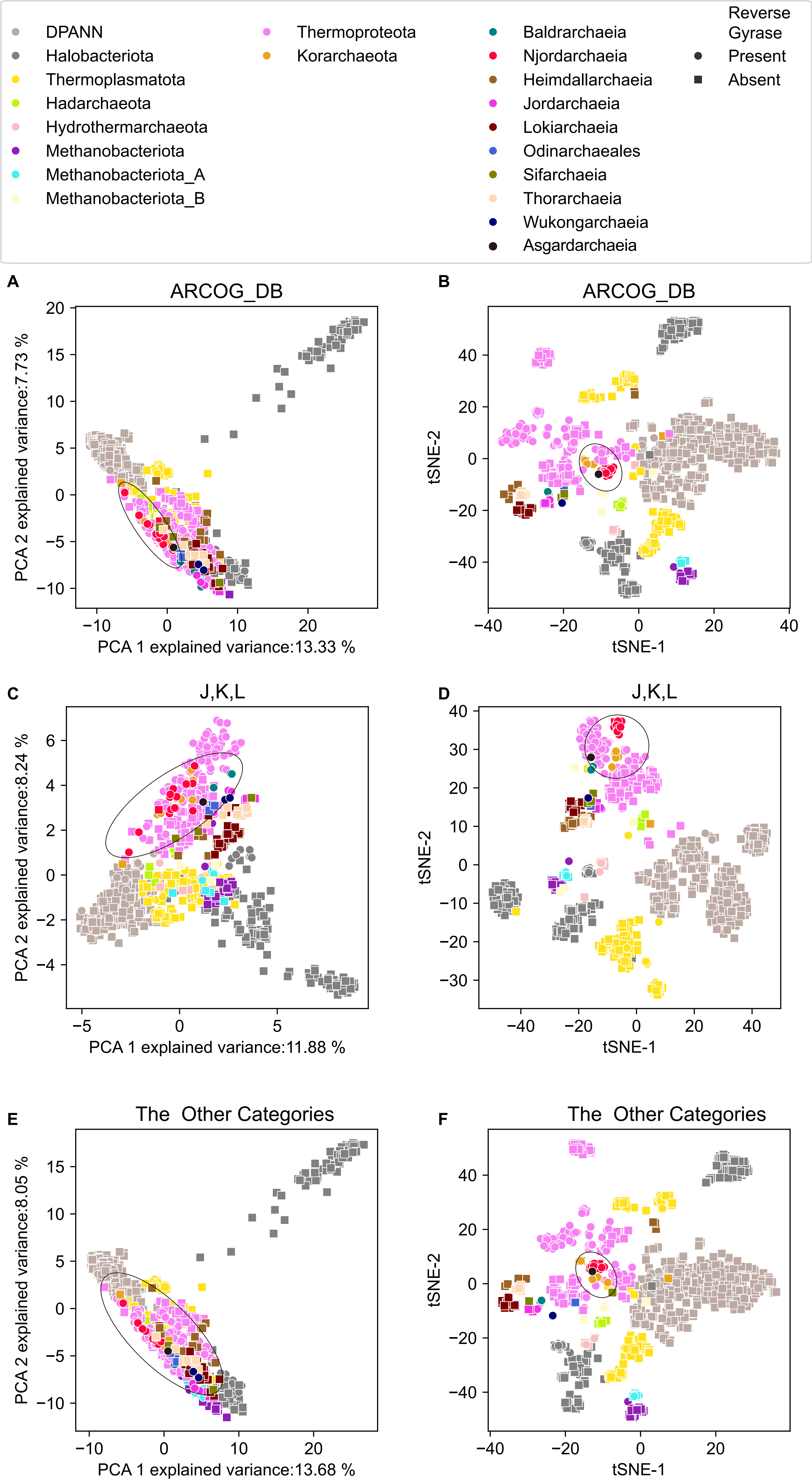


**Supplementary Figure 22. Comparison of presence and absence pattern of genes based on arCOG database.** Comparison based on the full arCOG database using PCA (**A**) and t-SNE algorithm (**B**). Comparison of presence and absence pattern of genes related to transcription (K), translation (J) and replication (L) using PCA (**C**) and t-SNE algorithm (**D**). Comparison of presence and absence pattern of genes related to other categories using PCA (**E**) and t-SNE algorithm (**F**).

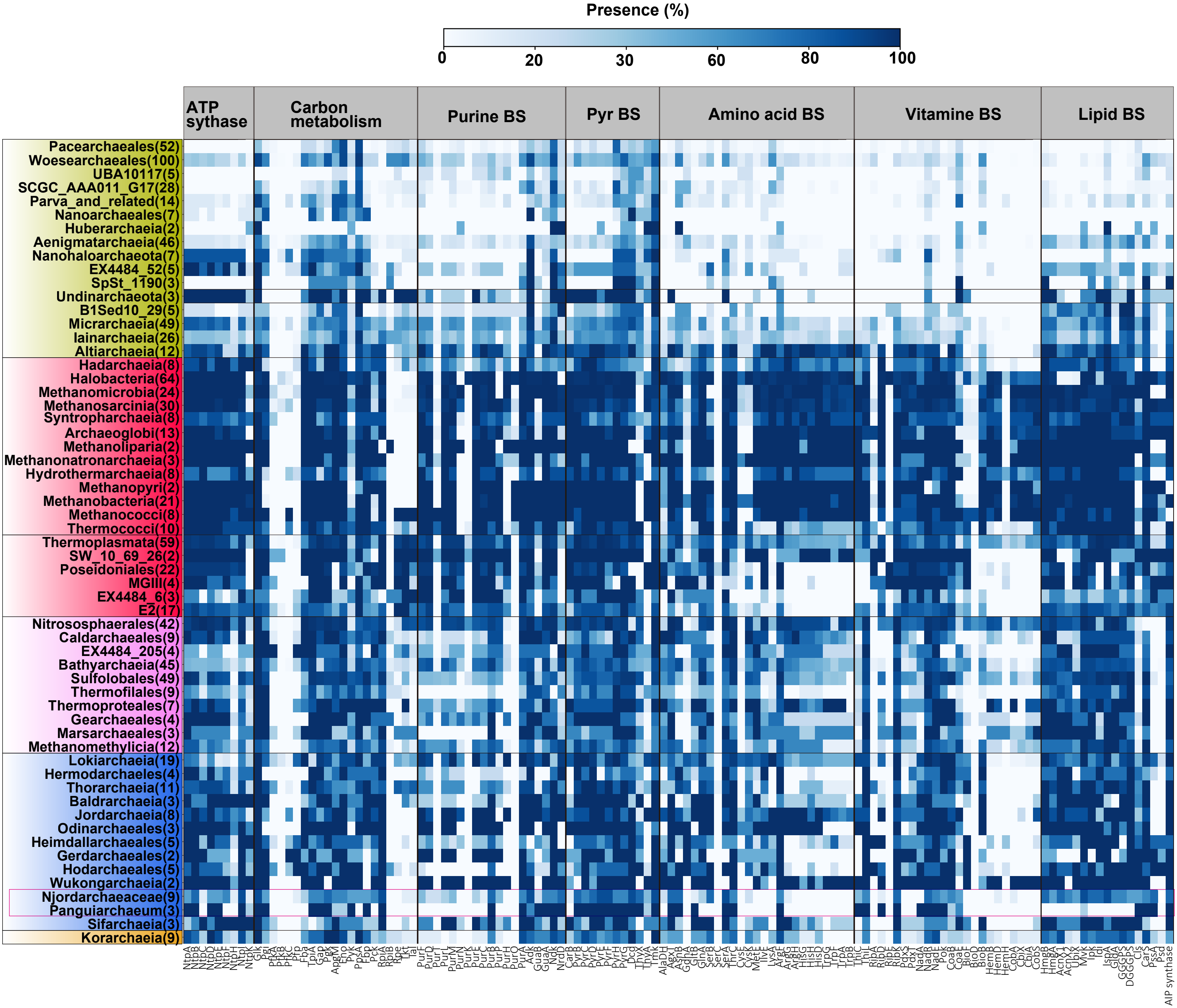


**Supplementary Figure 23. Presence of key metabolic proteins across major archaeal lineages.** Protein family presence was calculated by detecting key proteins of interest across 966 archaeal genomes and calculating the occurrence in percentage across the total number of genomes included in each phylogenetic cluster based on a presence/absence table. Only clusters with more than 2 genomes are shown. Number in parentheses: number of genomes included for each taxon of interest. BS: Biosynthesis.

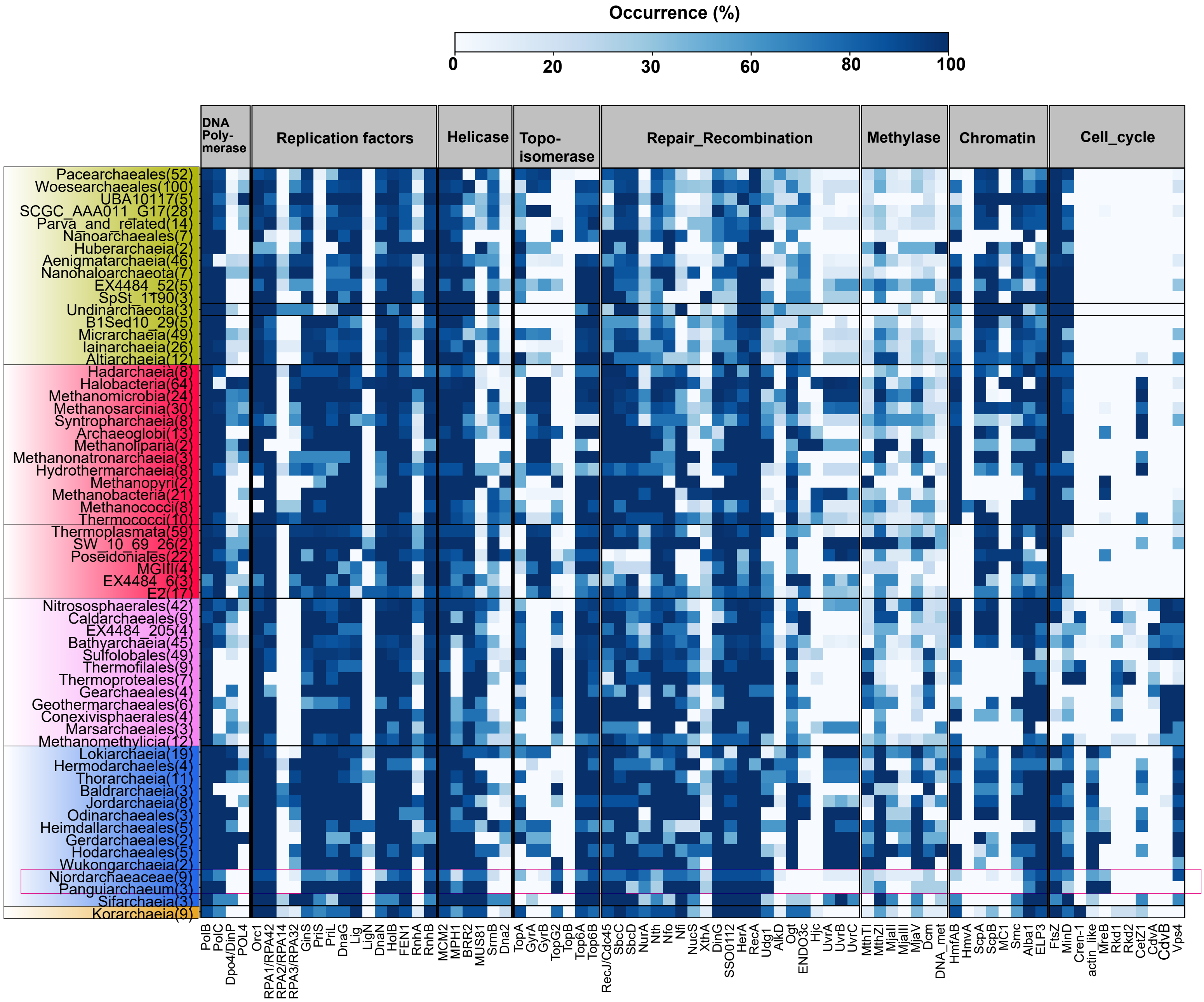


**Supplementary Figure 24. Occurrence of informational processing gene across major archaeal lineages.** Protein occurrence was calculated by detecting key proteins of interest across 966 archaeal genomes and calculating the occurrence in percentage across the total number of genomes included in each phylogenetic cluster based on a presence/absence table. Only clusters with more than two genomes are shown here.

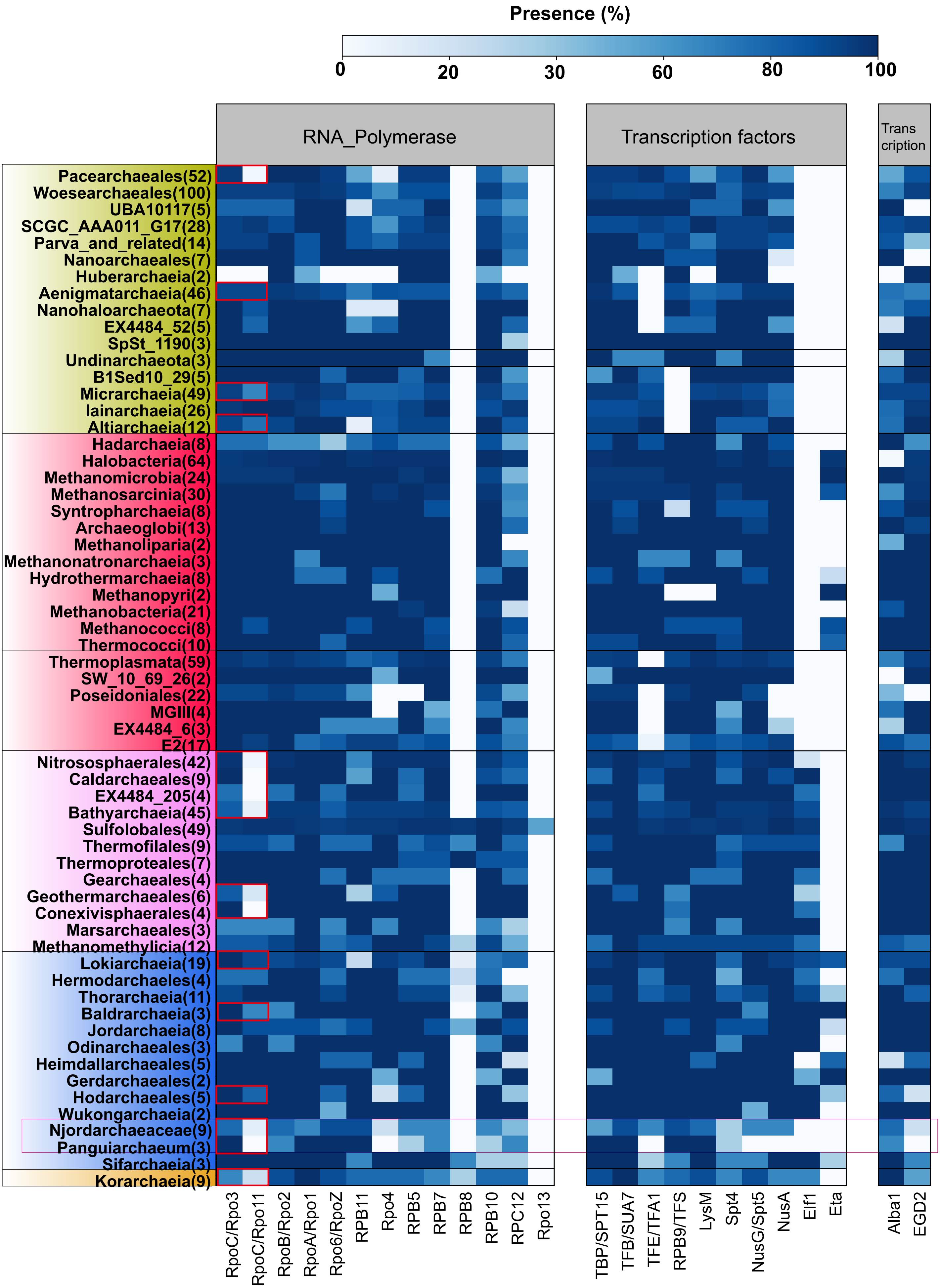


**Supplementary Figure 25. Presence of transcription genes across major archaeal lineages.** Protein presence was calculated by detecting key proteins of interest across 966 archaeal genomes and calculating the occurrence in percentage across the total number of genomes included in each phylogenetic cluster based on a presence/absence table. Only clusters with more than two genomes are shown here. Red circles indicated that A fused DNA-directed RNA polymerase was identified in the genome of this lineage.

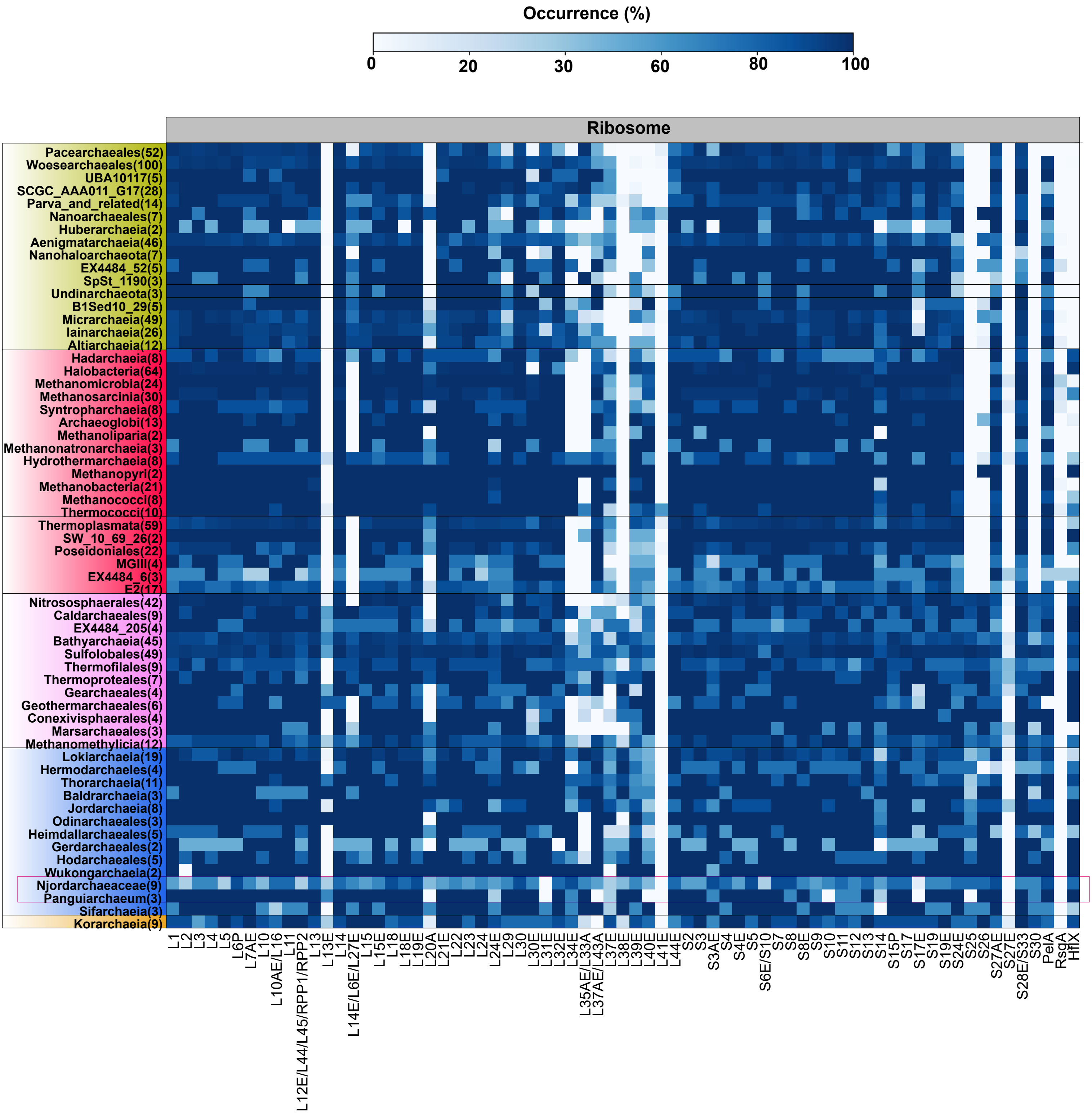


**Supplementary Figure 26. Occurrence of ribosomal genes across major archaeal lineages.** Protein occurrence was calculated by detecting key proteins of interest across 966 archaeal genomes and calculating the occurrence in percentage across the total number of genomes included in each phylogenetic cluster based on a presence/absence table. Only clusters with more than two genomes are shown here.

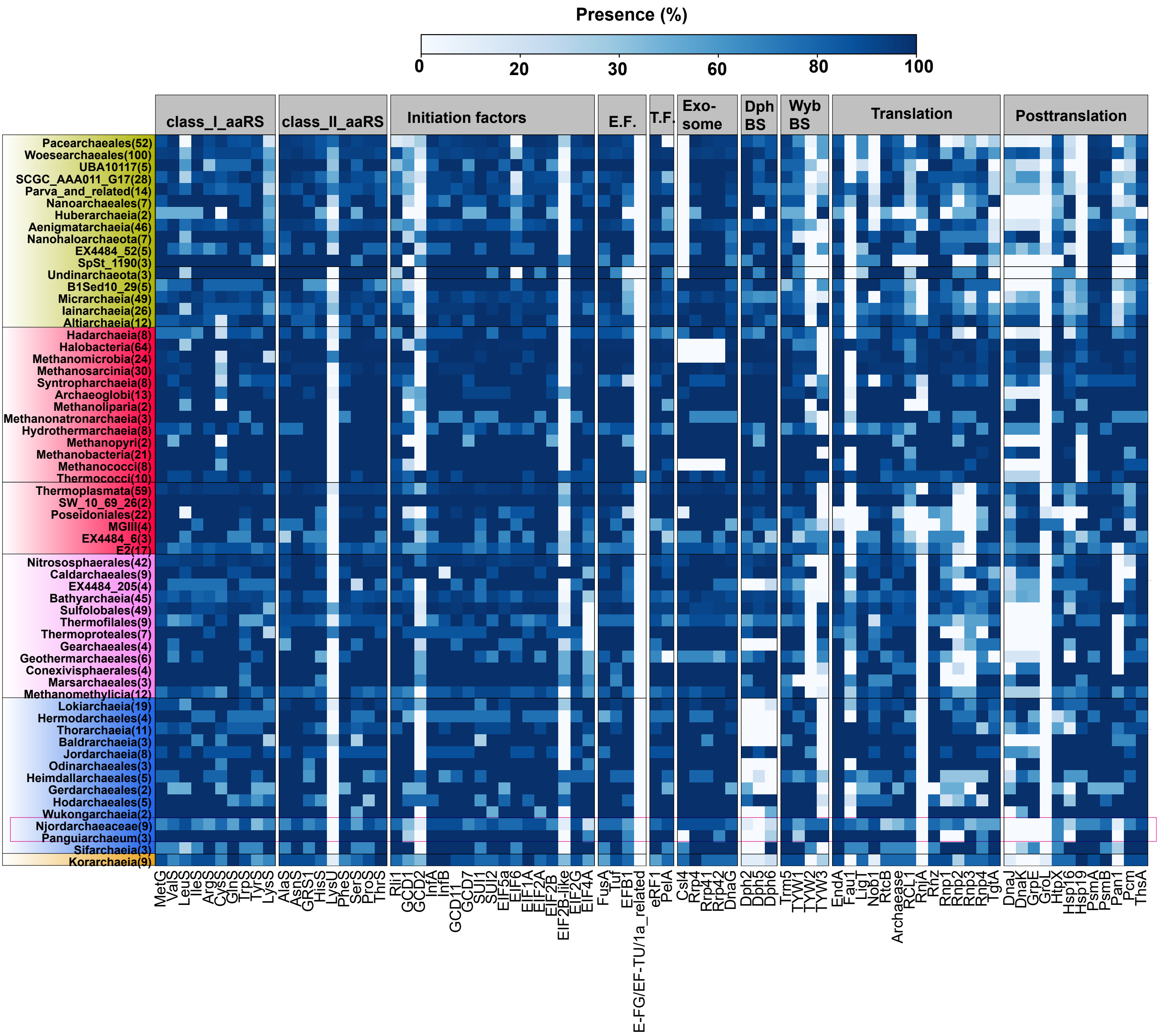


**Supplementary Figure 27. Presence of translation-related genes across major archaeal lineages.** Protein presence was calculated by detecting key proteins of interest across 966 archaeal genomes and calculating the occurrence in percentage across the total number of genomes included in each phylogenetic cluster based on a presence/absence table. Only clusters with more than two genomes are shown here.

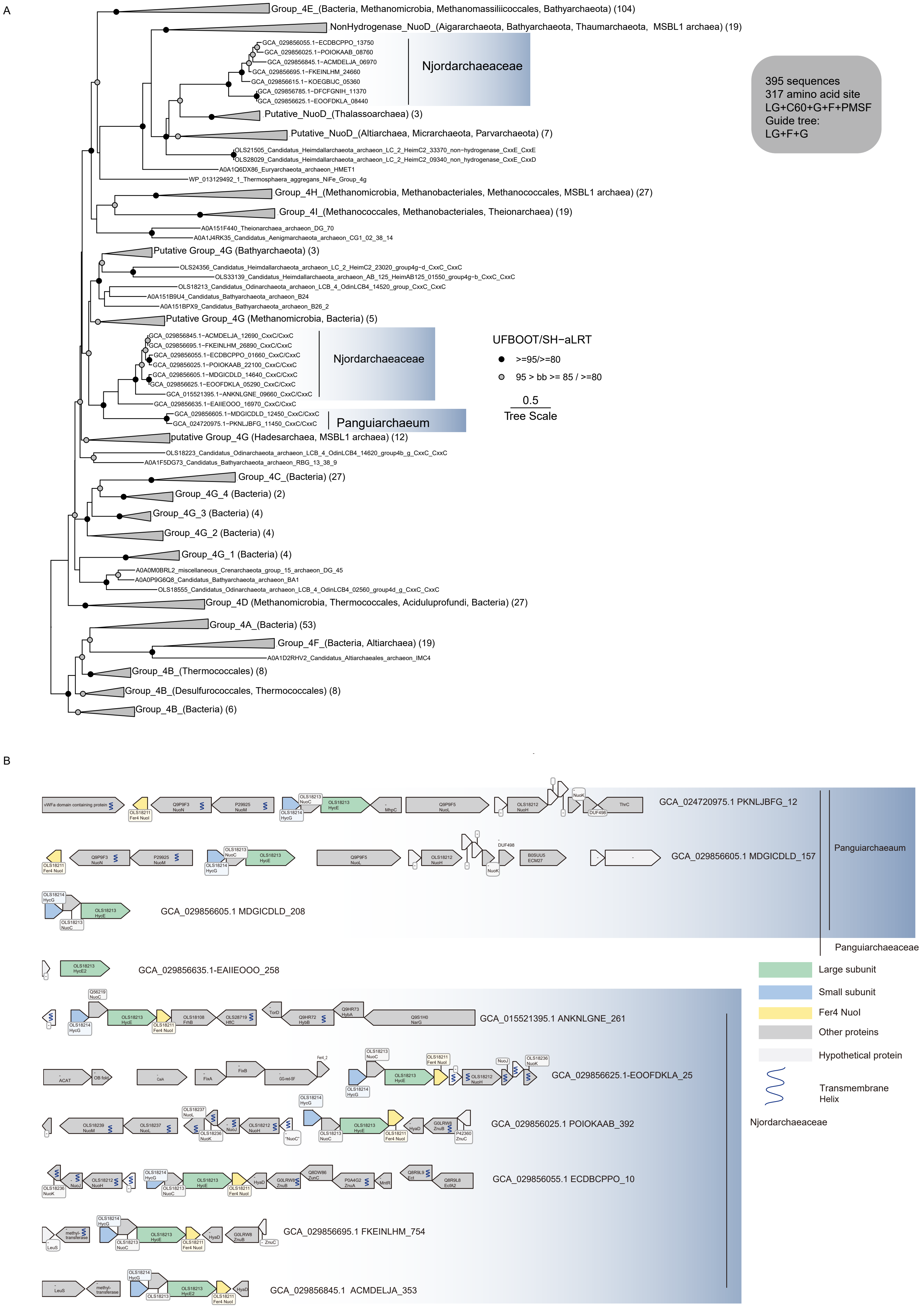


**Supplementary Figure 28. Structure of [NiFe] hydrogenase group 4 gene clusters and phylogenetic analysis of its large subunit in Njordarchaeia. A**. Maximum-likelihood phylogenetic analysis of the large subunit of group 4 [NiFe] hydrogenases. N-terminal and C-terminal CxxC motifs that ligate $H_2$-binding metal centres are annotated in the Njordarchaeia homologs. The phylogeny of the large subunit of group 4 [NiFe] hydrogenase was rooted manually. The alignment has 395 sequences with 317 amino acid sites. The analysis was performed using the LG+C60+G+F+PMSF model with a guide tree inferred using the LG+F+G model. Scale bar: Average substitution per site. **B**. Structure of [NiFe] hydrogenase group 4 gene clusters. Neighbouring genes of the large subunits of [NiFe] group 4g hydrogenase located in the same contig were shown (Maximum: 10 downstream and 10 upstream). All genes are drawn to scale. Key abbreviations are as follows. Nuo: NAHD dehydrogenase. Hyc: Hydrogenase component. Hya: [NiFe] hydrogenase maturation factor. MhpC: Alpha/beta superfamily hydrolase. ECM27: Ca2+/Na+ antiporter. ACAT: Acetyl-CoA acetyltransferase. CaiA: Acyl-CoA dehydrogenase. Fix: Electron transfer flavoprotein. LeuS: Leucyl-tRNA synthetase. Znu: ABC-type Mn/Zn transport system. Ecf: Energy-coupling factor transporter. MntR: Mn-dependent transcriptional regulator. NarG: Nitrate reductase alpha subunit. Detailed annotations can be found in Supplementary Data 8 and 10.

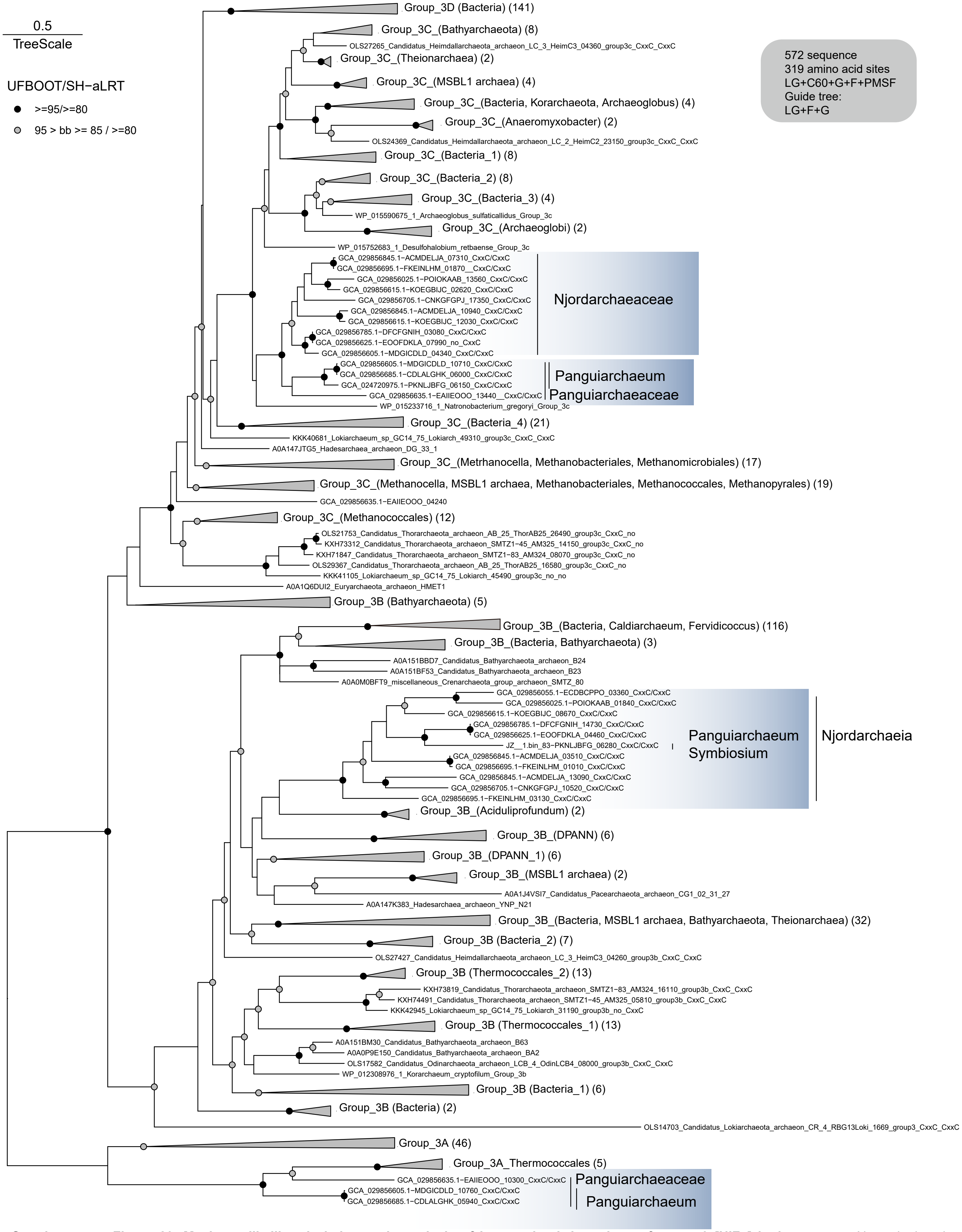


**Supplementary Figure 29. Maximum-likelihood phylogenetic analysis of large subunit homologs of group 3 [NiFe] hydrogenases.** N-terminal and C-terminal CxxC motifs that ligate $H_2$-binding metal centres are annotated in the Njordarchaeia homologs. The phylogeny of the large subunit of group 3 [NiFe] hydrogenase was rooted manually. The alignment has 572 sequences with 319 amino acid sites. The analysis was performed using the LG+C60+G+F+PMSF model with a guide tree inferred using the LG+F+G model. Scale bar: Average substitution per site.

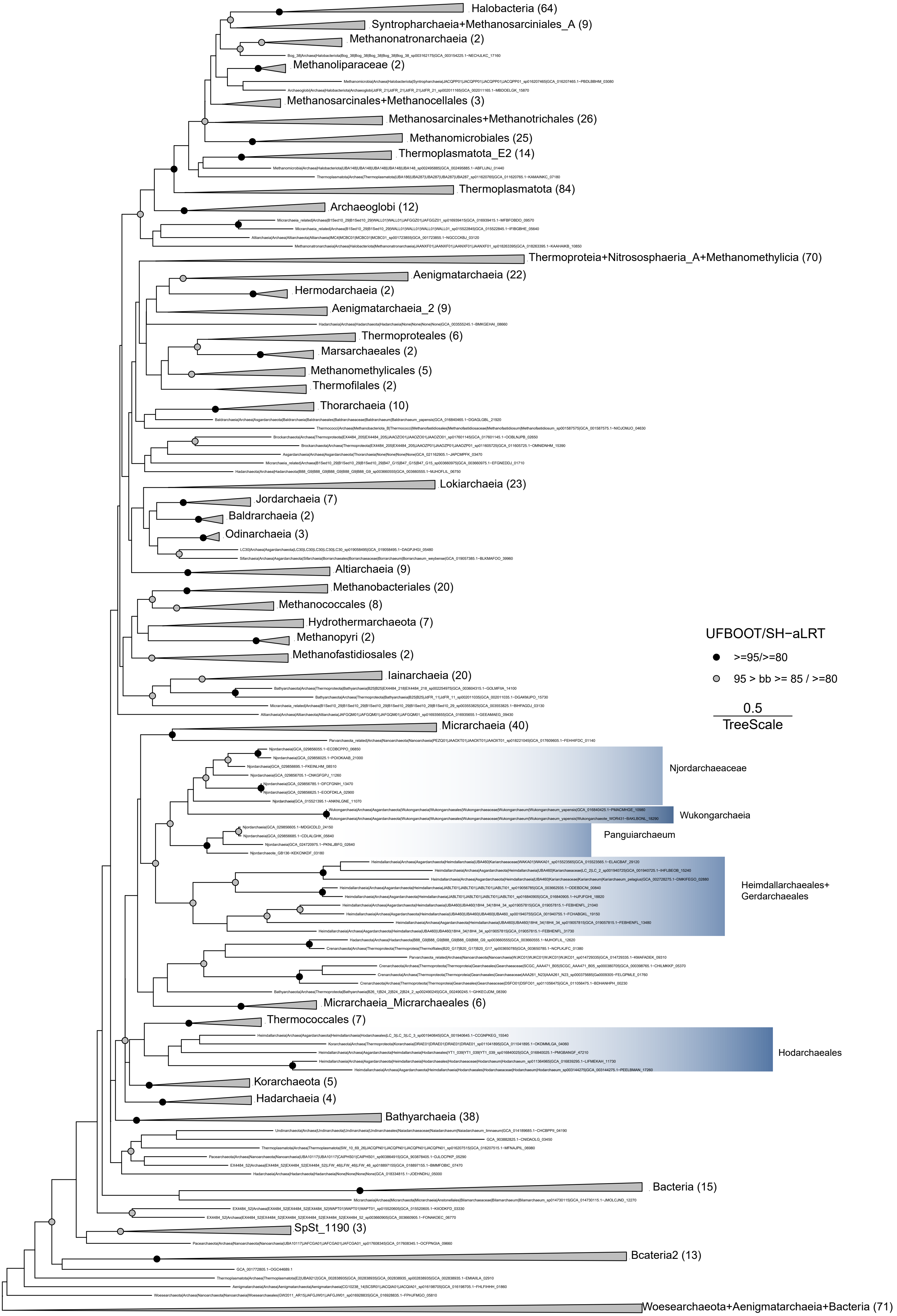


**Supplementary Figure 30. Maximum-likelihood phylogenetic analysis of CDP-archaeol synthase (CarS).** The alignment includes 746 sequences and 156 amino acid sites. The phylogeny is mid-point rooted. The analysis was performed using the LG+C60+G+F+PMSF model with a guide tree inferred using the LG+F+G model. Scale bar: Average substitution per site.

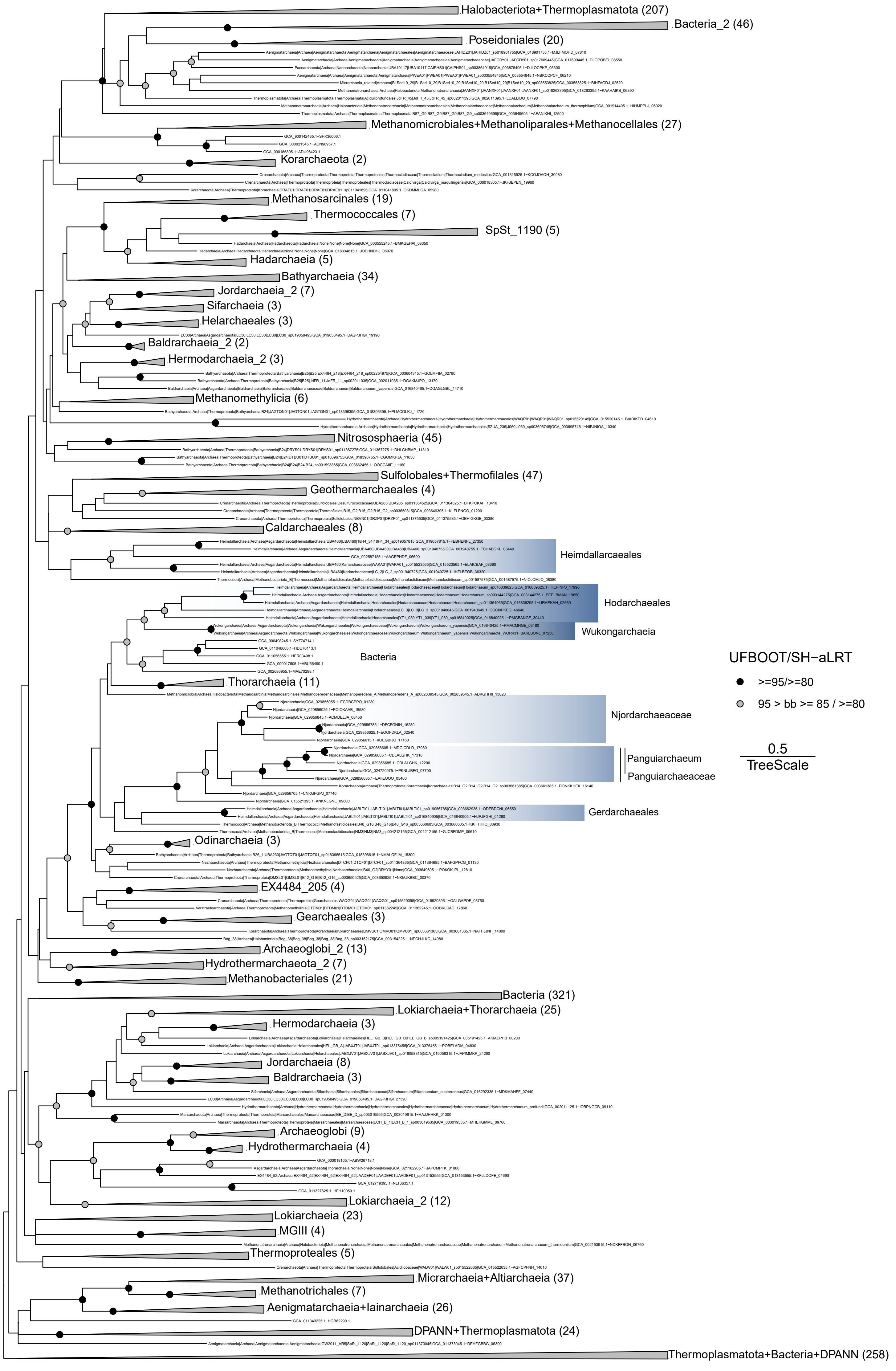


**Supplementary Figure 31. Maximum-likelihood phylogenetic analysis of phosphatidylglycerophosphate synthase (PgsA).** The alignment includes 1423 sequences and 184 amino acid sites. The phylogeny is mid-point rooted. The analysis was performed using the LG+C60+G+F+PMSF model with a guide tree inferred using the LG+F+G model. Scale bar: Average substitution per site.

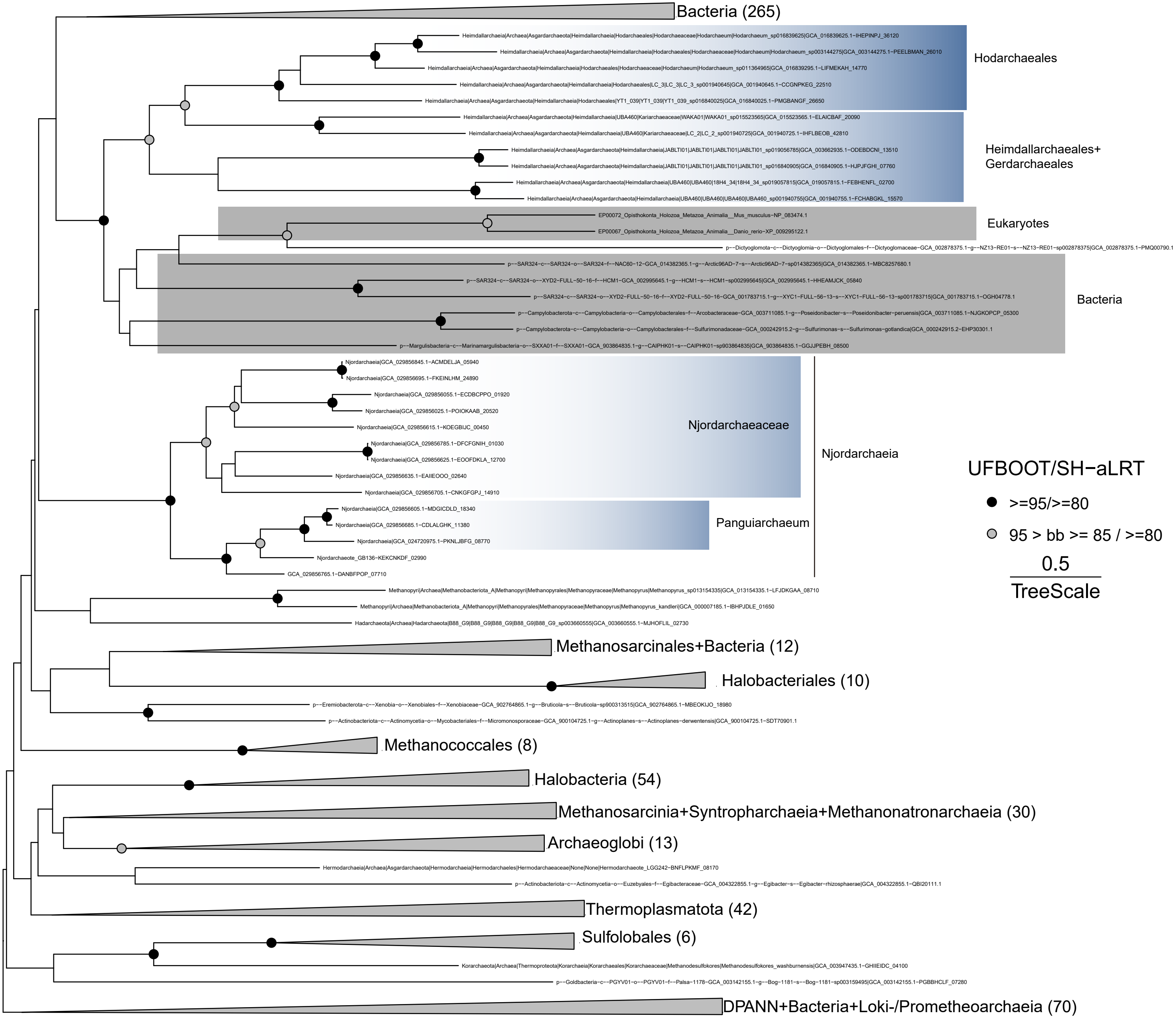


**Supplementary Figure 32. Maximum-likelihood phylogenetic analysis of phosphatidylglycerophosphate synthase (PssA).** The alignment includes 553 sequences and 211 amino acid sites. The phylogeny is mid-point rooted. The analysis used the LG+C60+G+F+PMSF model based on a guide tree inferred using LG+G. Scale bar: Average substitution per site.

A

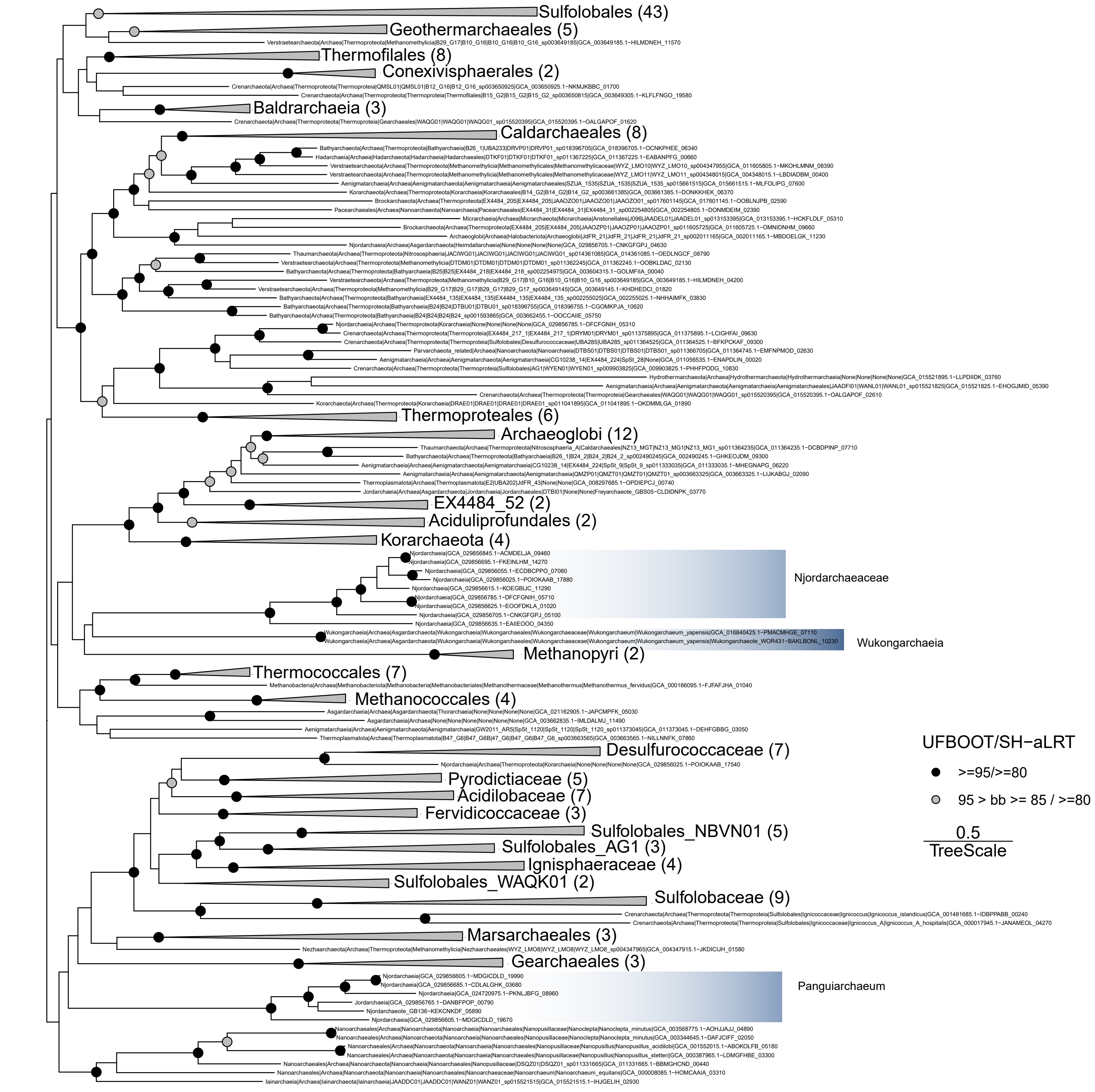


B

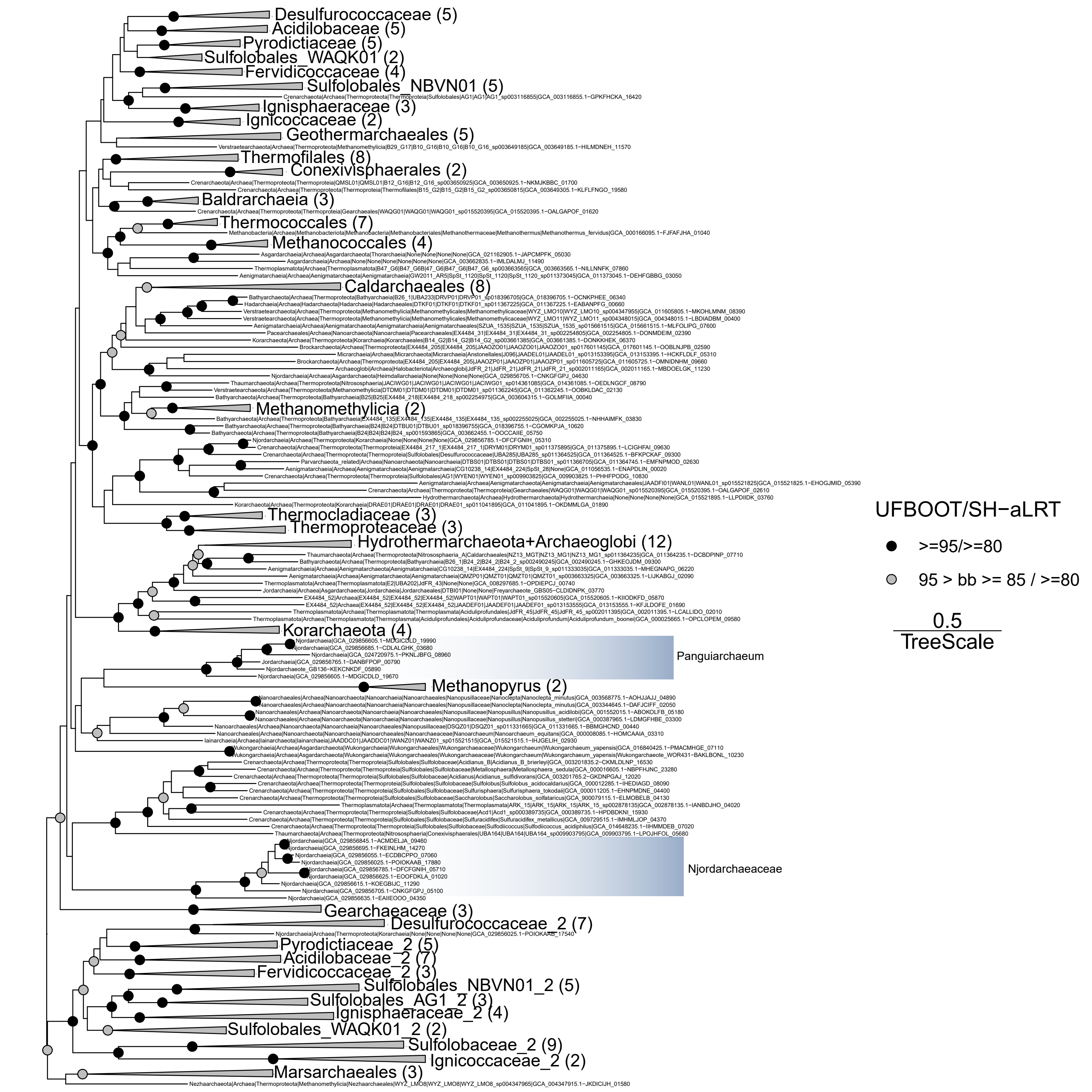


**Supplementary Figure 33. Phylogenetic analyses of reverse gyrase in archaea.** Homologs were identified using arCOG01526. Njordarchaeia sequences are highlighted in blue. A. Maximum-likelihood phylogeny of reverse gyrase. The analysis used the LG+C60+G+F+PMSF model with a guide tree based on LG +G+F model. The alignment contains 232 sequences with 849 amino acid sites. B. Maximum-likelihood phylogeny of reverse gyrase with 20% most compositionally-bi-ased site removed. The analysis used the LG+C60+G+F+PMSF model with a guide tree based on LG+G+F model. The alignment contains 232 sequences with 680 amino acid sites. Scale bar: Average substitution per site.

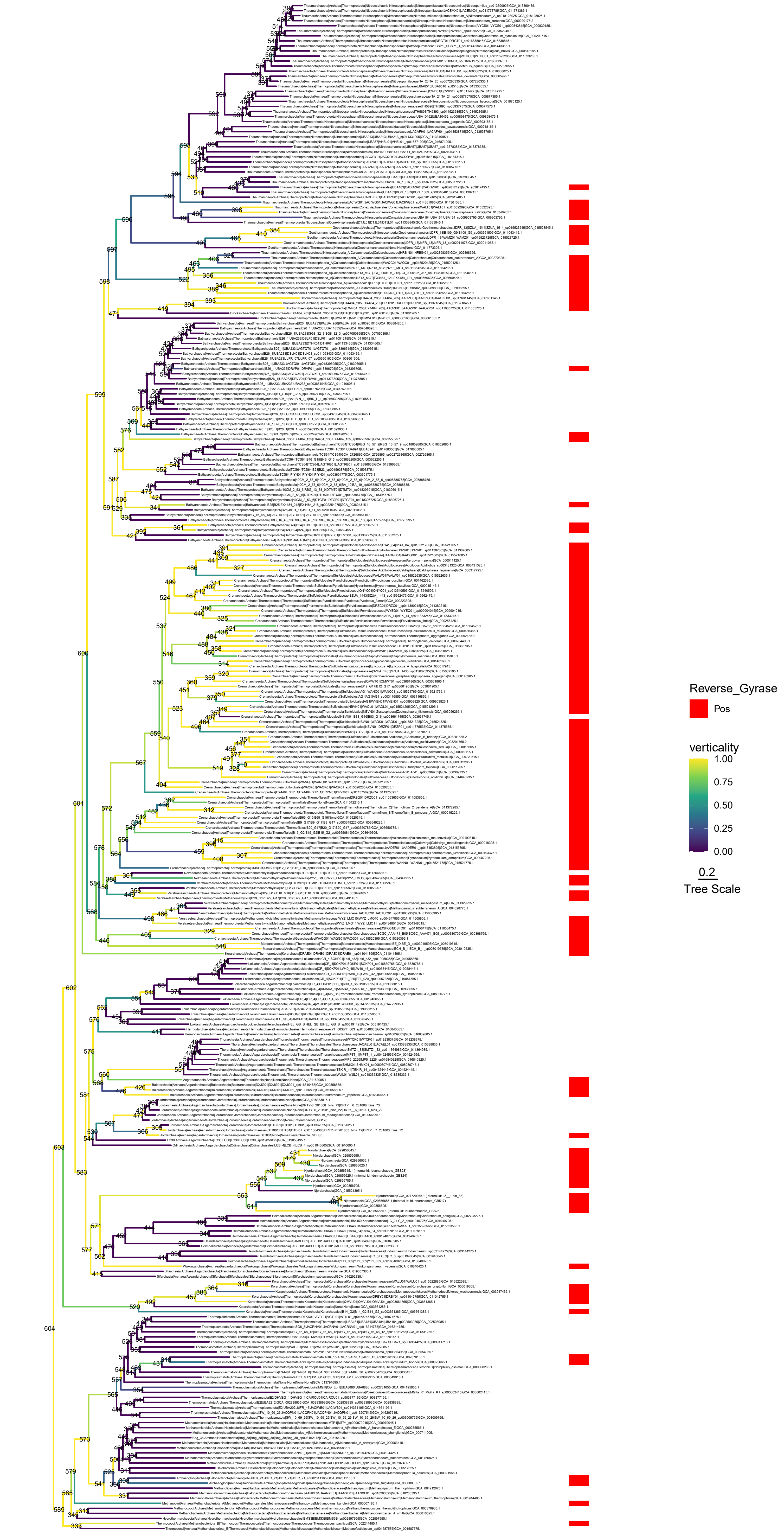


**Supplementary Figure 34. Verticality of reverse gyrase in the reconciliation analysis.** The species tree was inferred using a custom substitution mixture model, as in Figure 3A. Red tiles indicate the presence of Reverse gyrase (arCOG01526) in MAGs. Note that the node number is provided in the figures. Leaves were decorated with taxonomy based on Genome taxonomy database release 207. Scale bar: Average substitution per site.

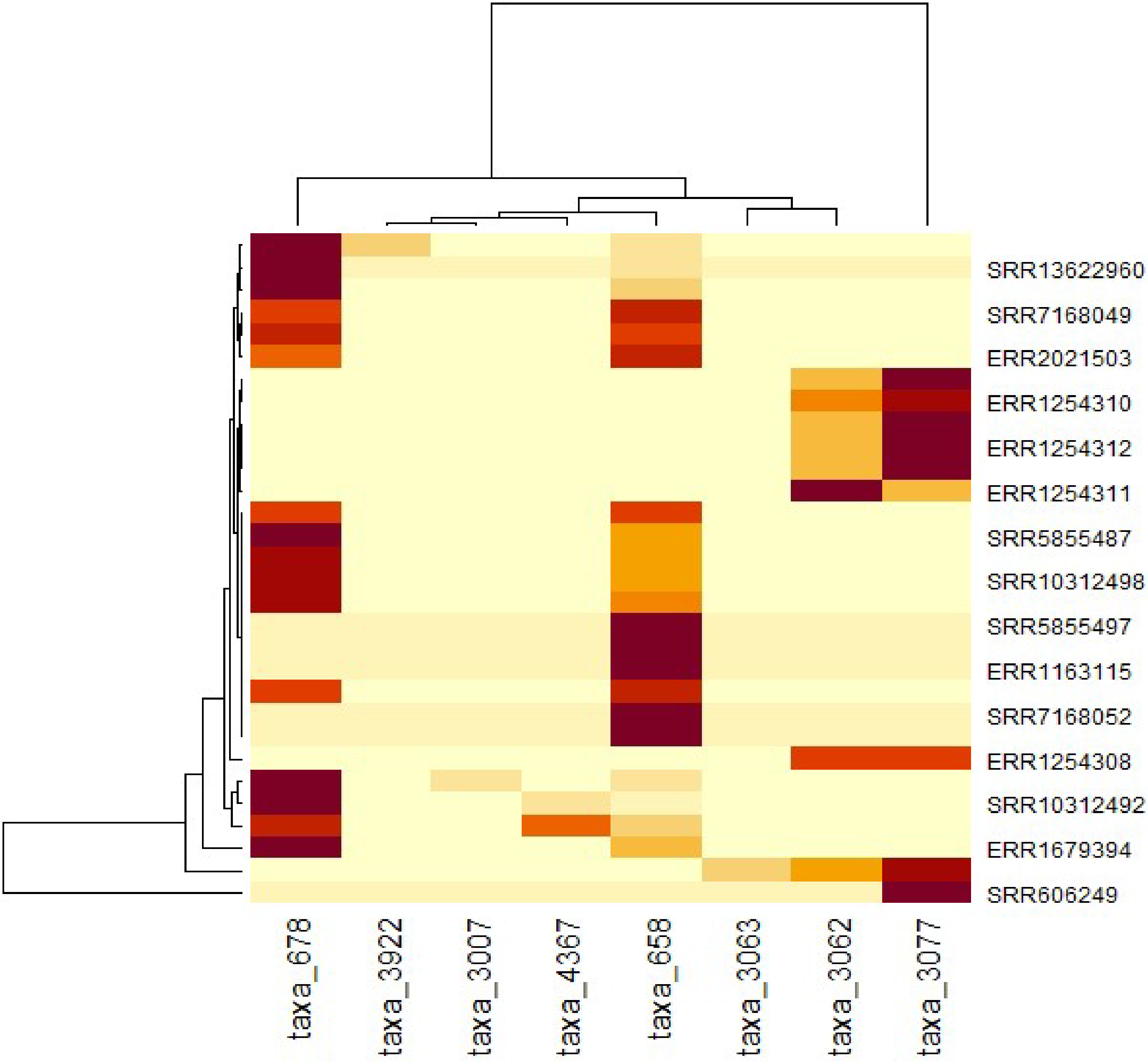


**Supplementary Figure 35.** Heatmap of Nanoarchaeum (taxa_658, taxa_3077) and Ignicoccus taxa (taxa_3007, taxa_678, taxa_3062, taxa_3063, taxa_3922, taxa_4367) across all samples in the dataset.

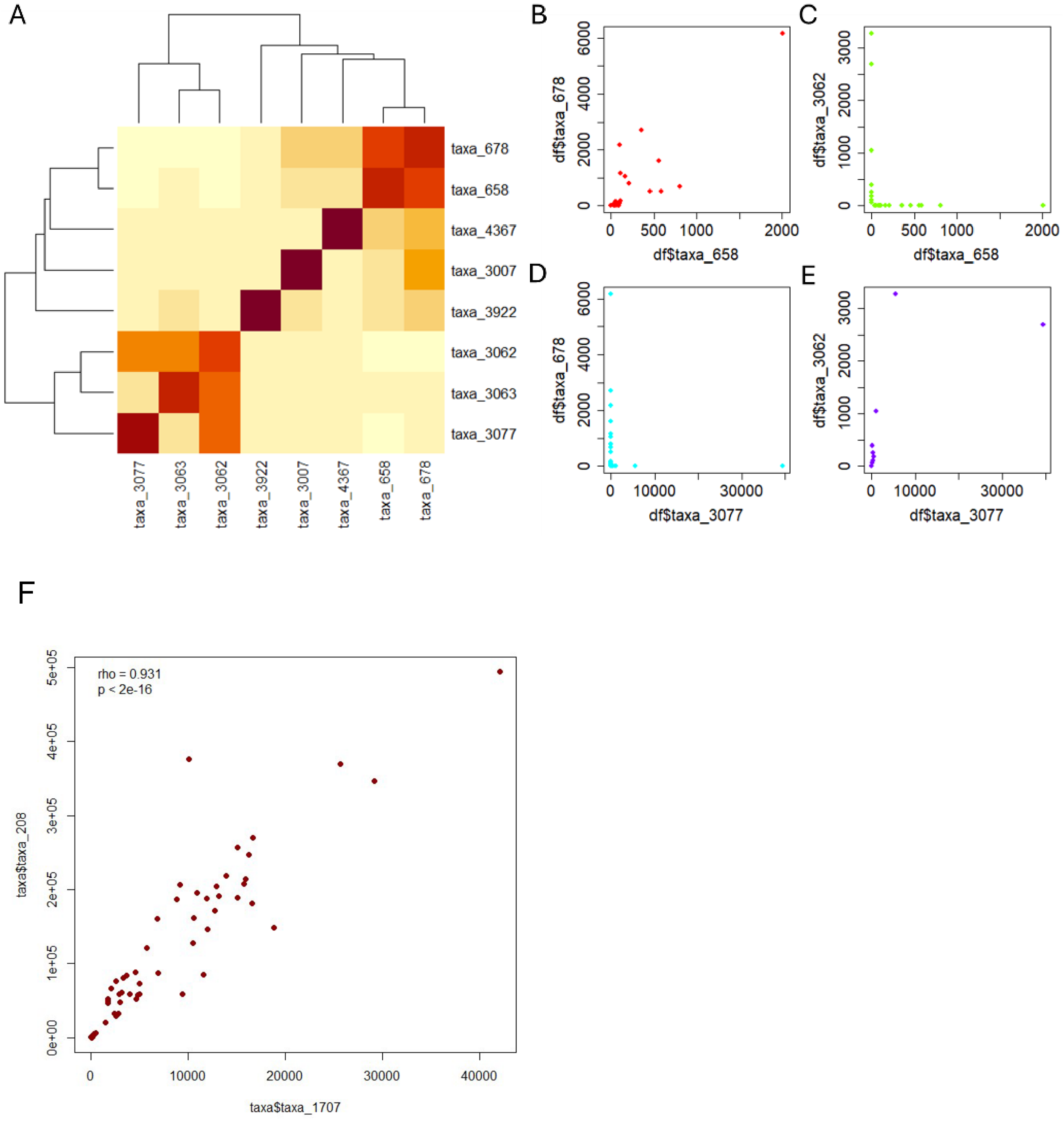


**Supplementary Figure 36.** Overview of Nanoarchaeum and Ignicoccus taxa and their relation across the dataset. (**A**) Correlation heatmap across all taxa annotated as either Nanoarchaeum or Ignicoccus. (**B-E**) Abundance plots of those pairs with significant Spearman correlation (p<0.05). **F.** Distribution of counts of Huberarchaeum (taxa_1707) and Altiarchaeum (taxa_208)
